# Interplay between store-operated calcium entry and mitochondrial phosphate handling modulates force and fatigue during exercise

**DOI:** 10.1101/2025.05.22.655415

**Authors:** Emmet A. Francis, Juliette Hamid, Anusha Kumar, Padmini Rangamani

**Affiliations:** Department of Pharmacology, University of California San Diego, La Jolla, CA, USA; Department of Mechanical and Aerospace Engineering, University of California San Diego, La Jolla, CA, USA

## Abstract

The dynamics of calcium ions (Ca^2+^) in skeletal muscles link electrochemical activation and contractile force generation. Recent experimental data suggest that store-operated Ca^2+^ entry (SOCE), the process of extracellular Ca^2+^ influx upon depletion of Ca^2+^ from the sarcoplasmic reticulum (SR), helps delay the onset of muscle fatigue during exercise. We hypothesize that SOCE regulates force generation during prolonged muscle activity by allowing for sustained Ca^2+^ release from the SR. We test this hypothesis with a quantitative biophysical model that simulates the biochemical events of muscle contraction, from depolarization at the T-tubules to Ca^2+^ release from the SR to Ca^2+^ binding and force generation throughout the myoplasm. We also consider the balance between Ca^2+^ removal from the myoplasm and SOCE through the T-tubule membrane, along with mitochondrial uptake of free Ca^2+^ and phosphate. We use the model to test the effects of SOCE inhibition on force production. The magnitude of myoplasmic Ca^2+^ and force are lower in SOCE knockout cells, especially when SOCE reduction is combined with impaired uptake of phosphate by mitochondria. We then test the effects of SOCE during resistance exercise or high-intensity interval training. These simulations predict a context-dependent relationship between force generation and SOCE – increased SOCE is associated with greater force production during resistance exercise, but worsens the effects of fatigue in certain cases of high-intensity training. Such SOCE-induced fatigue is attributed to phosphate accumulation in the myoplasm and can be mitigated by increased rates of mitochondrial phosphate uptake.

**Key points:**

- Store-operated calcium entry (SOCE) provides a mechanism for calcium ion (Ca^2+^) influx following depletion of Ca^2+^ from intracellular stores such as the sarcoplasmic reticulum (SR).
- Recent experiments suggest that SOCE is an important modulator of contractile force generation in skeletal muscle.
- Here, we develop a computational model of Ca^2+^ handling in the myoplasm, SR, and mitochondria and the resulting effects on force generation in skeletal muscle fibers to examine the role of SOCE during extended periods of activity.
- Our model predicts that increasing SOCE leads to enhanced force over periods of repeated stimuli during resistance exercise due to sustained Ca^2+^ release.
- Our simulations show a complex relationship between SOCE and force production during high-intensity interval training, with exacerbated phosphate accumulation in the myoplasm leading to force reduction for very high levels of SOCE. This effect can be mitigated by enhanced mitochondrial phosphate uptake.

**First author profile:** Emmet Francis is a K99/R00 awardee in the Rangamani Lab at UC San Diego whose research explores the intersection between cell signaling and mechanics. His doctoral research in the Heinrich Lab at UC Davis examined the role of calcium bursts in neutrophil chemotaxis and phagocytosis. More recently, he has used spatial modeling approaches to shed light on the role of nanoscale membrane curvature and nuclear deformation in YAP/TAZ mechanotransduction. In his own research lab, he plans to use both experiments and computational models to probe the mechanisms of bidirectional mechanotransduction in neutrophils.

**Abstract figure:** This study uses systems modeling to demonstrate a role for SOCE in sustained force generation during exercise. SOCE leads to two competing effects on contractile force in myofibers – increased crossbridge cycling due to elevated myoplasmic Ca^2+^ enhances force, whereas increased accumulation of myoplasmic phosphate (due to increased ATP hydrolysis) can lead to force reduction (fatigue). The tradeoff between these two effects is modulated by phosphate uptake into mitochondria via the phosphate carrier PiC. Figure created in BioRender.

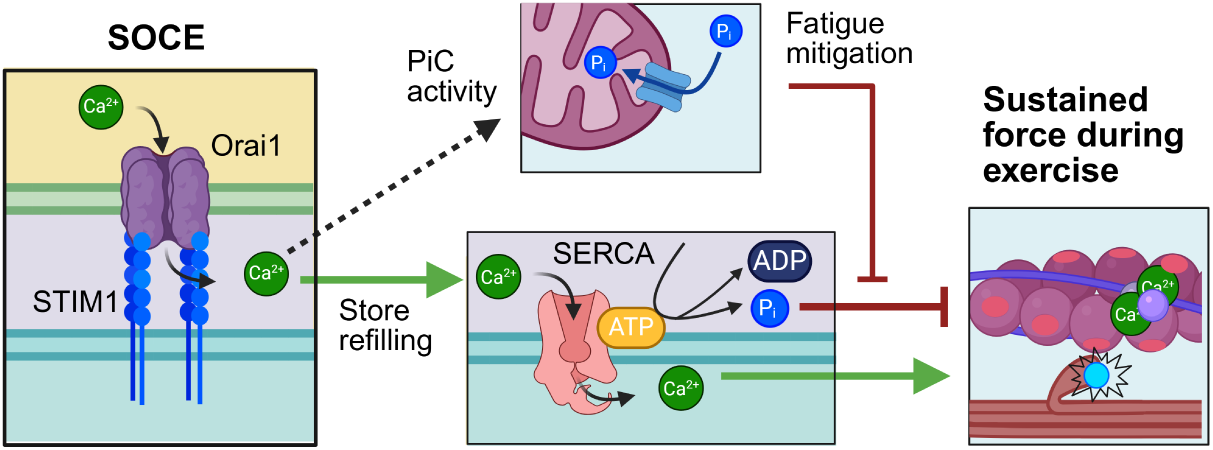

## Introduction

Skeletal muscle fibers (myofibers) are responsible for generating force to support locomotion of the animal body. They are positioned on the mesoscale of the muscle hierarchy, between the whole muscle composed of many bundles of fibers (fascicles) and subcellular myofibrils consisting of thick and thin filaments (Uchida and Delp, 2021; Mukund and Subramaniam, 2020). Contraction is initiated at the myofiber level where each fiber is stimulated by a motor neuron at the neuromuscular junction (NMJ). Understanding how myofibers integrate electrochemical inputs to initiate and sustain contraction is essential to dissect muscle performance in different contexts, from peak performance to rapid fatigue.

The evolution of contractile force in a myofiber is largely governed by the pattern of input stimulus received from its motor neuron. This stimulus is provided mainly through acetylcholine (ACh) released into the synaptic cleft, which binds to its receptor in the myofiber membrane (sarcolemma) allowing an influx of positively charged ions (Rodríguez Cruz et al., 2020). The frequency of ACh release events determines the extent of fiber contraction via rate encoding (Enoka and Duchateau, 2017; Fuglevand et al., 1993). At low frequencies, individual contractions (twitches) relax prior to the next stimulus, resulting in low force generation. In contrast, at higher stimulus frequencies, twitches fuse into a single tetanic contraction, producing much higher forces. This provides a mechanism for modulating muscle force during different activities. Consequently, different exercises are characterized by different patterns of stimulus at the NMJ. Force production itself is mediated by Ca^2+^ elevation in the myoplasm, which is initiated by voltage changes along specialized membrane invaginations known as transverse tubules (T-tubules). As voltage increases, voltage-gated Ca^2+^ channels are activated, triggering the opening of ryanodine receptors (RyRs) in the sarcoplasmic reticulum (SR) membrane to facilitate Ca^2+^ release from the SR. Myoplasmic Ca^2+^ then binds troponin on thin filaments, uncovering myosin binding sites to initiate cross-bridge cycling leading to myofiber contraction (Calderón et al., 2014).

The efficient contraction of muscle fibers relies on rapid transport of Ca^2+^ into and out of the myoplasm. Certain ATP-dependent membrane channels such as the plasma membrane Ca^2+^ ATPase (PMCA) in the sarcolemma and T-tubules and sarco/endoplasmic reticulum ATPase (SERCA) in the SR membrane pump Ca^2+^ back into the extracellular space or SR lumen, respectively. Additionally, excess free Ca^2+^ is absorbed by mitochondria via transport through the mitochondrial Ca^2+^ uniporter (MCU) (Dong and Tsai, 2023). Due to the constant removal of some Ca^2+^ from the cell through PMCA, SR Ca^2+^ levels are depleted over time in the absence of additional Ca^2+^ entry through the sarcolemma. The phenomenon of store-operated Ca^2+^ entry (SOCE) provides a compensates for this Ca^2+^ loss by coupling SR Ca^2+^ depletion to an additional influx through Ca^2+^ channels in the plasma membrane (Protasi et al., 2021; Hogan and Rao, 2015). SOCE relies on stromal interaction molecule 1 (STIM1) in the SR membrane, which acts as a Ca^2+^ sensor by binding SR Ca^2+^ through its luminal domain (Furukawa et al., 2014; Hoover and Lewis, 2011). Following Ca^2+^ dissociation from this binding site, STIM1 oligomerizes and binds to the Ca^2+^ channel Orai1 in the T-tubules, allowing Ca^2+^ influx into the myoplasm (Hoover and Lewis, 2011; Luik et al., 2008).

The importance of SOCE was originally appreciated in nonexcitable cells, but many recent studies have uncovered its importance in excitable cells such as neurons, cardiomyocytes, and myofibers (Basnayake et al., 2021; Wei-LaPierre et al., 2013; Boncompagni et al., 2017; Saftenku, 2022; Hermes et al., 2023; Koenig et al., 2019). For instance, during exercise in mice, STIM1 and Orai1 channels were found to colocalize at the SR-T-tubule junction, functioning as Ca^2+^ entry units (CEUs) (Wei-LaPierre et al., 2013; Boncompagni et al., 2017; Protasi et al., 2021). STIM-Orai assembly was shown to facilitate SOCE in the junctional region during each action potential, a phenomenon referred to as phasic SOCE (pSOCE) (Lilliu et al., 2025, 2020; Koenig et al., 2018, 2019; Lilliu et al., 2021). CEUs have been proposed to facilitate rapid recovery of Ca^2+^ within the fiber and delay the onset of fatigue (Boncompagni et al., 2017; Wei-LaPierre et al., 2013). Indeed, muscles from dominant-negative Orai1 mice (dnOrai1) were more susceptible to fatigue and produced lower contractile force ex vivo (Wei-LaPierre et al., 2013). However, other recent data suggests that SOCE can worsen fatigue in some cases; phosphorylation of STIM1 by AMP-activated protein kinase (AMPK) was found to both decrease SOCE activity and prevent fatigue (Nelson et al., 2020).

Understanding the role of SOCE in muscle fatigue requires a comprehensive understanding of the molecular mechanisms underlying fatigue in single muscle fibers, including rapid accumulation of phosphate in the myoplasm, reduced Ca^2+^ release from the SR, and reduced myofibrillar Ca^2+^ sensitivity (Shorten et al., 2007; Cheng et al., 2018; Allen and Trajanovska, 2012; Hendry et al., 2025). On the one hand, SOCE allows for increased store refilling and might be expected to alleviate fatigue. However, high levels of SOCE can also lead to Ca^2+^ overload, causing accumulation of phosphate in the myoplasm due to increased rates of cross bridge cycling and increased activity of SERCA and PMCA. Such phosphate accumulation causes a reduction in contractile force due to enhanced reversal of the power stroke (Allen and Trajanovska, 2012; Allen and Westerblad, 2001). Removal of phosphate from the myoplasm relies, in part, on transport into the SR (Shorten et al., 2007) and into the mitochondrial matrix through the mitochondrial phosphate carrier (PiC) (Seifert et al., 2015; Dong and Tsai, 2023). It is crucial to understand how the dynamics of phosphate uptake by mitochondria affects fatigue onset in cells with elevated SOCE.

Here, we use a computational model to predict when SOCE enhances force production and when it accelerates the onset of fatigue in myofibers. We first fit our model to experimental measurements of myoplasmic Ca^2+^ and sarcolemma voltage in the literature, and then show that this model recapitulates measurements of Ca^2+^ and force in SOCE-deficient myofibers. We furthermore consider how the effects of SOCE deficiency in cells depend on phosphate uptake by mitochondria. Finally, we predict the role of SOCE for different patterns of stimuli chosen to mimic high-intensity interval training and resistance exercise. Our findings reconcile apparent contradictions in the literature, showing that the role of SOCE is context-dependent – force production is generally enhanced by SOCE at lower stimulus frequencies, but can worsen fatigue during extended, high-frequency activation due to phosphate accumulation in the myoplasm. This effect is tuned by the rates of phosphate uptake by mitochondria in the myoplasm.

## Results

### Model Development

To investigate the impact of SOCE on Ca^2+^ dynamics in skeletal muscle, we developed a well-mixed multi-compartment model describing action potential generation, Ca^2+^ handling, phosphate handling and force generation in fast-twitch myofibers. We included junctional and bulk regions of the myoplasm, SR, mitochondria, and extracellular space (Figure 1A). Junctional regions correspond to the volumes and surfaces at the triad, which consists of a T-tubule surrounded by two sections of the SR on either side (terminal cisternae). We did not consider propagation of electrochemical signals along the length of the fiber, instead choosing to focus on a small representative region of the myofiber over which molecules remain relatively well-mixed. Furthermore, we assumed that the action potential propagates into the T-tubules much faster than other events in our model. Throughout, we differentiate between each subcompartment using subscripts for clarity; for instance, Ca^2+^ concentration in the junctional myoplasm is written as [Ca^2+^]_myo,junc_ (all species are included in Table A4). Species in the myoplasm and SR are allowed to diffuse between the bulk and junctional regions, whereas mitochondrial species are confined to either junctional or bulk regions.

**Figure 1:**
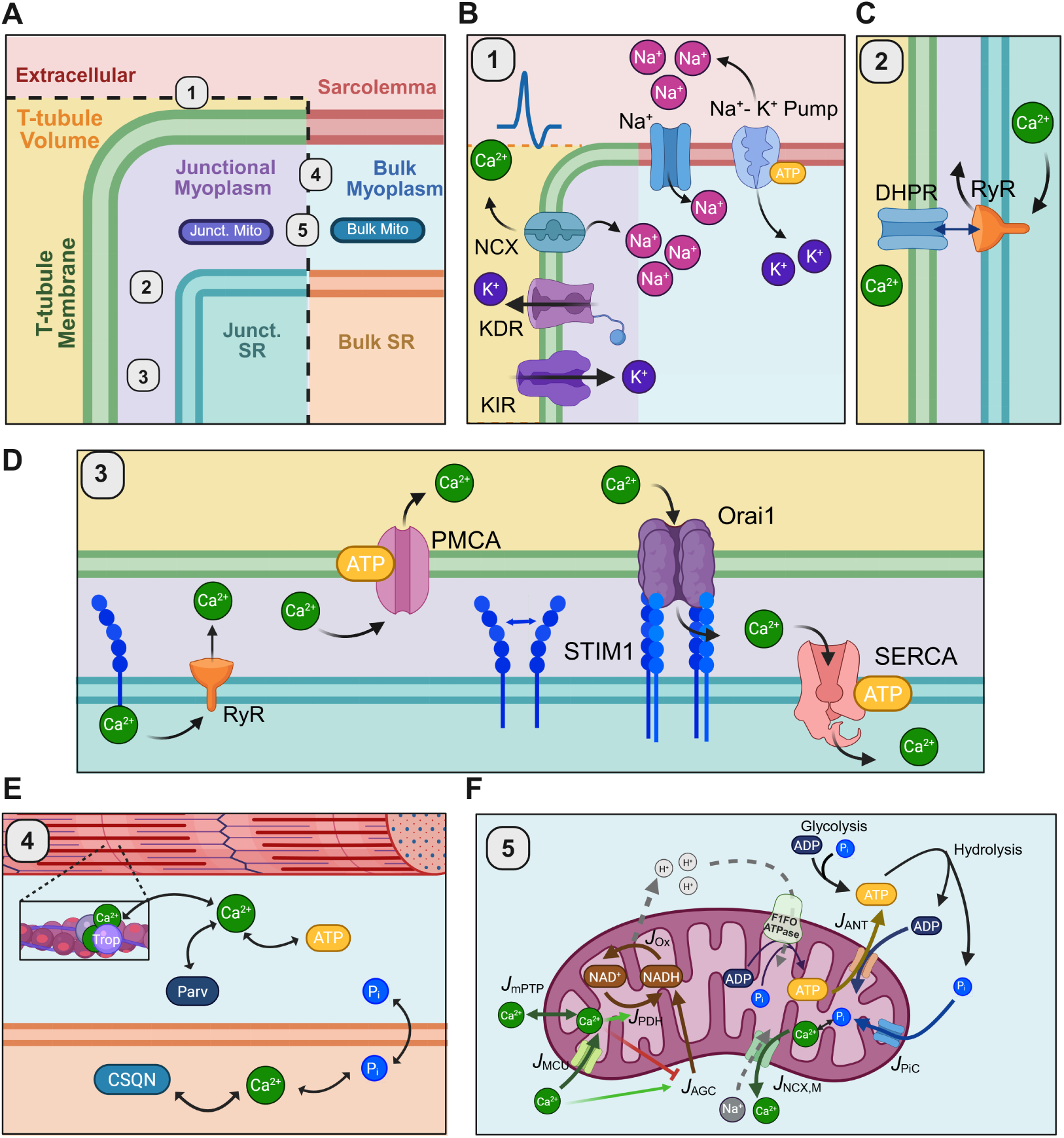
Schematic for model components of myofiber Ca^2+^ signaling. A) A subregion of the cross-section of the muscle fiber is divided into compartments, each labeled with different colors. Ions freely diffuse between bulk and junctional regions of each non-mitochondrial compartment, separated by black dashed lines here. B-D) The sequence of signaling events directly correlated to force generation are grouped into five modules. In module 1, the applied current triggers an action potential through Na^+^ and K^+^ fluxes (B). In module 2, action potential propagation leads to DHPR-RyR allosteric interactions, triggering Ca^2+^ release from the SR (C). In module 3, Ca^2+^ unbinds from STIM1 during SR depletion, causing STIM1 to oligomerize and form complexes with Orai1 and allowing Ca^2+^ influx from the extracellular space into the myoplasm (D). In module 4, Ca^2+^ binds to proteins in the SR and myoplasm and induces the cross-bridge cycle, producing contractile force (E). In module 5, Ca^2+^ and phosphate are taken up by the mitochondria, leading to enhanced ATP production via oxidative phosphorylation (F). Figure created in BioRender.

Our model is divided into 5 modules, including action potential generation (Module 1, Figure 1B), Ca^2+^ influx and release (Module 2, Figure 1C), Ca^2+^ removal and SOCE (Module 3, Figure 1D), Ca^2+^ buffering and cross-bridge cycling (Module 4, Figure 1E), and Ca^2+^ and phosphate handling by mitochondria (Module 5, Figure 1F). The complete model consists of a system of 65 ordinary differential equations (ODEs), including flux balance equations and equations governing changes in receptor or channel state. We briefly summarize each module below, with a full description of the model given in Appendix A (Tables A3 to A7).

#### Module 1: Action potential generation

Action potentials were modeled using a Hodgkin-Huxley-type formulation, as in previous models developed for mammalian skeletal muscle (Cannon et al., 1993; Wallinga et al., 1999; Shorten et al., 2007; Senneff and Lowery, 2021) (Figure 1B). We considered a Na^+^ current through a voltage-gated Na^+^ channel (*I*_Na_+), followed by K^+^ efflux through the K^+^ delayed rectifier (*I*_KDR_), governed by equations from (Wallinga et al., 1999). We also included currents through the K^+^ inward rectifier (*I*_KIR_), which helps to maintain the resting potential of the membrane (Standen and Stanfield, 1978), and the Na^+^-K^+^ pump (*I*_Na_+_-K_+), which restores gradients in Na^+^ and K^+^ across the sarcolemma (Wallinga et al., 1999). Other secondary currents regulate the resting membrane potential, including Na^+^ import through the Na^+^-Ca^2+^ exchanger (*I*_NCX_) and a Cl^−^ current (*I*_Cl_*−*). Altogether, the total ionic currents through the T-tubule membrane (subscript TT) and bulk sarcolemma (subscript SL) are given by

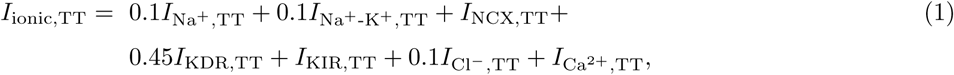

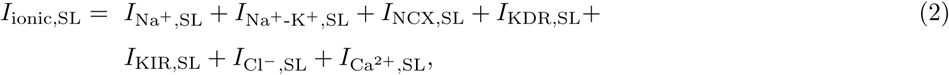

where *I*_Ca^2+^_,_TT_ and *I*_Ca^2+^_,_SL_ are additional Ca^2+^ currents defined in Module 3. Full expressions for each current are given in Table A5 and their relative contribution at T-tubules vs. sarcolemma are summarized in Table A3. Balancing these ionic currents with a capacitive current at the sarcolemma and accounting for conduction between the sarcolemma and T-tubules yields

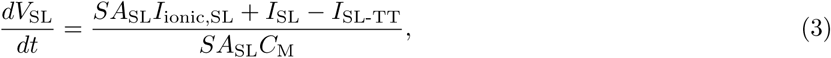

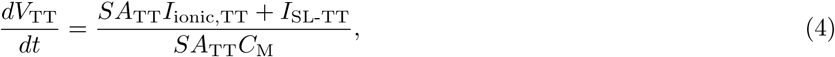

where *SA*_SL_ is the surface area of the sarcolemma, *SA*_TT_ is the surface area of the T-tubules, *C*_M_ is the specific membrane capacitance, and *I*_SL_ is the stimulus current through the sarcolemma. This stimulus was modeled as square waves with a pulse width of 1 ms. *I*_SL-TT_ is the current from the sarcolemma into the T-tubule membrane, calculated as in (Wallinga et al., 1999):

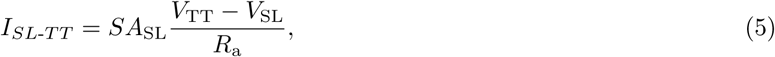

where *R*_a_ is the access resistance at the entrance to the T-tubules.

#### Module 2: Ca^2+^ release

Ca^2+^ release from the SR occurs in the junctional region upon a sufficient increase in T-tubule membrane voltage. The L-type Ca^2+^ channel, the dihydropyridine receptor (DHPR), acts as a voltage sensor in the T-tubule membrane, allosterically interacting with the ryanodine receptor (RyR) in the SR membrane to facilitate its opening (Ríos et al., 1993). In line with previous literature, we assumed that the Ca^2+^ flux through DHPR itself is negligible and it mainly serves as a voltage sensor to gate the RyR (Dayal et al., 2017).

The resulting flux through the RyR is given by:

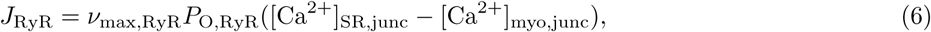

where *ν*_max,RyR_ is the maximum permeability of the RyR and *P*_O,RyR_ is the open probability of the RyR governed by Ca^2+^ concentration and T-tubule membrane voltage. The voltage dependence of *P*_O,RyR_ is dictated by a 10-state Monod, Wyman, and Changeux (MWC)-type model (as in (Ríos et al., 1993)) and the Ca^2+^ dependence captures slow deactivation, as introduced by Senneff and Lowery (Senneff and Lowery, 2021) (see expressions in Table A5).

#### Module 3: Ca^2+^ removal and SOCE

Ca^2+^ homeostasis relies on rapid removal from the myoplasm via ATP-dependent fluxes through PMCA into the extracellular space and through SERCA into the SR lumen. We represented these fluxes phenomenologically using first-order Hill equations that depend on both Ca^2+^ concentration and ATP concentration (Table A5). Ca^2+^ is also passively transported from the myoplasm to the extracellular space through NCX, as dictated by *J*_NCX_ (proportional to *I*_NCX_ in Equations (1) and (2)) and into mitochondria via the mitochondria Ca^2+^ uniporter (MCU), described in Module 5 below.

Given that not all Ca^2+^ released from the SR is pumped back in through SERCA, SOCE is required to restore SR Ca^2+^ levels following rapid depletion. We drew from previous work across several cell types (Saftenku, 2022; Dolan and Diamond, 2014; Hoover and Lewis, 2011; Rincón et al., 2021) to mathematically describe SOCE in skeletal muscle. The SR luminal domain of STIM1 binds SR Ca^2+^, allowing it to act as a Ca^2+^ sensor (Furukawa et al., 2014). Upon depletion of SR Ca^2+^, Ca^2+^ unbinding leads to STIM1 oligomerization and binding to tetrameric Orai1 channels at SR-T-tubule junctions (Hoover and Lewis, 2011), leading to Ca^2+^ influx through Orai1. Previous experimental measurements (Luik et al., 2008; Furukawa et al., 2014) demonstrate that the relationship between Orai1 conductance and SR Ca^2+^ is well-represented by a Hill equation with a coefficient close to 4, as utilized by multiple previous models (Saftenku, 2022; Koenig et al., 2019; Rincón et al., 2021; Mansour et al., 2026). Accordingly, we modeled the steady-state open probability of Orai1 channels, *P*_O,Orai1,_*_∞_*, as:

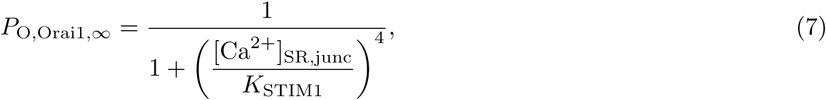

where *K*_STIM1_ is an equilibrium constant dictating the junctional SR Ca^2+^ concentration at which SOCE is at half maximum activation. We expected the system to approach steady state over some characteristic time *τ*_SOCE_ required for STIM1 oligimerization, diffusion, and binding with Orai1 to form CEUs (Hoover and Lewis, 2011):

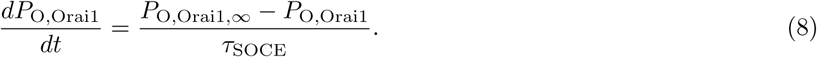

This approach allowed us to represent SOCE without introducing a large number of additional free parameters. The Ca^2+^ flux due to SOCE was localized to the T-tubules and was given by

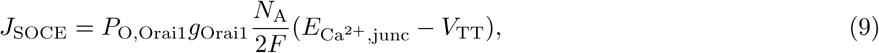

where *g*_Orai1_ is the Orai1 conductance, *N*_A_ is Avogadro’s number, and *F* is Faraday’s constant. *E*_Ca_2+,_junc_ is the Nernst potential for Ca^2+^ across the T-tubule membrane:

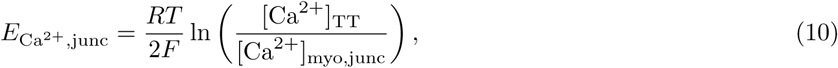

where *R* is the gas constant and *T* is the temperature.

#### Module 4: Ca^2+^ binding and crossbridge cycling

Many proteins in the myoplasm and SR act as Ca^2+^ buffers, such that the total Ca^2+^ concentration in either compartment is much higher than the free Ca^2+^ concentration (Butera et al., 2021; Murphy et al., 2009; Nogueira et al., 2022). In the myoplasm, we considered Ca^2+^ binding to parvalbumin, ATP, and troponin, in addition to competitive binding of Mg^2+^ to parvalbumin and ATP (Baylor and Hollingworth, 2012, 2007; Shorten et al., 2007; Senneff and Lowery, 2021). In the SR lumen, we accounted for Ca^2+^ binding to calsequestrin (Cannell and Allen, et al., 2007).

Ca^2+^ binding to troponin on the thin filament in the bulk myoplasm leads to cross-bridge cycling. In short, two Ca^2+^ ions bind sequentially to a single troponin complex, causing tropomyosin to change shape and expose myosin binding sites on the thin filament (Shorten et al., 2007; Uchida and Delp, 2021; Calderón et al., 2014). We modeled this using a six-state cross-bridge model, modified from the eight-state model in Shorten et al (Shorten et al., 2007) as detailed in Tables A5 and A6 and depicted in Figure A1. The production of contractile force was assumed to be proporational to the density of cross-bridges in a post-power stroke (force-generating) conformation *A*_Post_:

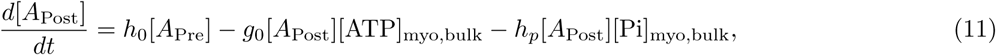

where [*A*_Pre_] is the density of pre-power stroke cross bridges, *h*_0_ is the power stroke rate, *g*_0_ is the rate of myosin head detachment post-power stroke, and *h_p_* is the reverse rate to the pre-power stroke conformation. In the above relationship, we explicitly assumed that reactions depended on concentrations of ATP ([ATP]_myo,bulk_) and inorganic phosphate ([Pi]_myo,bulk_) according to mass action, an effect neglected in some previous models (Shorten et al., 2007; Senneff and Lowery, 2021). This is an important consideration, as the dependence of power stroke reversal on phosphate has been suggested as a mechanism for fatigue (Allen and Trajanovska, 2012).

#### Module 5: Adenine nucleotide, phosphate, and Ca^2+^ handling by mitochondria

In the final module, we include the effects of mitochondrial phosphate uptake, Ca^2+^ uptake, and ATP generation via oxidative phosphorylation. ATP hydrolysis due to cross-bridge cycling and activity of ion pumps and other intracellular processes in the above modules was explicitly tracked in the myoplasm. In line with the dominant role for glycolytic fluxes in fast-twitch muscle fibers (Murgia et al., 2021; Schiaffino and Reggiani, 2011), we assumed that ATP was generated within the myoplasm via glycolysis to restore ATP concentrations to a setpoint during simulations of extended exercise. This mathematical description of glycolysis was not meant to capture the full complexity of glycolytic flux regulation in skeletal muscle, but to implicitly the established dependence of glyolysis rates on muscle activation (Conley et al., 1998) and concentrations of phosphate, ATP, and ADP (Crowther et al., 2002).

We adopted models of Ca^2+^ handling and ATP generation in mitochondria from several previous studies (Wacquier et al., 2016; Bertram et al., 2006; Leung et al., 2021), assuming that ATP generation is positively regulated by mitochondrial Ca^2+^ uptake due to enhanced production of NADH required for oxidative phosphorylation (Figure 1F). We accounted for several Ca^2+^ fluxes across the mitochondrial membranes due to the mitochondrial Ca^2+^ uniporter (MCU), mitochondrial permeability transition pore (mPTP), and mitochondrial NCX (Figure 1F). In addition, phosphate is transported into the mitochondria via the mitochondrial phosphate carrier (PiC) (Seifert et al., 2015). We simulated this flux using a reversible Hill equation:

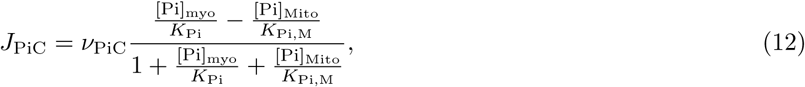

where [Pi]_myo_ and [Pi]_Mito_ are the free phosphate concentrations in the myoplasm and in the mitochondrial matrix, *K*_Pi_ and *K*_Pi,M_ are the associated Hill constants, and *ν*_PiC_ is the maximal phosphate transport rate through PiC. This expression carries the same dependence of PiC on mitochondrial and extra-mitochondrial phosphate as in previous models (Bazil et al., 2010; Nguyen et al., 2007). We also accounted for retention of Ca^2+^ in mitochondria due to Ca^2+^-phosphate formation (Wei et al., 2015; Seifert et al., 2015), which increases the mitochondrial capacity for

#### Assembly of the full model

The fluxes outlined in Modules 2-5 were assembled into systems of ODEs for Ca^2+^ in each region of the myoplasm, SR, and mitochondria. Furthermore, we computed the additional Ca^2+^ currents through the T-tubule membrane and bulk sarcolemma introduced in Equations (3) and (4):

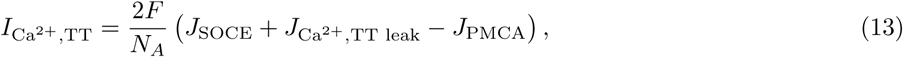

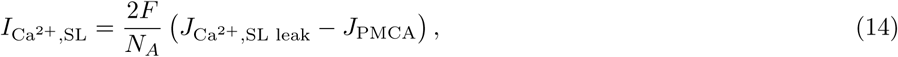

where *J*_Ca_2+,_TT leak_ and *J*_Ca_^2+^,_SL leak_ are leak fluxes through the T-tubule membrane and sarcolemma, respectively.

Most species in the model were assumed to exist in both junctional and bulk regions, with the exception of Orai1 (T-tubules only) and all cross-bridge related variables (bulk myoplasm only). Furthermore, concentrations of ions were tracked within the T-tubule volume, but bulk extracellular ion concentrations were assumed to be constant.

### Calibrated model recapitulates dynamics of membrane potential and myoplasmic Ca^2+^

Our model utilizes a collection of channels and receptors not combined in previous models. Accordingly, while we used initial parameter estimates from the literature, it was necessary to formally estimate best-fit values for each parameter. This allowed us to reliably consider the importance of SOCE relative to other Ca^2+^ fluxes throughout the myofiber. Excluding geometric parameters, there were 142 free parameters (Table A7) in the model. 40 of these parameters were specific to mitochondria (Module 5) and were calibrated independently to data from (Wei et al., 2015) as described in Methods and shown in Figure A3. After fixing these parameters, we conducted a global Morris sensitivity analysis to identify the most influential parameters in the model (Morris, 1991) (Figure 2A-B,Figure A2). As quantities of interest (QOIs), we considered the maximum and time-averaged values of the myoplasmic Ca^2+^ concentration and sarcolemma voltage over a single excitation event. Myoplasmic Ca^2+^ was defined as a weighted average by volume:

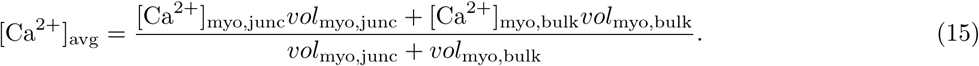

**Figure 2:**
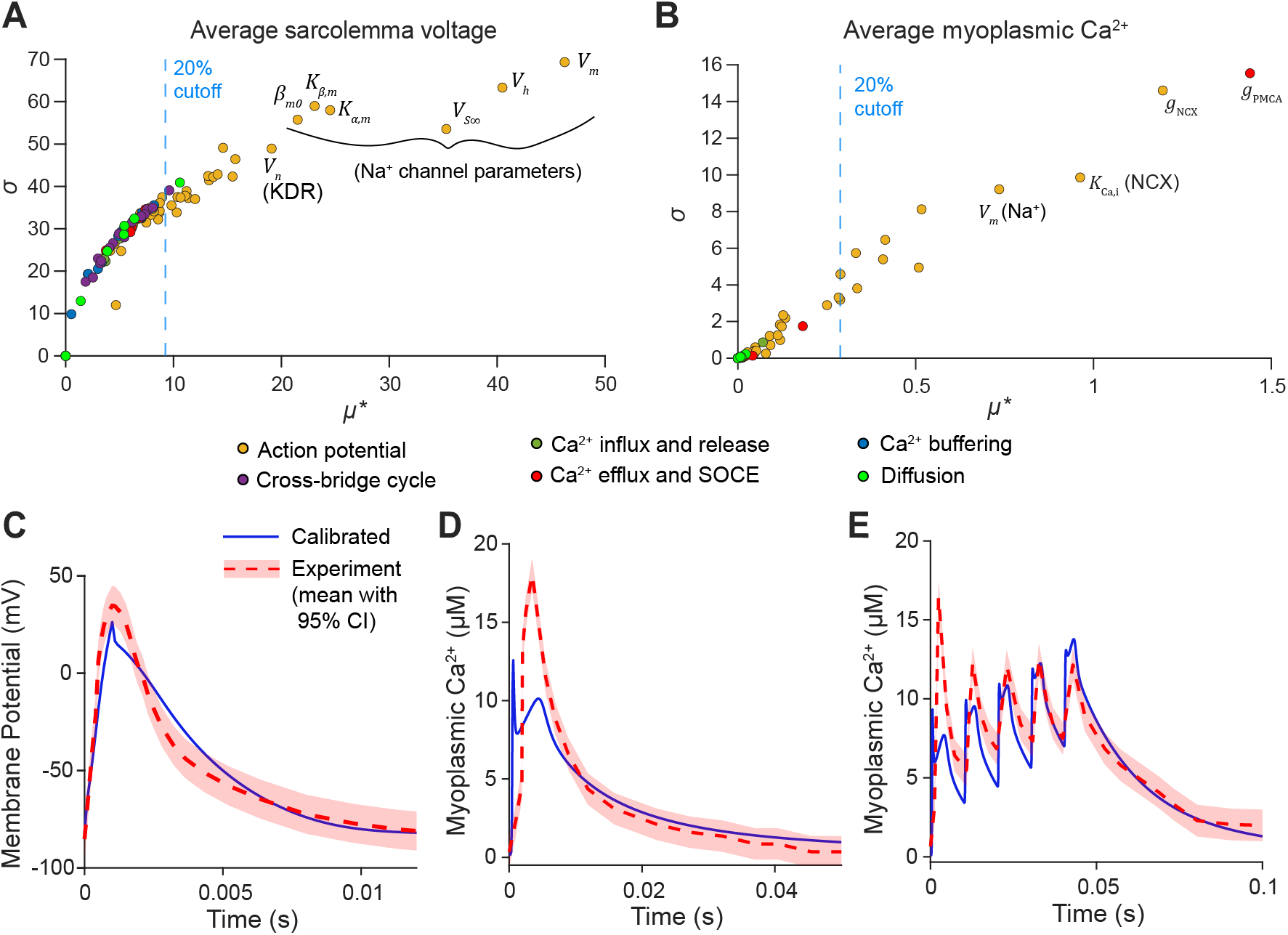
Sensitivity analysis and model calibration. A, B) Results from Morris sensitivity analysis of average sarcolemma voltage (A) and average [Ca^2+^]_myo_ (B) as quantities of interest (QOIs). The horizontal axis displays the absolute value of the mean, *µ^∗^*, and the vertical axis shows the standard deviation, *σ*, of elementary effects. The parameters with the highest values of *µ^∗^* are labeled individually. C) Results of model calibrated to experimental membrane potential measurements (Miranda et al., 2020) (C) and to [Ca^2+^]_myo_ dynamics data for a single pulse stimulus (D) (Baylor and Hollingworth, 2003) or a train of five stimulus pulses at 100 Hz (E) (Rincón et al., 2021). All experimental data were extracted from source papers using PlotDigitizer; specifically, from Fig 2B (WT curve) in (Miranda et al., 2020), from Fig 2A (fast twitch curve) in (Baylor and Hollingworth, 2003), and from Fig 1B (IIB curve) in (Rincón et al., 2021). The minimized weighted sum of square errors (defined in Equations (21) to (23)) was 32.375. Conditions were varied according to extracellular buffer and temperature in each experiment. (Miranda et al., 2020): [Na^+^]_EC_ = 1.18 × 10^5^ µM, [K^+^]_EC_ = 5.33 × 10^3^ µM, [Cl*^−^*]_EC_ = 1.26 × 10^5^ µM, [Ca^2+^]_EC_ = 1.80 × 10^3^ µM, *T* = 295.15 K; (Baylor and Hollingworth, 2003): [Na^+^]_EC_ = 1.50 × 10^5^ µM, [K^+^]_EC_ = 2.00 × 10^3^ µM, [Cl*^−^*]_EC_ = 1.58 × 10^5^ µM, [Ca^2+^]_EC_ = 2.00 × 10^3^ µM, *T* = 293.15 K; (Rincón et al., 2021): [Na^+^]_EC_ = 1.38 × 10^5^ µM, [K^+^]_EC_ = 3.90 × 10^3^ µM, [Cl*^−^*]_EC_ = 1.44 × 10^5^ µM, [Ca^2+^]_EC_ = 1.00 × 10^3^ µM, *T* = 295.15 K.

Average sarcolemma voltage was most sensitive to parameters associated with the Na^+^ channel (*V_m_*, *V_h_*, *V_S∞_*, *V_αm_*, and *V_βm_* ; defined in Table A7) and with the KDR channel (*V_n_*), in agreement with previous work (Shorten et al., 2007; Wallinga et al., 1999) (Figure 2A). Ca^2+^ dynamics were most sensitive to Ca^2+^ export through PMCA (*g*_PMCA_) and efflux through the Na^+^-Ca^2+^ exchanger (*g*_NCX_ and *K*_Ca,i_), validating the central importance of Ca^2+^ efflux mechanisms (Figure 2B).

We proceeded to estimate parameters in two additional stages, as fully described in Methods. We first fit the model to measurements of membrane potential (Miranda et al., 2020), only varying the parameters with greater than 20 % relative sensitivity with respect to average and/or peak voltage (Figure A2A-B). Then, starting from these parameter values as new initial estimates and allowing all parameters to vary, we simultaneously fit our model to voltage and to myoplasmic Ca^2+^ data collected from multiple papers that conducted fluorescence or voltage measurements in single mouse fast-twitch myofibers (Miranda et al., 2020; Baylor and Hollingworth, 2003; Rincón et al., 2021). In general, the fitting process resulted in a strong fit with all the experiments considered (Figure 2C-E).

We then examined the baseline dynamics of our model in terms of membrane voltage, Ca^2+^, and contractile force (Figure 3). In response to a 100 Hz stimulus (Figure 3A), we observed consistent action potentials that match the overall magnitude of a typical skeletal muscle action potential (Figure 3B). Simultaneously, Ca^2+^ accumulates within the myoplasm (Figure 3C), leading to increases in contractile force as indicated by the relative density of post-power stroke cross-bridges (Figure 3D). As expected for this high-frequency stimulus, individual contractile events (twitches) fuse and rapidly approach maximum activation, corresponding to tetanic contraction.

**Figure 3:**
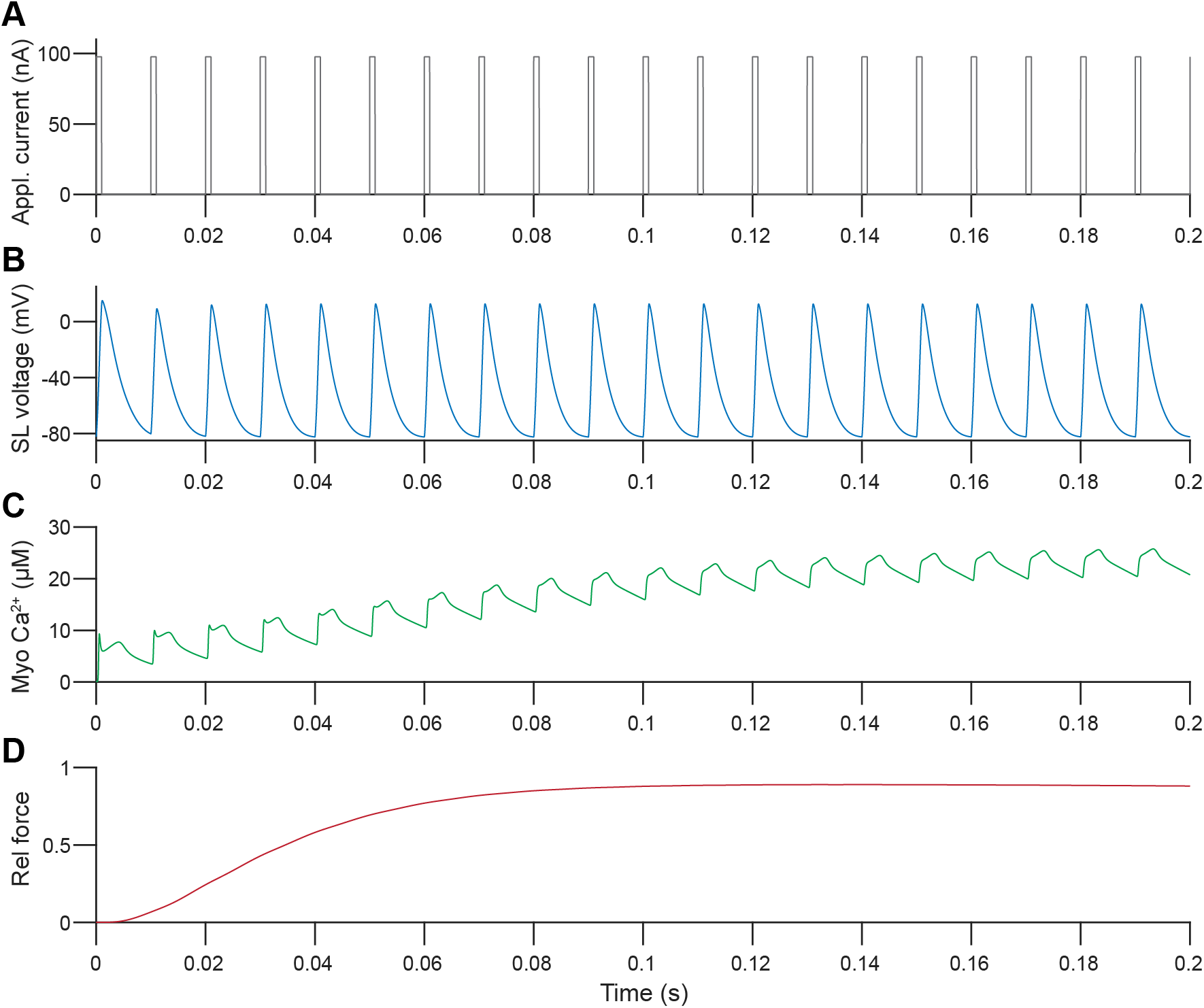
Model output for membrane potential and Ca^2+^ dynamics at 100 Hz stimulus. Input 100 Hz square wave current at the bulk sarcolemma (A) gives rise to action potentials in the sarcolemma (SL) (B) and subsequent increases in myoplasmic Ca^2+^ (C). Binding of Ca^2+^ to troponin leads to cross bridge cycling and the associated increase in contractile force (D). Here and throughout, relative force is defined as the concentration of post-power stroke cross-bridges relative to their concentration at saturating Ca^2+^ concentrations.

We then analyzed these events in terms of their constitutive currents and fluxes to determine the relative contributions of the various channels (Figure 4). Membrane depolarization leads to Na^+^ channel opening in the T-tubules and sarcolemma, followed by opening of the K^+^ delayed rectifier during repolarization (Figure 4B). The Na^+^-K^+^ pump counteracts these fluxes to restore the resting gradients of both ions, while other currents through the Na^+^-Ca^2+^ exchanger and the K^+^ inward rectifier play secondary roles (Figure 4B). In parallel, depolarization triggers activation of DHPRs in the T-tubules, which allosterically interact with RyRs in the junctional SR membrane to facilitate their opening (Figure 4B). This large Ca^2+^ release through RyRs is the dominant source for Ca^2+^ elevations in the myoplasm and results in higher concentrations in the junctional compartment compared to the bulk myoplasm (Figure A4). The features of the RyR flux itself lead to the shape of Ca^2+^ concentration curve observed here and throughout, with one initial peak followed by a secondary phase of more gradual Ca^2+^ release (Figure 4C). Myoplasmic Ca^2+^ then decreases due to buffering and removal to the extracellular, SR, and mitochondrial spaces through PMCA, SERCA, and MCU/mPTP, respectively. (Figure 4C-D). During this phase of store refilling, we observed a concomitant increase in SOCE, but *J*_SOCE_ remained small compared to other Ca^2+^ fluxes in the model (Figure 4D).

**Figure 4:**
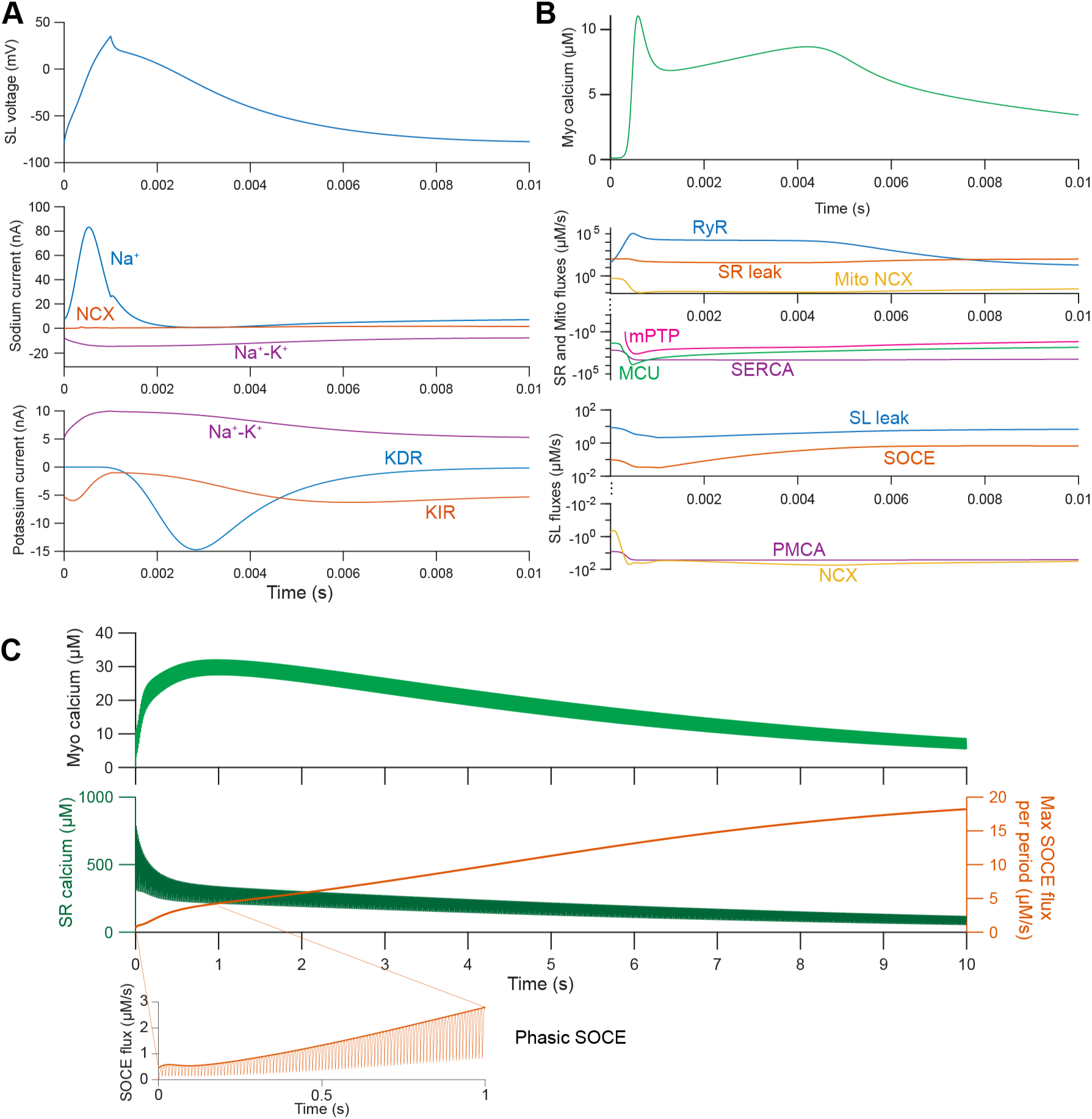
Currents and Ca^2+^ fluxes during the first action potential of a train. A) Sarcolemma voltage and Ca^2+^ concentration plotted over a single action potential, with the stimulus occurring from 0 to 0.001 s. B) Sarcolemma depolarization and repolarization coincide with an inward Na^+^ current (upper plot) and outward K^+^ delayed rectifier (KDR) current (lower plot), respectively. Currents through the Na^+^-K^+^ pump, Na^+^-Ca^2+^ exchanger (NCX), and K^+^ inward rectifier (KIR) are also plotted for reference. C) Fluxes through the SR and mitochondrial membranes, with the upper axes showing release from organelles and the lower axes showing uptake into organelles. D) Fluxes through the cell membrane (SL and TT), with the upper axes showing influxes and the lower axes showing effluxes.

We then considered the predictions of our model out to longer time points, maintaining 100 Hz stimulus for 10 s (Figure 5). During this period, as SR calcium was further depleted, SOCE continued to increase gradually (Figure 5B), suggesting that SOCE may play an increasingly important role over extended periods of stimulus. Furthermore, SOCE is activated separately in the junctional region during each action potential, in agreement with previous measurements of phasic SOCE in skeletal muscle (Lilliu et al., 2025, 2020; Koenig et al., 2018, 2019; Lilliu et al., 2021) (Figure 5C).

**Figure 5:**
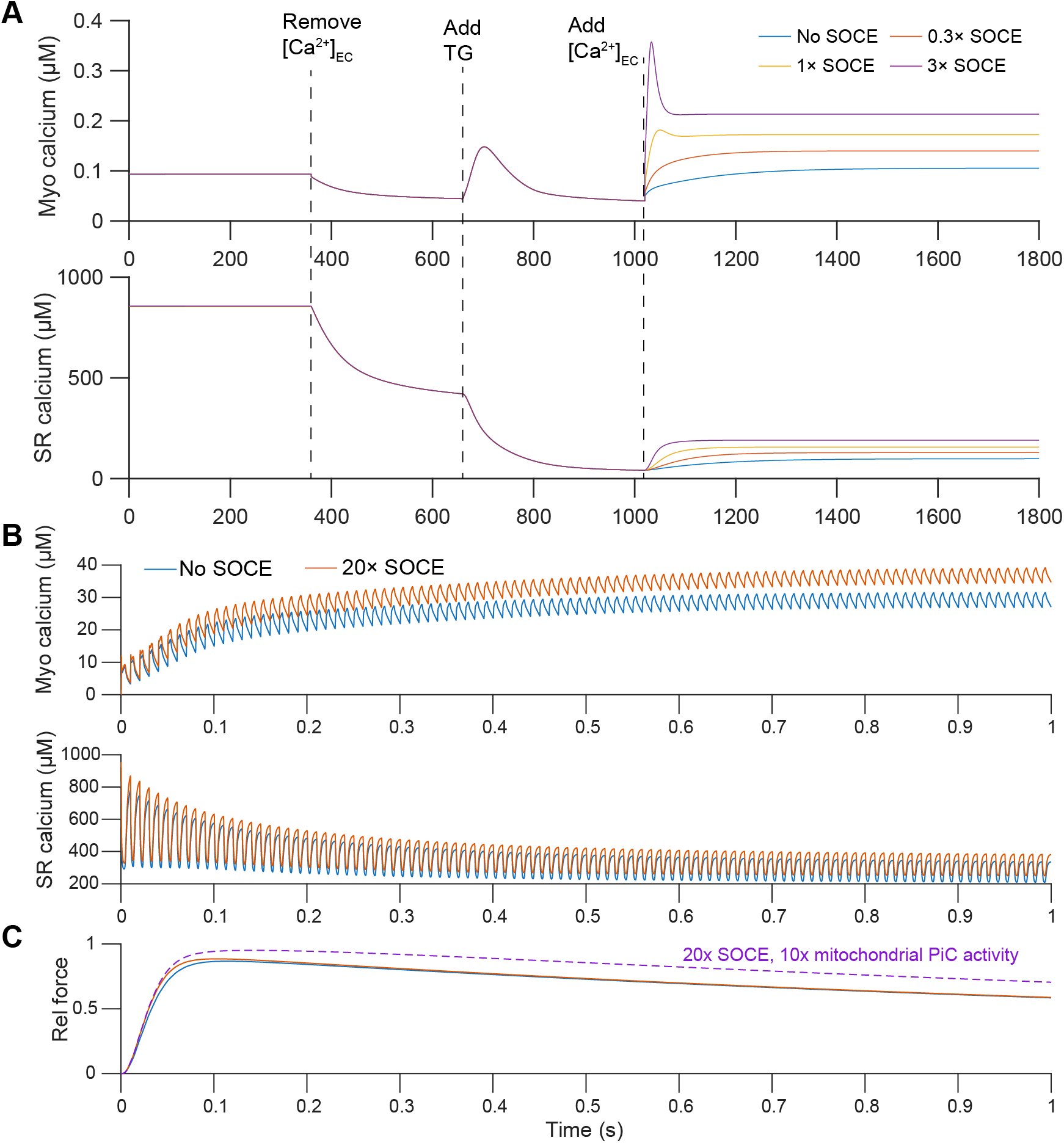
Ca^2+^ concentrations and SOCE flux during 10 s of 100 Hz stimulus. A-B) Myoplasmic Ca^2+^ (A) and SR Ca^2+^ (B) are plotted over time, with the maximum SOCE flux over each stimulus period plotted on the right *y*-axis in panel B. C) Insets showing the full dynamics of SOCE within the first 1 s of simulation and the final 1 s of simulation.

### SOCE facilitates sustained calcium elevations

Having established an agreement between our baseline model and the physiological signaling events in skeletal muscle fibers, we systematically examined the importance of SOCE for Ca^2+^ dynamics and force production. We first considered the treatment of cells with the SERCA inhibitor, thapsigargin (TG), commonly used to assess SOCE across cell types (Nelson et al., 2020; Hoover and Lewis, 2011; Luik et al., 2008; Rabesahala de Meritens et al., 2025). In such experiments, cells are placed in Ca^2+^-free media and then treated with TG, causing Ca^2+^ to leak from the SR into the myoplasm and slowly exit the cell due to efflux through PMCA. Ca^2+^ is then reintroduced into the media, leading to a large store-operated Ca^2+^ influx. This experiment was previously conducted in L6 myoblasts, (Nelson et al., 2020), revealing that the increase in myoplasmic Ca^2+^ following TG treatment was smaller and more short-lived than the elevation due to store-operated Ca^2+^ influx once extracellular Ca^2+^ was reintroduced (see Fig 5 in (Nelson et al., 2020)). We recapitulated this experiment in our simulations by altering extracellular Ca^2+^ and SERCA activity in a time-dependent manner as indicated in Figure 6A. Our simulations show agreement with the measurements by (Nelson et al., 2020), with a smaller TG-induced flux followed by extended SOCE, with the peak Ca^2+^ concentration determined by the strength of SOCE. Notably, our model allowed us to predict the dynamics of other quantities not readily accessible in experiments, such as concentration of Ca^2+^ in the SR (Figure 6A).

**Figure 6:**
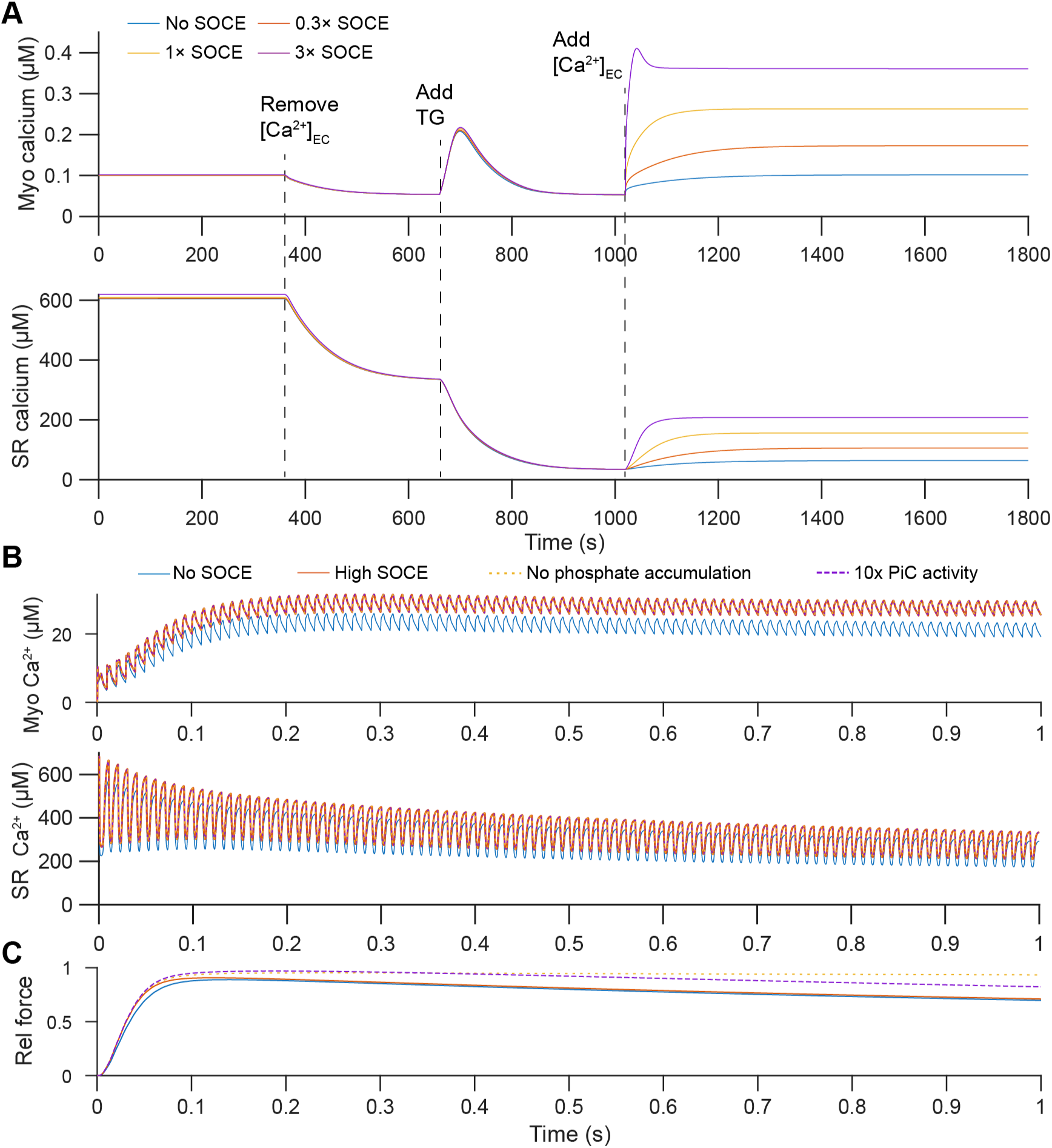
Crosstalk between SOCE and phosphate handling in computational experiments. A) Predicted myoplasmic (upper) and SR (lower) Ca^2+^ dynamics in a conventional thapsigargin (TG) assay, modeled after tests in (Nelson et al., 2020). Extracellular Ca^2+^ is initially 1300 µM, then reduced to 0 µM at *t* = 360 s (e.g., via addition of a Ca^2+^ chelator such as EGTA). Addition of thapsigargin at *t* = 660 s reduces SERCA activity to 10% of its resting activity, inducing Ca^2+^ release from the SR. Extracellular Ca^2+^ (1300 µM) is then reintroduced at *t* = 1020 s, triggering SOCE, with Orai1 conductance varied relative to the baseline (uncalibrated) value in Table A7. B) Predicted myoplasmic (upper) and SR (lower) Ca^2+^ dynamics in response to 100 Hz stimulus with high SOCE (*g*_SOCE_ = 5000 fS per square micron) vs. no SOCE. Results are also shown for high SOCE and either no phosphate accumulation or 10 times baseline PiC transport rate (these conditions overlap with the high SOCE curves here). C) Predicted relative contractile force for conditions shown in panel B.

We then considered stimulation of a myofiber at 100 Hz with and without SOCE. We increased Orai1 conductance to represent enhanced SOCE activity in skeletal muscle. This increase can be attributed to channel aggregation, or to formation of CEUs outside the triadic junction, as suggested by recent experiments (Boncompagni et al., 2017; Michelucci et al., 2019; Protasi et al., 2021). We determined a reasonable value for increased conductance by assuming a total density of about 200 receptors per square micron (within the range of measurements of Orai1 density in STIM1-Orai1 puncta (Peckys et al., 2021)) and single channel conductance of 10 fS (Amcheslavsky et al., 2015), resulting in a total conductance of 2000 fS per square micron in the T-tubules. In this test, SOCE played a significant role in Ca^2+^ dynamics – the increased Orai1 conductance led to a 21.3% increase in peak myoplasmic Ca^2+^ (Figure 6B) We also examined the contractile force exerted by fibers with and without SOCE. The predicted dynamics of contractile force were similar in both cases, with only a minor (1.98%) increase in the maximum relative force for cells with strong Orai1 conductance over cells with no SOCE.

With or without SOCE, contractile force declined after less than 1 s of stimulation. This effect was largely attributed to phosphate accumulation in the myoplasm, as fixing the myoplasmic phosphate to a constant value almost entirely eliminated the observed force reduction (Figure 6C, Figure A5). This agrees well with the established role of phosphate in muscle fatigue (Shorten et al., 2007; Cheng et al., 2018; Allen and Trajanovska, 2012; Allen et al., 2008; Hendry et al., 2025). In our model, one of the main mechanisms for phosphate clearance from the myoplasm is transport through the mitochondria phosphate carrier (PiC). We hypothesized that changes in the rate of phosphate uptake by mitochondria (dictated by *ν*_PiC_, the permeability of mitochondria PiC) could provide an important factor modulating fatigue in our model. To test this, we increased *ν*_PiC_ by a factor of 10. Without altering Ca^2+^ dynamics, this led to much lower accumulation to phosphate in the myoplasm (Figure A5), and, consequently, partially mitigated the decrease in force (Figure 6C). Given the established interplay between SOCE and mitochondrial network morphology (Nan et al., 2021), changes in phosphate transport through PiC may provide another mechanism for SOCE-mediated modulation of fatigue, as explored in further detail below.

### SOCE and phosphate transport modulate contractile force in a frequency-dependent manner

Our initial simulations suggested that SOCE is an important regulator of Ca^2+^ dynamics and force production in skeletal muscle, in conjunction with changes in mitochondrial phosphate uptake. We next used our model to predict whether the effects of SOCE depend on stimulus frequency. We considered a case previously tested experimentally, in which dominant-negative Orai1 (dnOrai1) mouse EDL muscle exhibited frequency-dependent inhibition of force generation (Wei-LaPierre et al., 2013). As in these experiments, we applied different frequencies of stimuli consisting of square-wave pulse trains for a total of 0.5 s each. In each test, we allowed basal levels of phosphate accumulation by running the simulation for an initialization period of 10 s without stimulus, as shown in Figure A6.

At low stimulus frequencies (less than 20 Hz), we observed low force production regardless of SOCE, as tetani remained unfused (Figure 7A). SOCE did enhance force production by a small amount at moderate stimulus frequencies (Figure 7B), but had almost no effect on force production at higher (*>* 80 Hz) frequencies (Figure 7C). Compared to the experimental data, these simulations generally overestimated contractile force in SOCE knockout fibers at high frequencies (Figure 7C-D). We reasoned that additional SOCE-regulated processes not considered in our model might contribute increasingly at higher stimulus frequencies. SOCE mutants have been known to exhibit increased mitochondrial fragmentation and volume loss (Maus et al., 2017; Choi et al., 2019; Collins et al., 2014), which can disrupt the normal proton gradient (Alan and Scorrano, 2022) and accordingly, reduce phosphate transport through PiC. We therefore reduced the PiC activity by a factor of 10 in SOCE-deficient cells, resulting in a significant reduction in force at higher stimulus frequencies in agreement with experiments (Figure 7C-D). This reinforces findings from our initial simulations in Figure 6, suggesting that SOCE and PiC activity co-modulate force reduction in myofibers.

**Figure 7:**
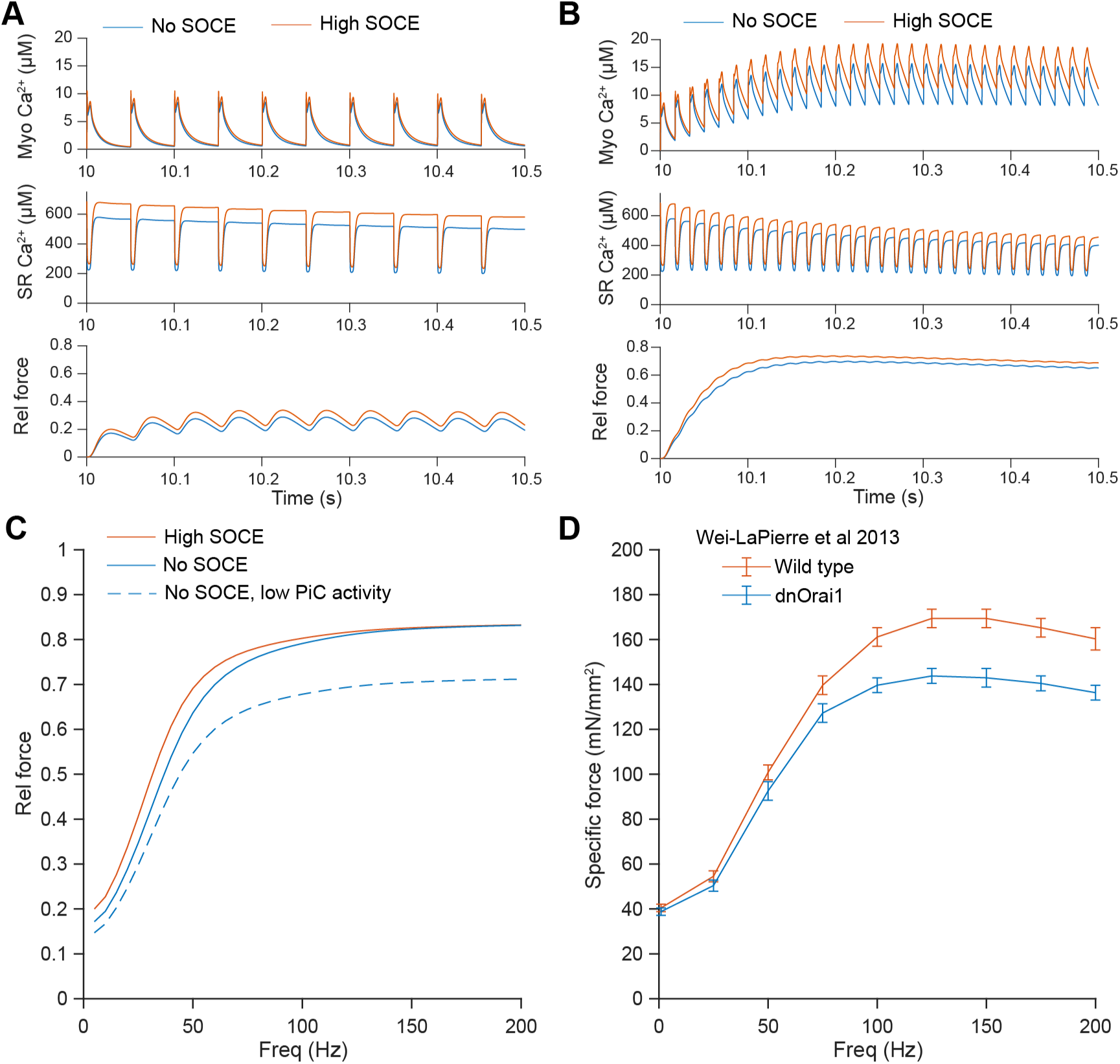
Frequency-dependent force generation with vs. without SOCE. A-B) Predicted effects of SOCE on myoplasmic Ca^2+^, SR Ca^2+^, and contractile force are plotted for 20 Hz (A) and 60 Hz (B) stimuli. C) Frequency-dependent maximum contractile force with and without SOCE. Dashed line indicates case without SOCE and with 10% PiC activity. D) Experimental measurements of maximum force during in vitro testing of muscle from the EDL of wild type or dominant negative Orai1 (dnOrai) mice (Wei-LaPierre et al., 2013). *n* = 14 for both wild type and dnOrai1 conditions and error bars denote standard error of the mean. Data were extracted from the original figure using manual analysis in ImageJ.

### Computational predictions suggest context-dependent role of SOCE in different forms of exercise

Finally, we considered the predictions of our model for different frequencies and patterns of stimuli relevant to resistance exercise and high-intensity interval training (HIIT). EMG measurements suggest that these exercises are marked by different stimulus frequencies at the NMJ – about 10-50 Hz for resistance exercise (Pucci et al., 2006; Bellemare et al., 1983) and 60-180 Hz for HIIT (Ament et al., 1993). Some studies indicate that firing frequencies can increase up to about 70-100 Hz in certain cases of rapid maximal volumetric contraction (MVC) (Škarabot et al., 2024). Accordingly, we modeled high-load (approaching 100% MVC) resistance exercise using either 40 Hz or 70 Hz stimuli, with 3 s of contraction followed by 3 s of rest for each repetition, modeled after the isometric contraction measured in the quadriceps in previous work (Pucci et al., 2006). During high-intensity exercise such as sprinting, stimulus of a given muscle occurs in discrete phases during each stride. Considering running data collected in muscles that have a high abundance of fast twitch fibers (gastrocnemius medialis or the gastrocnemius lateralis) (Hamner and Delp, 2013), we assumed that stimulus occurs over a 0.1625 s time interval for each 0.65 s stride, with stimulus frequencies of either 70 or 120 Hz. In the following tests, we considered stimulus trains designed to mimic 20 s of running or 10 repetitions (60 s total) of resistance exercise.

To examine the role of SOCE in force production during exercise, we varied Orai1 conductance (*g*_Orai1_) and maximized STIM1 Ca^2+^ sensitivity (*K*_STIM1_ = 0.5[Ca^2+^]_SR,_*_∞_*, where [Ca^2+^]_SR,_*_∞_* is the steady-state SR Ca^2+^ concentration). Shifts in *g*_Orai1_ can be attributed changes in SR/T-tubule organization (Boncompagni et al., 2017), transport of STIM1 (Wu et al., 2007; Grigoriev et al., 2008), or Orai receptor turnover (Yeh et al., 2020). Changes in *K*_STIM1_ are associated with biochemical modifications (e.g., phosphorylation) that alter STIM1-Orai1 binding or STIM1 autoinhibition (Nelson et al., 2020; Pozo-Guisado and Martin-Romero, 2013). To examine the role of SOCE in fatigue, we simultaneously varied other parameters that were important determinants of fatigue – the troponin-Ca^2+^ binding rate (*k*_on,T1_, determining the cross-bridge Ca^2+^ sensitivity) and the permeability of mitochondrial PiC (*ν*_PiC_).

In the case of resistance exercise, we found that force production was enhanced by elevating Orai1 conductance (Figure 8A-B). This effect held at both stimulus frequencies tested and regardless of the values for *k*_on,T1_ and *ν*_PiC_ (Figure A7). Mechanistically, these higher forces were attributed to a significantly higher fraction of troponin binding to Ca^2+^ in the myoplasm, allowing for faster crossbridge cycling despite increased accumulation of myoplasmic phosphate (Figure 8A). The rate of phosphate uptake by mitochondria did not influence the final force output for low levels of SOCE, but at higher Orai1 conductance, increased rates of PiC activity were required for optimal force production (Figure 8B, Figure A7).

**Figure 8:**
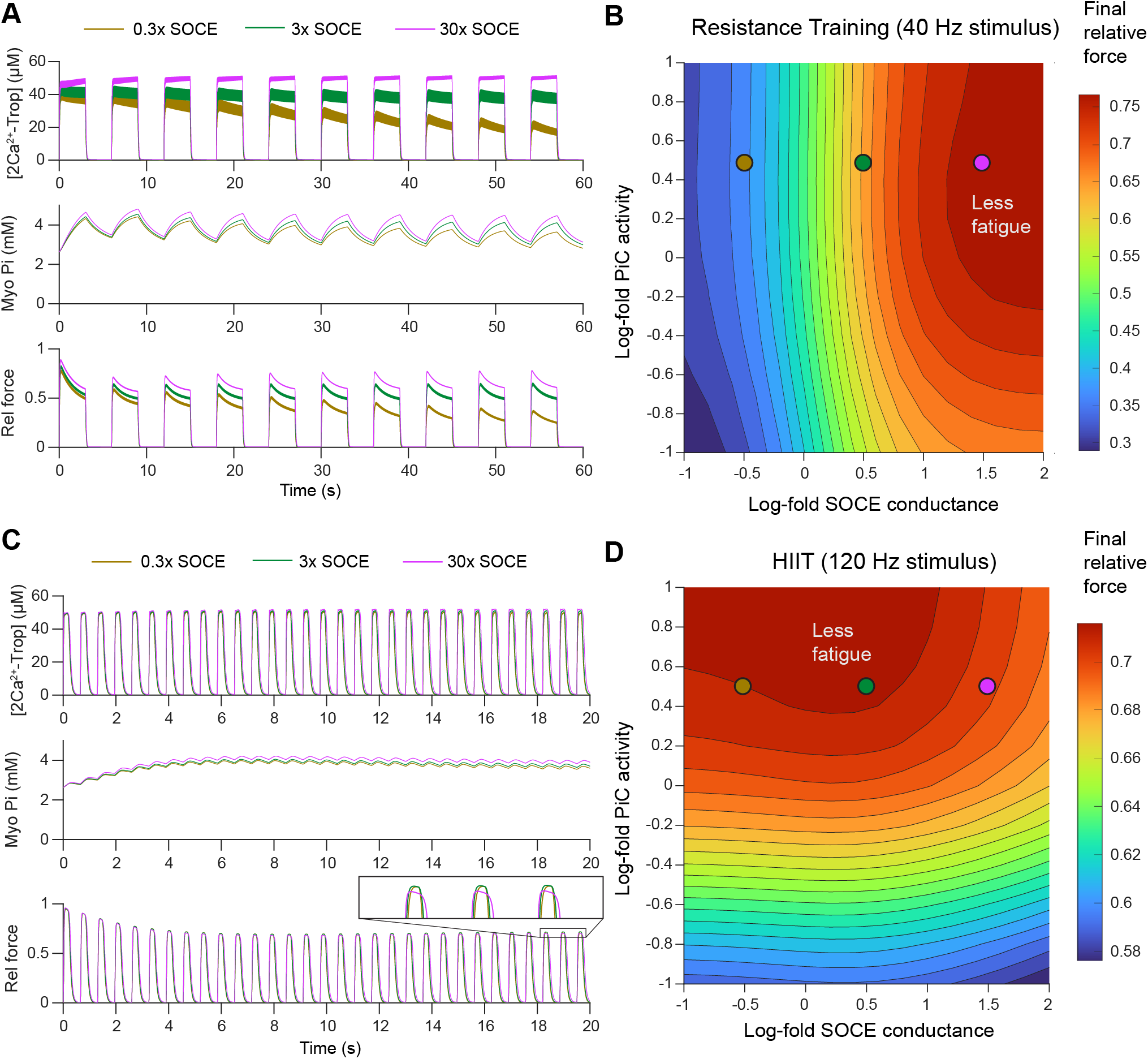
Predicted effects of altering SOCE and phosphate uptake rate in different exercise regimes. A) Predicted dynamics of [Trop-2Ca^2+^], myoplasmic phosphate, and force over time during patterns of 40 Hz stimuli designed to mimic vastus lateralis activity during resistance training (Pucci et al., 2006). B) Phase diagram summarizing the maximum contractile force during the final repetition of a 10-repetition resistance training set as a function of PiC permeability (*ν*_PiC_) and SOCE conductance (*g*_Orai1_). C) Predicted dynamics of [Trop-2Ca^2+^], myoplasmic phosphate, and force over time during patterns of 100 Hz stimuli designed to mimic gastrocnemius medialis activity during running (Hamner and Delp, 2013). D) Phase diagram summarizing the maximum contractile force during the final stride of a 20 s running interval as a function of *ν*_PiC_ and *g*_Orai1_. Tests in panels A and C are conducted for low (0.3×), moderate (3×), and high (30×) the original (uncalibrated) Orai1 conductance at 3×*ν*_PiC_, as indicated by points on the phase diagrams in panels B and D. All tests shown here used *K*_STIM1_ = 0.5[Ca^2+^]_SR,_*_∞_* and 3 times the baseline estimated value for *k*_on,T1_.

Surprisingly, increasing levels of SOCE did not always increase the final force in HIIT (Figure 8C-D). Rather, at higher rate of phosphate uptake, increasing Orai1 conductance actually reduced the force at the final stride (Figure 8C-D). At very high levels of SOCE, the fraction of troponin bound to Ca^2+^ was already saturated, and so further increases in SOCE reduced force production due to phosphate accumulation (Figure 8C). This effect was particularly apparent at the highest stimulus frequency (120 Hz) and highest Ca^2+^-troponin binding rate (Figure A8). In these simulations, increasing PiC activity has a crucial modulatory effect – fatigue was only minimized when the rates of phosphate clearance were sufficiently high (Figure 8D). Thus, while increased SOCE often enhances force output, very high rates of Ca^2+^ entry can worsen fatigue. This reveals an emergent interplay between SOCE and mitochondrial phosphate handling in excitation-contraction coupling in skeletal muscle.

## Discussion

Given the well-established role of Ca^2+^ release in skeletal muscle contraction, understanding conditions that optimize Ca^2+^ release and contractile force generation in exercise, and how such factors contribute to muscle failure in fatigue is important for establishing predictive relationships between Ca^2+^ dynamics and athletic performance. Here, we investigate the particular role of store-operated Ca^2+^ entry in excitation-contraction coupling in skeletal muscle. While many recent experimental studies had suggested a central role of SOCE in enhancing force and mitigating fatigue (Boncompagni et al., 2017; Wei-LaPierre et al., 2013), a majority of the existing computational models of excitation-contraction coupling in skeletal muscle did not include this important pathway. Our model not only addresses this gap in the literature but combines action potential machinery with detailed Ca^2+^and phosphate handling, constrains model parameters to experimental data, and generates testable predictions about the impact of SOCE on fatigue in skeletal muscle.

Our model predicts that SOCE can significantly enhance elevations in myoplasmic Ca^2+^ and contractile force, in line with our original hypothesis (Figures 6 and 7). However, in contrast to our initial expectations, we found that this relationship was not consistent across all parameter regimes, which helps resolve an apparent contradiction in the current literature regarding SOCE in skeletal muscle. Multiple studies have measured reduced force and increased susceptibility to fatigue in SOCE-deficient mice (Pan et al., 2002; Wei-LaPierre et al., 2013). In contrast, recent data suggests that SOCE deficiency might actually mitigate fatigue in certain cases (Nelson et al., 2020). Here, we show that these two effects can be understood as the outcomes of the same biophysical system operating in different regimes. At lower stimulus frequencies and higher rates of phosphate uptake by mitochondria, we predict that SOCE enhances Ca^2+^ levels, leading to increased contractile force. However, for high-frequency stimulus, increasing SOCE can lead to reduced force due to accumulation of phosphate in the myoplasm. In this case, reducing SOCE mitigates fatigue, in agreement with experiments that show silencing STIM1 or inhibiting it via phosphorylation results in less susceptibility to fatigue (Nelson et al., 2020). These shifts may also be attributable to organismal differences since some studies consider mice (Pan et al., 2002; Wei-LaPierre et al., 2013), while Nelson et al examine fatigue in Drosophila (Nelson et al., 2020).

The interplay between SOCE and phosphate dynamics within myofibers played an important role throughout our simulations. Previous models included phosphate cycling in the myoplasm and SR (Senneff and Lowery, 2021; Shorten et al., 2007; Senneff and Lowery, 2022), but we additionally account for a major mode of phosphate uptake through the phosphate transporter in the mitochondrial inner membrane (Seifert et al., 2015). Not only does this provide a natural mechanism to mitigate fatigue in our model, but phosphate also serves as a buffer for mitochondrial Ca^2+^, allowing for increased Ca^2+^ uptake through the MCU. Increased levels of mitochondrial Ca^2+^ leads to increased rates of ATP production via oxidative phosphorylation. Our work suggests that mitochondrial phosphate handling is a key determinant of whether SOCE alleviates or worsens fatigue in myofibers. This connection is especially relevant given the established regulatory role for SOCE in mitochondrial fission and fusion (Nan et al., 2021). High levels of SOCE activity lead to increased mitochondria Ca^2+^ levels, which can induce fission (Han et al., 2008) or disruptions to mitochondrial morphology (Walkon et al., 2022). Such effects may perturb phosphate transport and increase the occurrence of SOCE-induced fatigue.

The control mechanisms that determine the relationship between SOCE and phosphate transport in myofibers require detailed considerations of metabolic regulation in skeletal muscle, necessitating models that extend beyond the time range considered here. Many such responses rely on metabolic regulation by AMP-activated protein kinase (AMPK), which previous models have shown to be modulated by the duration of stimulus during exercise (Linden-Santangeli et al., 2025b; Fowler et al., 2024). One recent computational study considered the resulting changes in mitochondrial fission and fusion downstream of different types of exercise, including inputs similar to those tested for resistance and HIIT here (Khalilimeybodi et al., 2026). This model predicted enhanced activation of mitochondrial fusion downstream from HIIT, suggesting a possible compensatory mechanism for the effects of fatigue observed in this work. Long-term responses can also involve volumetric growth of the muscle itself, as considered in other recent modeling work (Devold et al., 2025; Villota-Narvaez et al., 2021). Bridging the time and length scales between the single-fiber, short timescale considered here to models of long-term remodeling and growth at the whole muscle scale presents a distinct challenge for future multiscale modeling work.

In addition to their important roles in Ca^2+^ handling and mitochondrial fission and fusion, STIM and Orai also regulate other intracellular processes. STIM1 has been shown to engage a variety of other binding partners in addition to Orai1, including POST (partner of STIM1), which facilitates its interactions with SERCA, PMCA, the Na^+^-K^+^ pump, and nuclear transporters (Krapivinsky et al., 2011; Hooper et al., 2013). The multiplicity of roles for SOCE likely has many implications for the development of STIM and Orai germline knockout animals. Accordingly, the experimental data considered in Figure 7 must be interpreted carefully. Indeed, inducible Orai1 knockout mice do not exhibit the same reduction in maximal force during repeated stimulus, but still exhibit increased fatigability (Carrell et al., 2016; Michelucci et al., 2019). Additional work is required to untangle the effects of acute SOCE dysfunction versus effects observed due to lack of SOCE during development.

While our model incorporates many of the important players in excitation and contraction in skeletal muscle, we note some opportunities for future developments. We acknowledge that our treatment of SOCE is simplified, consisting of a single ODE governing Orai1 open probability; the detailed dynamics of STIM1 oligomerization, diffusion, and association with Orai1 would require a more detailed biophysical model. Furthermore, other channels such as transient receptor potential canonical proteins (TRPCs) can contribute to SOCE in skeletal muscle through interactions with STIM1 (Choi et al., 2020) and may be included in future models. However, detailed models introduce many free parameters, requiring additional experimental data for robust model calibration (Linden et al., 2022; Qiao et al., 2025). All of our parameters are currently estimated using data collected in single mouse myofibers at room temperature over a maximum of five action potentials (Baylor and Hollingworth, 2007; Miranda et al., 2020; Rincón et al., 2021); new datasets are needed to reliably calibrate models of human myofibers at physiological temperatures over extended times. Such data will better constrain many of the parameters to which the model was not currently sensitive, and may present opportunities for other approaches from uncertainty quantification such as Bayesian parameter estimation (Linden et al., 2022; Qiao et al., 2025) or Bayesian model selection (Linden-Santangeli et al., 2025b,a). Finally, future work is required to elucidate the spatial features of SOCE in skeletal muscle. Experimental data show that myofibers undergo dramatic structural reorganization following exercise, including changes in the T-tubule system and assembly of CEUs (Boncompagni et al., 2017; Cully et al., 2017) While our model differentiates between junctional and bulk regions within the myofiber, spatial modeling approaches using software packages such as VCell (Cowan et al., 2012) or SMART (Francis et al., 2025) are required to properly explore this aspect of Ca^2+^ signaling in skeletal muscle. Such models provide a natural opportunity to further explore how the transport of Ca^2+^ and phosphate into mitochondria depends on features of their morphology altered by fission or fusion. Additionally, spatial models can help elucidate the importance of structural features of the NMJ, which have been shown to significantly impact the strength of signal transduction (Santoso et al., 2021).

Our findings may have implications beyond skeletal muscle, including neurons and cardiomyocytes (Basnayake et al., 2021; Saftenku, 2022; Hermes et al., 2023; Benedetti et al., 2025), as well as nonexcitable cells such as immune cells (Clemens and Lowell, 2015; Vig and Kinet, 2009). For example, the frequency-dependent role of SOCE explored here may naturally extend to postsynaptic signaling in neuronal dendritic spines (Basnayake et al., 2021; Dembrow and Spain, 2022). Thus, our study positions SOCE as a key regulator of SR and myoplasmic Ca^2+^, particularly important for cellular events that trigger rapid store depletion.

## Materials and Methods

### 0.1 Ethical approval

No human or animal data were collected for this work, all experimental data shown was from previous studies as indicated.

### Numerical implementation

The full model is described in Supplementary Information (Tables A1 to A7) and consists of 65 nonlinear ordinary differential equations. Initial conditions were estimated by starting from default values of each variable (Table A4) and integrating the equations with no input current to *t* = 5000 s in MATLAB using the stiff ODE solver, ode15s. The initial values were then either defined by the values at *t* = 5000 s or the values when the following condition was met:

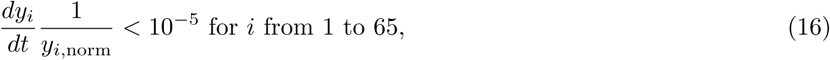

where *y_i_*is the state variable associated with index *i* and *y_i,_*_norm_ is a normalizing factor for *y_i_*, defined in Table A4.

In all cases, the steady-state solution with no SOCE (*g*_Orai1_ = 0) was computed before defining *K*_STIM1_ as a fraction of the resting SR Ca^2+^ concentration without SOCE (Table A7). This approach was adopted to avoid cases with unrealistic levels of SOCE under resting conditions. Steady-state values for each state variable were then estimated for the desired Orai1 conductance (*g*_Orai1_ *>* 0) at the assigned value of *K*_STIM1_, before solving the dynamics with various stimuli using ode15s. This strategy was used to compute steady-state for all variables except concentrations of phosphate, ADP, and ATP in each compartment, which were fixed to estimated parameter values (for myoplasmic ATP, myoplasmic phosphate, and SR phosphate) or to default values (for myoplasmic ADP and for mitochondrial ATP and phosphate) before starting the simulation (Table A7). SR Ca^2+^-phosphate precipitate was always assumed to start at 0 µM. MATLAB code from this study is freely available on Github and through Zenodo (Kumar et al., 2025).

### Sensitivity Analysis

Morris sensitivity analysis was conducted using the UQLab framework in MATLAB (Marelli and Sudret, 2014). As described in the main text, maximum and average values of myoplasmic Ca^2+^ and sarcolemma voltage were used as quantities of interest (QOIs). All non-mitochondrial parameters were varied according to a uniform distribution, from 0.5 to 2 times the reference value from literature (Table A7) Morris analysis yielded the absolute value of the mean (µ*^∗^*) and the standard deviation (*σ*) of the elementary effects. A parameter was treated as sensitive if µ*^∗^* was greater than 20% of the maximum µ*^∗^* over all parameters for that QOI.

### Parameter Estimation

To ensure good fits to sarcolemma voltage, myoplasmic Ca^2+^, and mitochondrial Ca^2+^, parameter estimation was conducted in three stages. First, mitochondria-related parameters were independently calibrated in a mitochondria-exclusive test described below. We carried these parameters to the full model and then fit all the voltage-related parameters to match physiological action potentials. In the final step, model parameters were further constrained by fitting the full model to both myoplasmic Ca^2+^ and sarcolemma voltage over time.

#### Mitochondrial parameter estimation

The model of mitochondrial Ca^2+^ and phosphate handling was first fit separately from the rest of the model. We considered experimental data from Wei et al (Wei et al., 2015), in which the authors measured mitochondrial Ca^2+^ (both free and bound) and mitochondrial membrane voltage over time in vitro. In these experiments, guinea pig heart mitochondria were isolated and initially depleted of phosphate and placed in a phosphate-free, Ca^2+^-low solution.

They then used fluorescence microscopy to simultaneously measure free matrix Ca^2+^, external Ca^2+^, and membrane voltage. Monitoring external Ca^2+^ together with free matrix Ca^2+^ allowed them to monitor the ratio of bound to free Ca^2+^ in the mitochondrial matrix. Each experiment started with adding Ca^2+^ to the external solution (shown at *t* = 110 s in Figure A3A), leading to Ca^2+^ uptake by mitochondria and an associated reduction in the mitochondrial membrane voltage. They then added free phosphate (either 1, 10, 100, or 1000 µM) to the solution (at *t* = 240 in Figure A3A), leading to further Ca^2+^ uptake and a dramatic increase in the ratio of free to bound matrix Ca^2+^, illustrating the role for matrix phosphate as a Ca^2+^ buffer. We used our model to simulate these experiments, applying external Ca^2+^ and phosphate as known quantities. The system was initialized by starting with [Ca^2+^]_Mito_ = 0.1 µM, [ATP]_Mito_ = 10 *×* 10^3^ µM, [NADH]_Mito_ = 50 µM, *V*_Mito_ = 150 mV, and no mitochondrial phosphate or Ca^2+^-phosphate, then running the system for 400 s with no applied phosphate or Ca^2+^ in the extra-mitochondrial solution. Model predictions could then be directly leveraged against their data (Figure A3C-D), with the overall error computed by the following weighted sum of square errors:

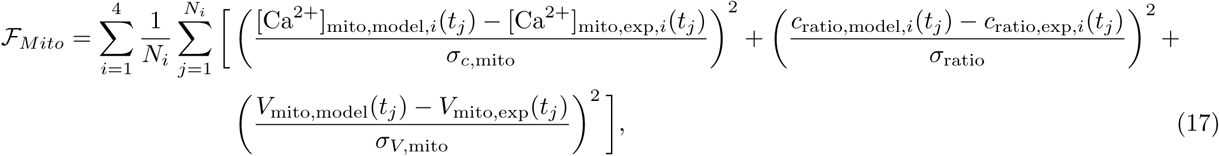

where *N_i_* is the number of time points considered for experiment *i* and the weighting factors (*σ_c,_*_mito_*, σ*_ratio_*, σ_V,_*_mito_ were chosen to represent measurement error (we chose *σ_c,_*_mito_ = 5 µM, *σ*_ratio_ = 100, and *σ_V,_*_mito_ = 10 mV, based on the magnitude of the standard error of the mean from (Wei et al., 2015)).

To compute *c*_ratio_ from our model, we accounted for both the rapid buffering and Ca^2+^ binding to phosphate in the matrix. The rapid-buffering approximation here assumes that, of the Ca^2+^ not bound to phosphate, a fraction 1 *− b_M_* is bound to other buffers; that is:

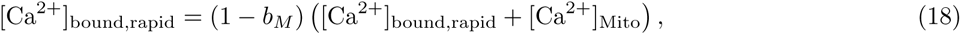

where [Ca^2+^]_bound,rapid_ is the concentration of Ca^2+^ bound to buffers other than phosphate.

The ratio of bound-to-free Ca^2+^ is

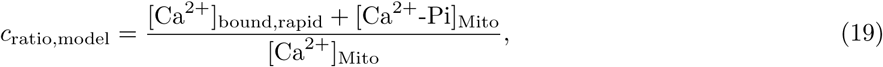

which, upon substituting Equation (18), yields the expression used to compute the bound-to-free ratio in our model:

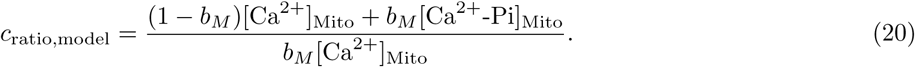

This fit was conducted using particle swarm optimization (PSO) in MATLAB with a swarm size of 30 and maximum stall iterations of 20. We allowed each parameter to vary from 0.5 to 2 times its baseline value, except for voltage-shift parameters (parameters M17 and M23 in Table A7), which could only vary *±* 10 mV. The fit reproduced key qualitative features of the experimental measurements (Figure A3C-D) with the exception of the mitochondrial membrane voltage, which decreased at the end of simulations with higher added amounts of phosphate. We attribute this to negative ion fluxes across the mitochondrial membrane not accounted for in our model, which are outside the scope of this study.

#### Membrane voltage parameter estimation

In the following step, the objective function was defined as the normalized sum of square errors between experimentally measured voltage and model predictions for the data from (Miranda et al., 2020):

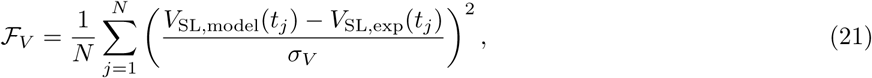

where *V*_SL,model_(*t_j_*) and *V*_SL,exp_(*t_j_*) are, respectively, the predicted and experimentally measured sarcolemma voltage at time *t_j_*. *N* is the number of time points considered and *σ_V_* is the standard deviation of the measurement error (fixed at 5 mV). For this step, only the 27 parameters with greater than 20% of the maximum sensitivity index for average or peak voltage (Figure A2A-B) were included in the fitting. Each of these parameter was allowed to vary between 0.5 and 2 times its baseline value, except for voltage-shift parameters (parameters 76-83 in Table A7), which could only vary *±* 10 mV. Fitting was conducted using particle swarm optimization in MATLAB, with a swarm size of 30 particles and maximum stall count of 20.

#### Full model parameter estimation

In the final step, a new objective function was defined, starting with the weighted sum of square errors between experimental and simulated myoplasmic Ca^2+^:

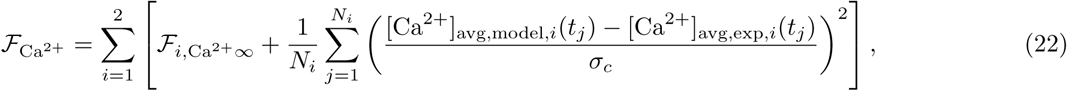

where [Ca^2+^]_avg,model,*i*_(*t_j_*) and [Ca^2+^]_myo,exp,*i*_(*t_j_*) are the predicted (defined in Equation (15)) and experimentally measured spatially averaged myoplasmic Ca^2+^ concentration at time *t_j_* for experiment *i* and *σ_c_* is the standard deviation of the measurement error (fixed at 0.5 µM). *F_i,_*_Ca_2+*_∞_* represents an additional fitting constraint for each experiment that ensures a good match with experimentally characterized myoplasmic and SR Ca^2+^ at steady-state:

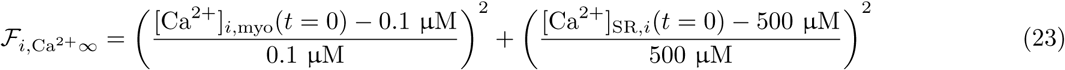

A total objective function was constructed to ensure good fits with both Ca^2+^ and voltage data simultaneously, *F_V_* + *F*_Ca^2+^_. All parameters were included in this fitting step, starting with initial guesses given by the previous estimation for applicable parameters. We used the calibrated values from the previous step as new baseline values for applicable parameters here. All parameters were allowed to vary between 0.5 and 2 times their baseline values, except for voltage-shift parameters (parameters 76-83 in Table A7), which could only vary *±* 10 mV. The same specifications for particle swarm optimization were used as in the previous step. The final optimized values for each parameter are reported in Table A7.

## Additional Information

### Data Availability

Our code, with all data necessary to reproduce figures in this manuscript, is freely available on Github and is deposited on Zenodo (Kumar et al., 2025).

### Competing Interests

P.R. is a consultant for Simula Research Laboratory in Oslo, Norway and receives income. The terms of this arrangement have been reviewed and approved by the University of California, San Diego in accordance with its conflict-of-interest policies.

### Author Contributions

E.A.F.: Conceptualization, Funding Acquisition, Supervision, Data Curation, Formal Analysis, Investigation, Methodology, Software, Visualization, Writing Original Draft, Review and Editing; J.H.: Conceptualization, Data Curation, Formal Analysis, Investigation, Methodology, Software, Visualization, Review and Editing; A.K.: Conceptualization, Data Curation, Formal Analysis, Investigation, Methodology, Software, Visualization, Writing Original Draft, Review and Editing; P.R.: Conceptualization, Funding Acquisition, Resources, Project Administration, Supervision, Review and Editing.

### Funding

This work was supported in part by the Wu Tsai Human Performance Alliance at the University of California, San Diego (to P.R.). E.A.F. was supported by the National Science Foundation under grant EEC-2127509 to the American Society for Engineering Education, by the Wu Tsai Human Performance Alliance, and by the National Institutes of Health, grant number 1K99GM162992-01A1.

### Use of Generative AI

No generative AI tools were used in the preparation of this manuscript.

## Acknowledgments

The authors would like to thank Ingvild Devold for her careful review and helpful feedback on our manuscript. We would also like to thank Profs. Samuel Ward, Andrew McCulloch, and Simon Schenk for insightful conversations on skeletal muscle biology and exercise. Simulation results presented in this paper benefited from the Triton Shared Computing Cluster (San Diego Supercomputer Center) and the Expanse Computing Cluster (Strande et al., 2021) at the San Diego Supercomputer Center.

## Appendix A Full model description

In our model, four major compartments (extracellular space, myoplasm, mitochondria, and SR lumen) are each divided into two well-mixed regions - one “junctional” region and one “bulk” region. Each junctional region is adjacent to the SR-T-tubule junction (either T-tubule volume, terminal myoplasm, junctional mitochondria, or terminal SR) and is treated as a separate compartment due to the highly localized ionic fluxes in the region. The myoplasm and extracellular space are separated by the plasma membrane (consisting of T-tubules and sarcolemma), whereas the myoplasm and SR lumen are separated by the SR membrane. In contrast, the junctional and bulk regions of each major compartment other than mitochondria are not separated by membranes. Accordingly, we assume that fluxes across the associated interface (denoted by dashed lines in Figure 1) are purely diffusive. That is, for a given species *c*:

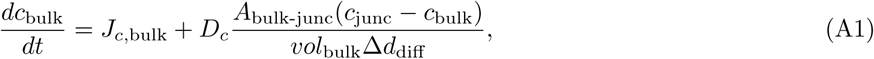

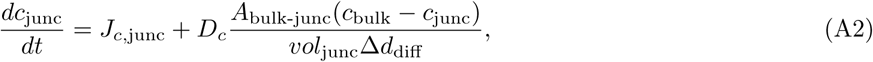

where Δ*d*_diff_ is the characteristic length for diffusion, *vol*_bulk_ and *vol*_junc_ are the volumes of the bulk and junctional regions, respectively, and *A*_bulk-junc_ is the area of the interface between the bulk and junctional regions. *J*_c,bulk_ and *J*_c,junc_ are the sum of reaction rates and other nondiffusive fluxes of species *c* in the bulk and junctional spaces, respectively. The above relation only applies to species in a volumetric compartment (SR lumen, myoplasm, or T-tubule volume). Variables associated with the state of surface species (e.g., ion channels in the sarcolemma) are assumed to be non-diffusive, with the exception of membrane voltage, which diffuses rapidly between the sarcolemma and T-tubules (Equations (3) and (4)).

To calculate the volume and surface area of each compartment (sarcolemma, T-tubule, myoplasm, SR membrane, SR lumen, and mitochondrial volume), we approximate the muscle fiber as a cylinder of radius 20 µm, matching the size of mouse fast twitch fibers (Wallinga et al., 1999; Luff and Atwood, 1972). The total volume and sarcolemma surface area are given by the volume and surface area of a cylindrical cross-section. We assume the fiber is homogeneous along the longitudinal direction, such that the length of our cylindrical volume of interest is arbitrary. The subcellular volumes (myoplasm, mitochondria, and SR lumen) are then calculated from previously determined volume fractions (Senneff and Lowery, 2021; Rincón et al., 2021), whereas the T-tubule and SR surface area are given by the surface area to volume ratios measured in (Peachey, 1965) and (Mobley and Eisenberg, 1975). The junctional myoplasmic volume is defined as the region within Δ*d*_diff_ of the T-tubule surface, and, similarly, the junctional SR volume is defined as the volume within Δ*d*_diff_ of the junctional SR membrane. The interface between the junctional SR and T-tubule membrane is assumed to occupy about 50% of the T-tubule surface, in good agreement with the 67% occupancy measured in frog skeletal muscle (Mobley and Eisenberg, 1975). Finally, the surface area of the junction between T-tubules and the extracellular space (*SA*_EC,bulk-junc_)is computed assuming the volume fraction of T-tubules is constant throughout the fiber. All geometric parameters are summarized in Table A1 and expressions for individual subvolumes and surface areas are given in Table A2.

**Table A1:** Geometric parameters.

| Name | Description | Value | Reference |
| --- | --- | --- | --- |
| $R_{\text{fiber}}$ | Fiber radius | 20 $\mu\text{m}$ | (Wallinga et al., 1999) |
| $L_{\text{fiber}}$ | fiber length | 100 $\mu\text{m}$ | (arbitrary) |
| $\phi_{\text{myo}}$ | Myoplasm volume fraction | 0.947 | (Shorten et al., 2007) |
| $\phi_{\text{SR}}$ | SR volume fraction | 0.05 | (Shorten et al., 2007) |
| $\phi_{\text{TT}}$ | T-tubule volume fraction | 0.003 | (Mathias et al., 1977) |
| $\theta_{\text{TT}}$ | T-tubule volume-to-surface-area ratio | 0.01 $\mu\text{m}$ | (Peachey, 1965) |
| $\theta_{\text{SR}}$ | SR volume-to-surface-area ratio | 0.1 $\mu\text{m}$ | (Mobley and Eisenberg, 1975) |
| $\phi_{\text{SRM,TT-SR}}$ | area fraction of SRM at SR-TT junction | 0.5 | (Mobley and Eisenberg, 1975) |
| $\phi_{\text{Mito}}$ | Mitochondria volume fraction (per myoplasm) | 0.05 | (Chen et al., 2001) |

**Table A2:** Equations for sub-compartment volumes and surface areas.

| Major compartment | Subregion | Equation |
| --- | --- | --- |
| Extracellular | Total | $vol_{\text{EC,tot}} \rightarrow \infty$ |
| | Junctional | $vol_{\text{TT}} = \phi_{\text{TT}} \pi R_{\text{fiber}}^2 L_{\text{fiber}}$ |
| | Bulk | $vol_{\text{EC}} \rightarrow \infty$ |
| | Bulk-junc interface | $SA_{\text{EC,bulk-junc}} = \phi_{\text{TT}} 2\pi R_{\text{fiber}} L_{\text{fiber}}$ |
| Plasma membrane | Total | $SA_{\text{PM}} = SA_{\text{SL}} + SA_{\text{TT}}$ |
| | Junctional | $SA_{\text{TT}} = \frac{\phi_{\text{TT}} \pi R_{\text{fiber}}^2 L_{\text{fiber}}}{\theta_{\text{TT}}}$ |
| | Bulk | $SA_{\text{SL}} = 2\pi R_{\text{fiber}} L_{\text{fiber}}$ |
| Myoplasm | Total | $vol_{\text{myo}} = \phi_{\text{myo}} (1 - \phi_{\text{Mito}}) \pi R_{\text{fiber}}^2 L_{\text{fiber}}$ |
| | Junctional | $vol_{\text{myo,junc}} = SA_{\text{TT}} \Delta d_{\text{diff}} (1 - \phi_{\text{Mito}})$ |
| | Bulk | $vol_{\text{myo,bulk}} = vol_{\text{myo}} - vol_{\text{myo,junc}}$ |
| | Bulk-junc interface | $SA_{\text{myo,bulk-junc}} = SA_{\text{TT}} (1 - \phi_{\text{SRM,TT-SR}})$ |
| SR membrane | Total | $SA_{\text{SRM}} = \frac{\phi_{\text{SR}} \pi R_{\text{fiber}}^2 L_{\text{fiber}}}{\theta_{\text{SR}}}$ |
| | Junctional | $SA_{\text{SRM,junc}} = SA_{\text{TT}} \phi_{\text{SRM,TT-SR}}$ |
| | Bulk | $SA_{\text{SRM,bulk}} = SA_{\text{SRM}} - SA_{\text{SRM,junc}}$ |
| SR | Total | $vol_{\text{SR}} = \phi_{\text{SR}} \pi R_{\text{fiber}}^2 L_{\text{fiber}}$ |
| | Junctional | $vol_{\text{SR,junc}} = SA_{\text{SR,junc}} \Delta d_{\text{diff}}$ |
| | Bulk | $vol_{\text{SR,bulk}} = vol_{\text{SR}} - vol_{\text{SR,junc}}$ |
| | Bulk-junc interface | $SA_{\text{SR,bulk-junc}} = SA_{\text{SRM,junc}}$ |
| Mito matrix | Total | $vol_{\text{Mito}} = \phi_{\text{myo}} \phi_{\text{Mito}} \pi R_{\text{fiber}}^2 L_{\text{fiber}}$ |
| | Junctional | $vol_{\text{Mito,junc}} = SA_{\text{TT}} \Delta d_{\text{diff}} \phi_{\text{Mito}}$ |
| | Bulk | $vol_{\text{Mito,bulk}} = vol_{\text{Mito}} - vol_{\text{Mito,junc}}$ |
|  | Bulk-junc interface | 0 |

**Table A3:** Ionic fluxes summarized by region. References provided motivate the distribution of channels between junctional and bulk regions; in absence of additional evidence, we assume equal distribution across surface compartments.

| Flux | ions (stoich.*) | TT | SL | Junctional SRM | Bulk SRM | Reference |
| --- | --- | --- | --- | --- | --- | --- |
| $J_{Na}$ | $Na^+$ (+1) | 0.1 | 1 | 0 | 0 | (Cannon et al., 1993) |
| $J_{NaK}$ | $Na^+$ (-3), $K^+$ (+2) | 0.1 | 1 | 0 | 0 | (Wallinga et al., 1999) |
| $J_{NCX}$ | $Na^+$ (+3), $Ca^{2+}$ (-1) | 1 | 1 | 0 | 0 | This study |
| $J_{KIR}$ | $K^+$ (+1) | 1 | 1 | 0 | 0 | (Ashcroft et al., 1985) |
| $J_{KDR}$ | $K^+$ (-1) | 0.45 | 1 | 0 | 0 | (Wallinga et al., 1999) |
| $J_{Cl^-}$ | $Cl^-$ (+1) | 0.1 | 1 | 0 | 0 | (Wallinga et al., 1999) |
| $J_{Ca^{2+} leak,M}$ | $Ca^{2+}$ (+1) | 1 | 1 | 0 | 0 | This study |
| $J_{SOCE}$ | $Ca^{2+}$ (+1) | 1 | 0 | 0 | 0 | (Protasi et al., 2021) |
| $J_{PMCA}$ | $Ca^{2+}$ (-1) | 1 | 1 | 0 | 0 | This study |
| $J_{RyR}$ | $Ca^{2+}$ (+1) | 0 | 0 | 1 | 0 | (Senneff and Lowery, 2021) |
| $J_{SERCA}$ | $Ca^{2+}$ (-1) | 0 | 0 | 0.1 | 1 | (Scheibe et al., 2008) |
| $J_{Ca^{2+} leak,SR}$ | $Ca^{2+}$ (+1) | 0 | 0 | 1 | 1 | (Wallinga et al., 1999) |

\*Stoichiometric coefficient is positive for fluxes into the myoplasm and negative for fluxes out of the myoplasm.

**Table A4:**
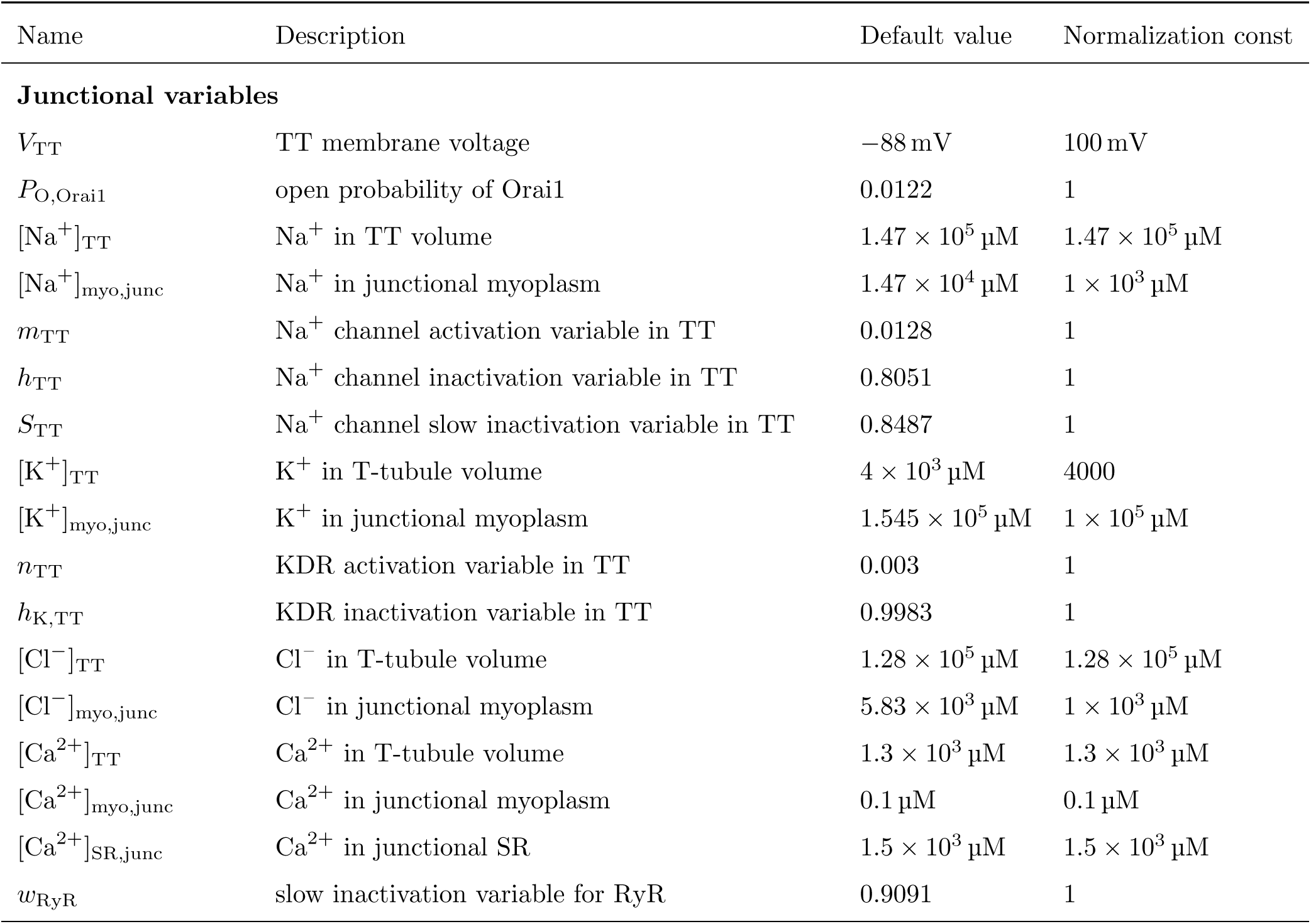
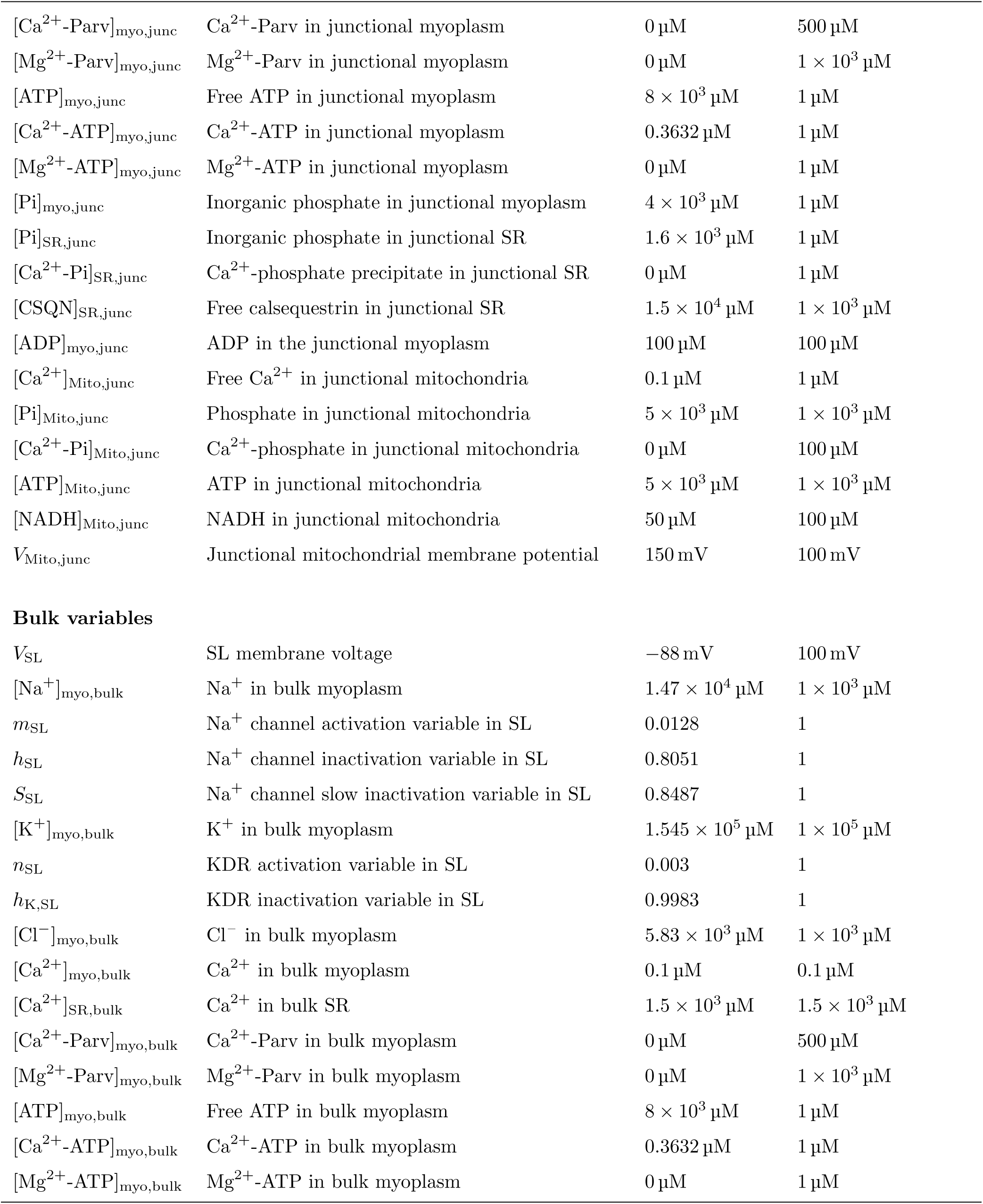
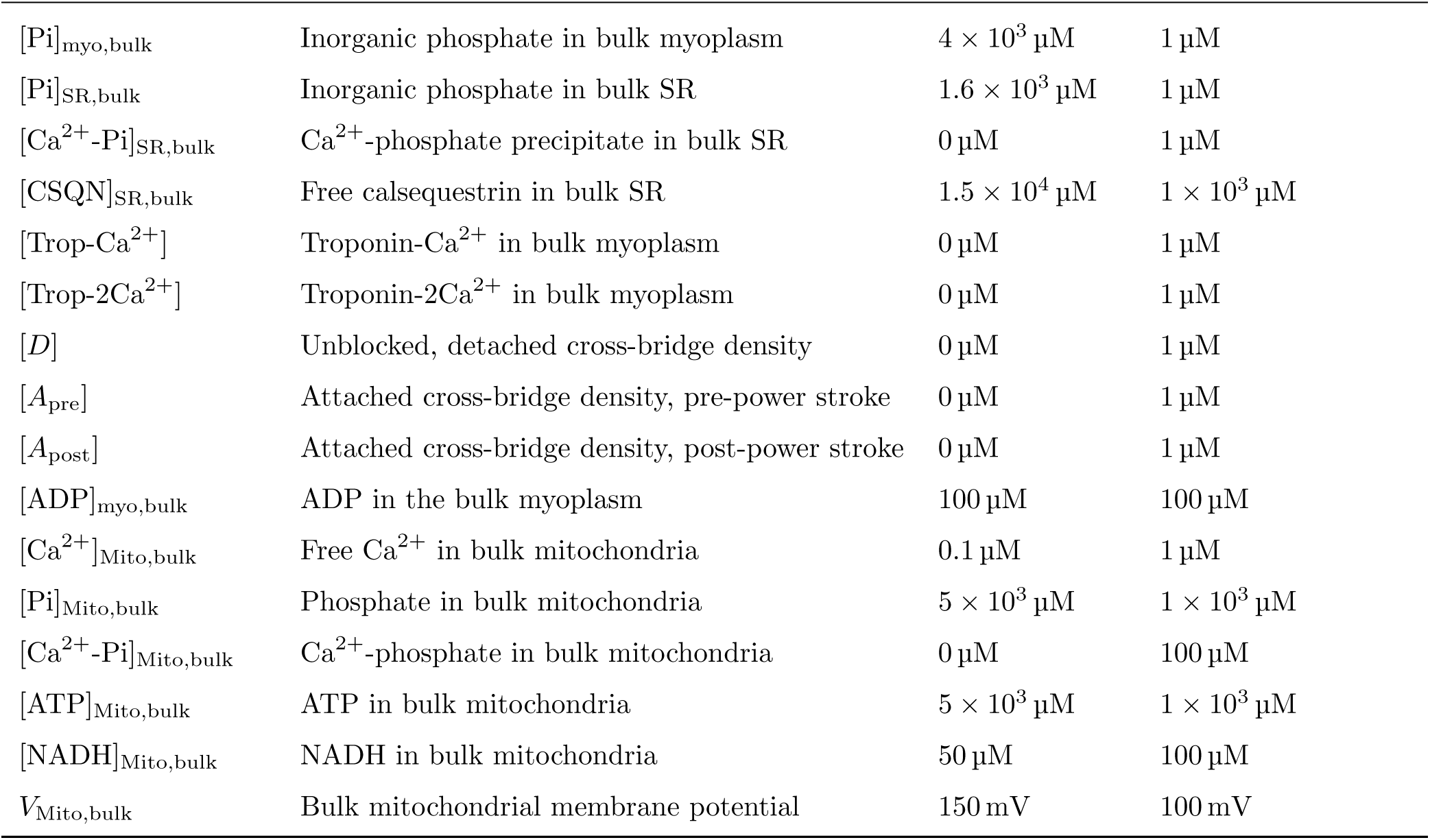
All state variables and their default values.

The equations used in our system of ordinary differential equations (ODEs) were adapted from different papers that incorporated Ca^2+^ signaling and action potential generation within mammalian muscle fibers. Reactions/fluxes are given for each module in Table A5. Throughout, whenever a given flux or variable is associated with both bulk and junctional regions, we denote the intracellular region with subscript i, the extracellular region with subscript o, and the associated region of the plasma membrane (either T-tubules or bulk sarcolemma) with subscript M. For instance, we write all Nernst potentials as:

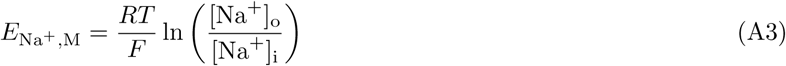

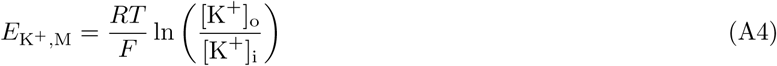

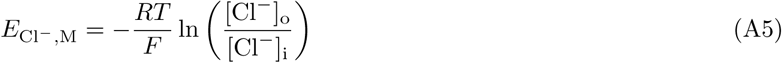

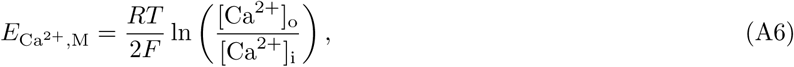

where [M,i,o] can either be [‘TT’,‘myo,junc’,‘TT volume’] or [‘SL’,‘myo,bulk’,‘EC’]. Similarly, ‘SR’ and ‘Mito’ are used to refer to the associated junctional compartment (‘SR,junc‘, Mito,junc‘) or bulk compartment (‘SR,bulk’, ‘Mito,junc’) where applicable. The resulting ODEs are summarized in Table A6 and all associated parameters provided in Table A7.

**Table A5:**
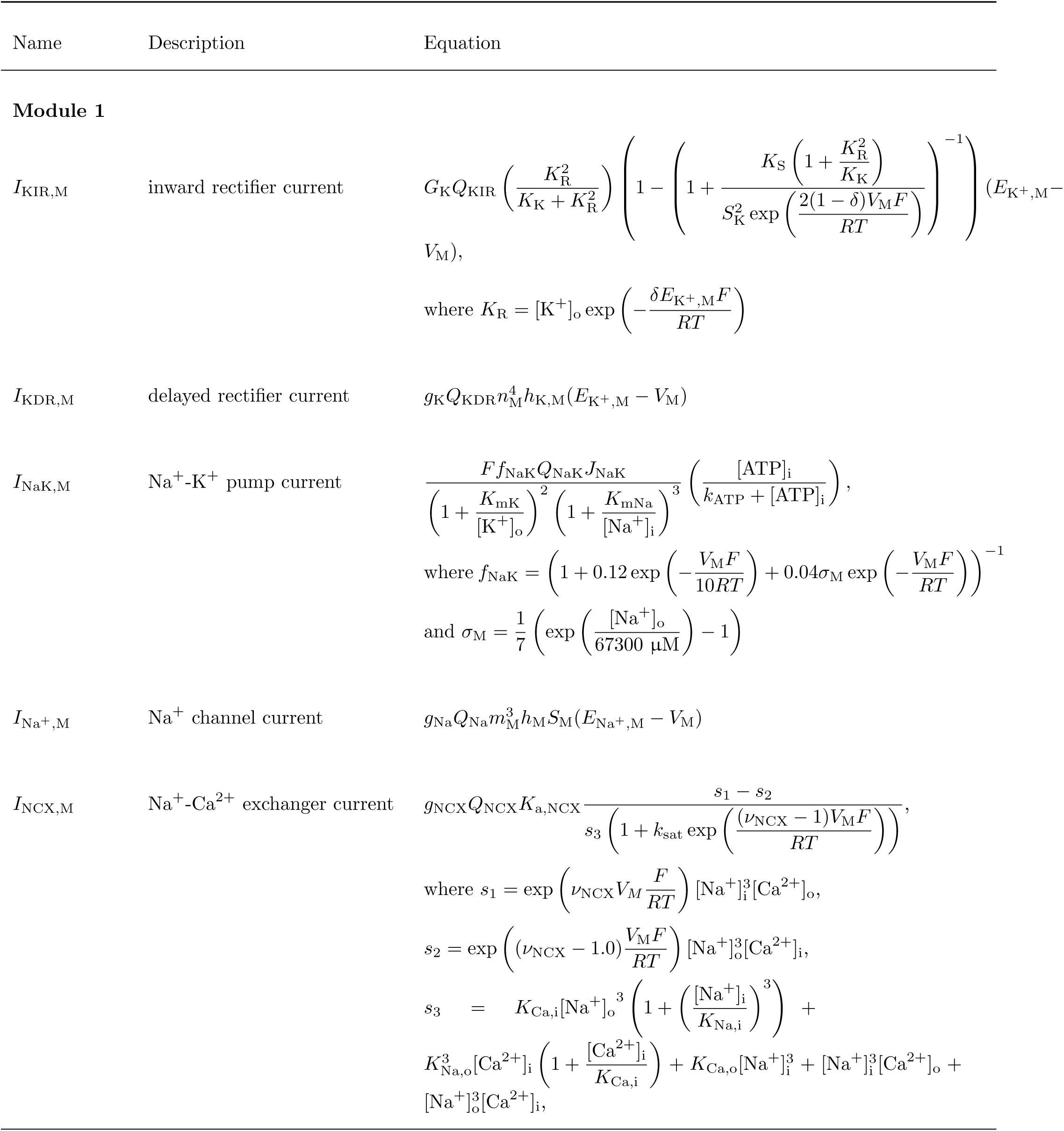
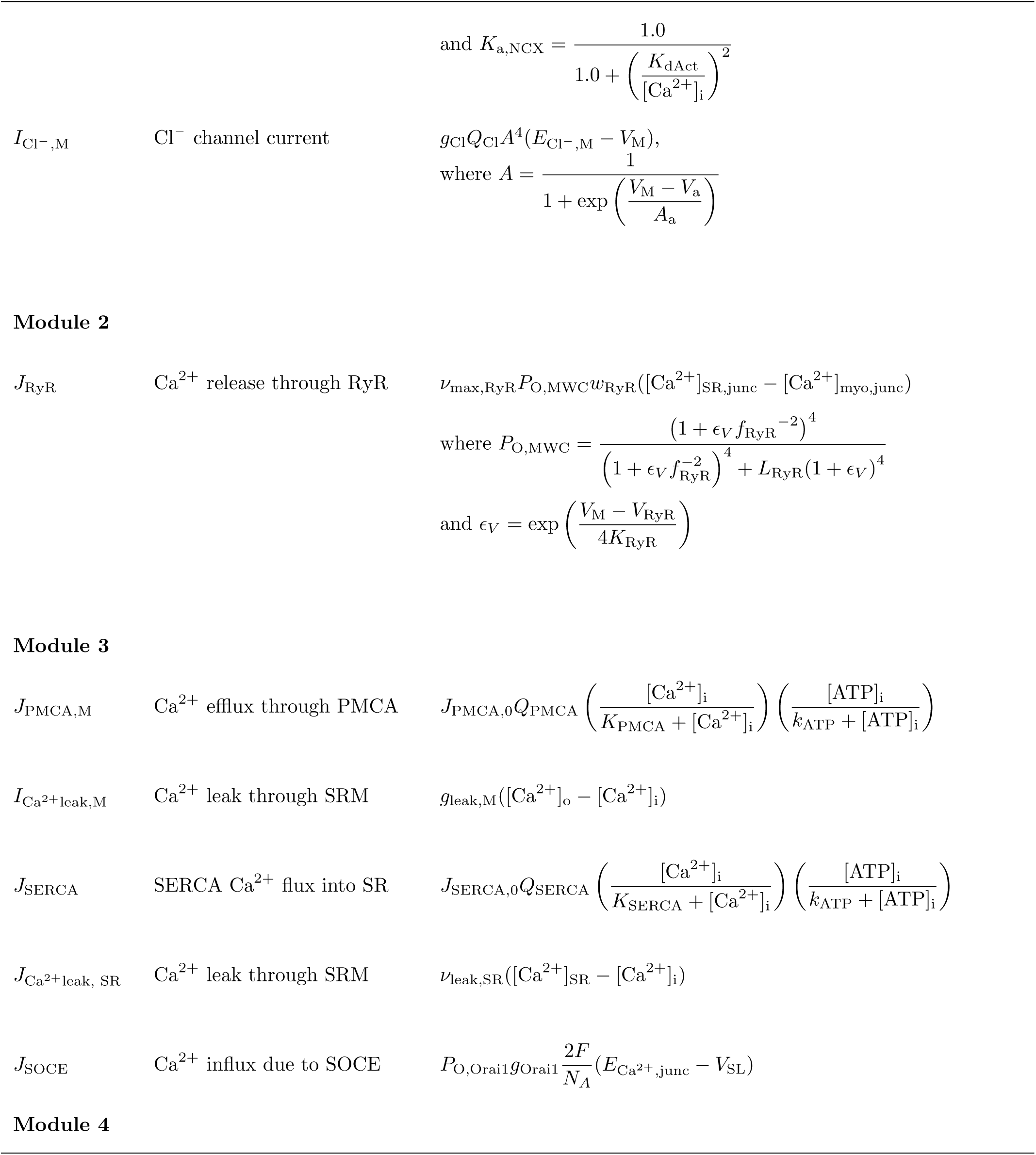
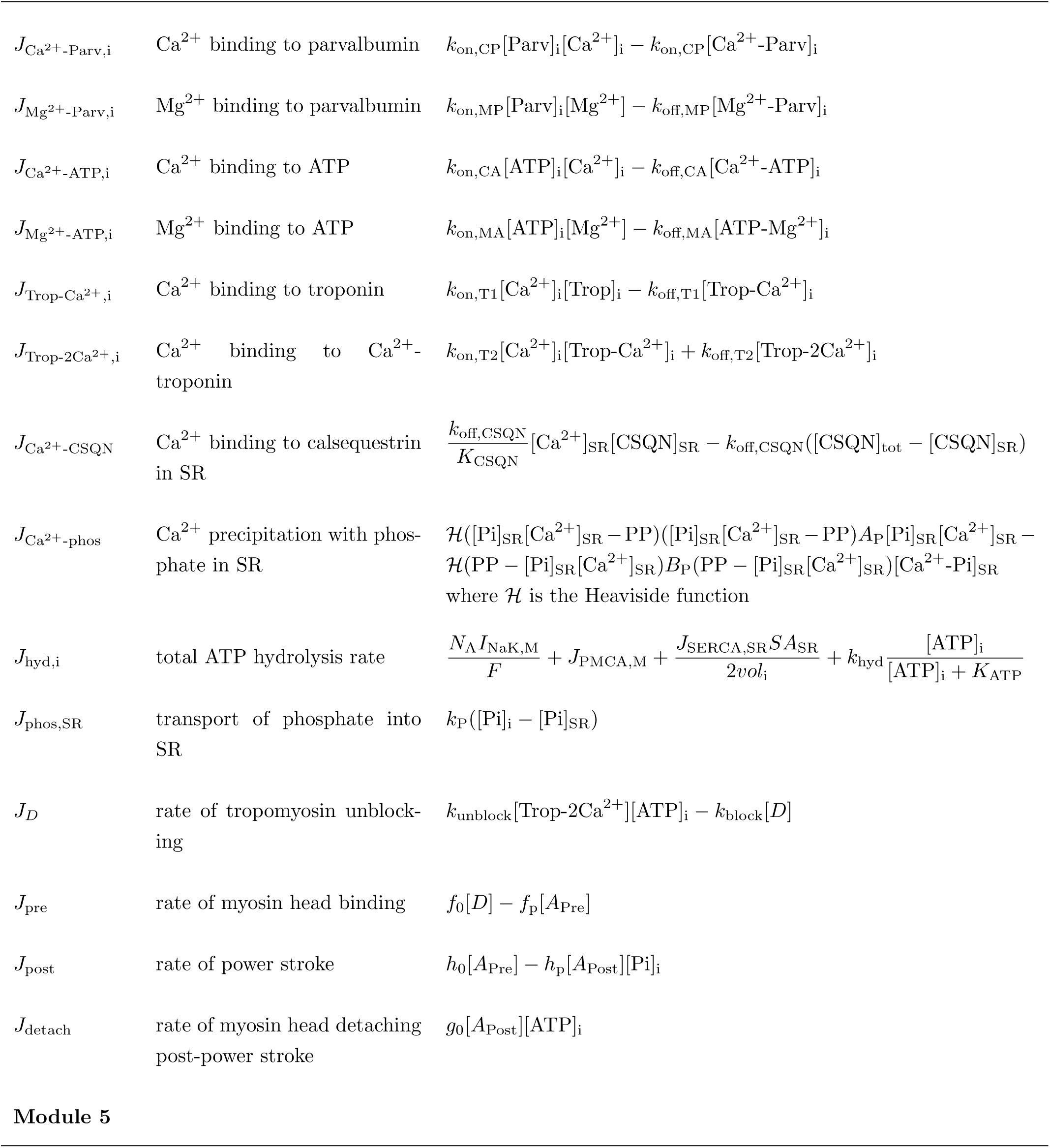
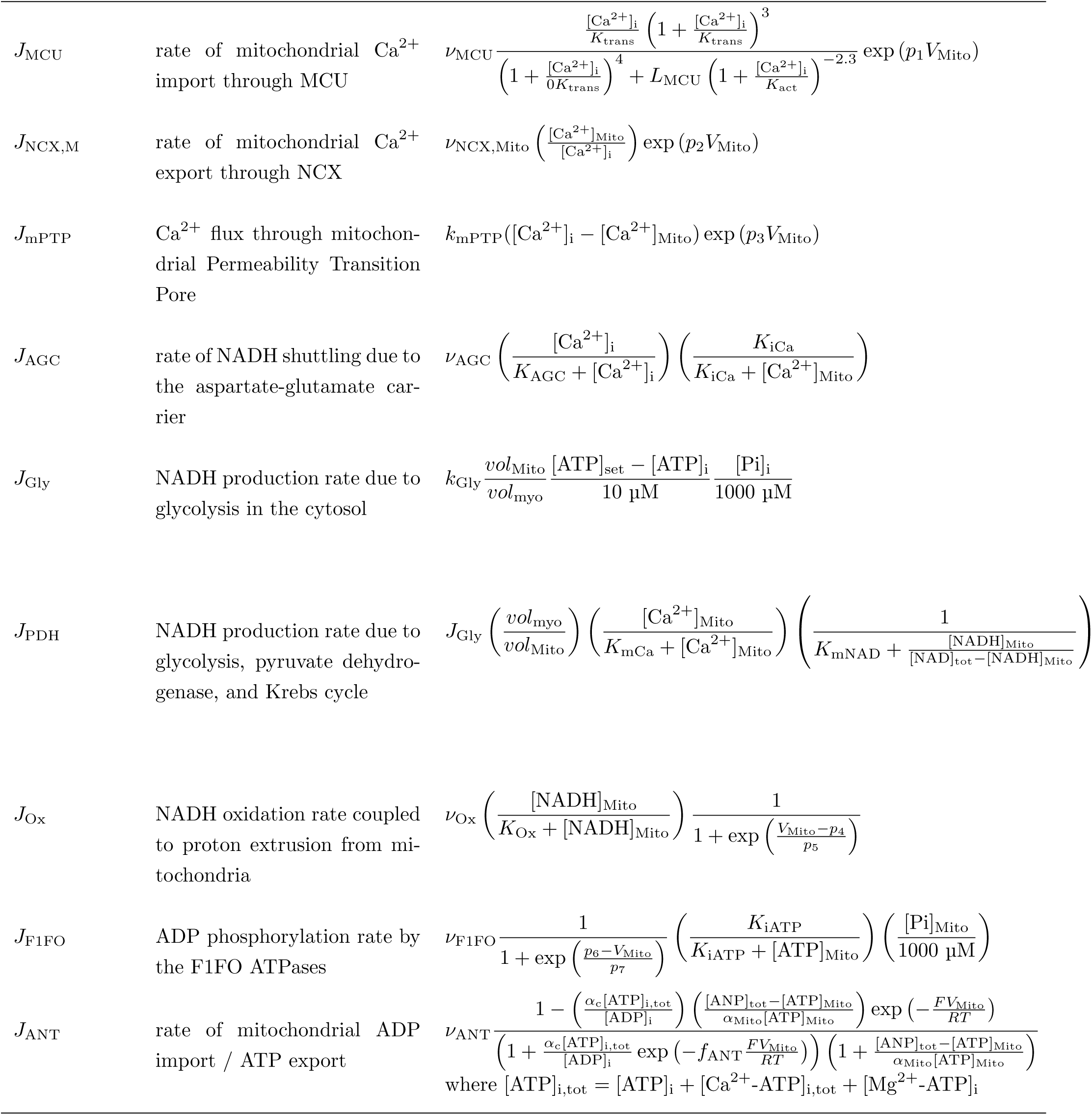
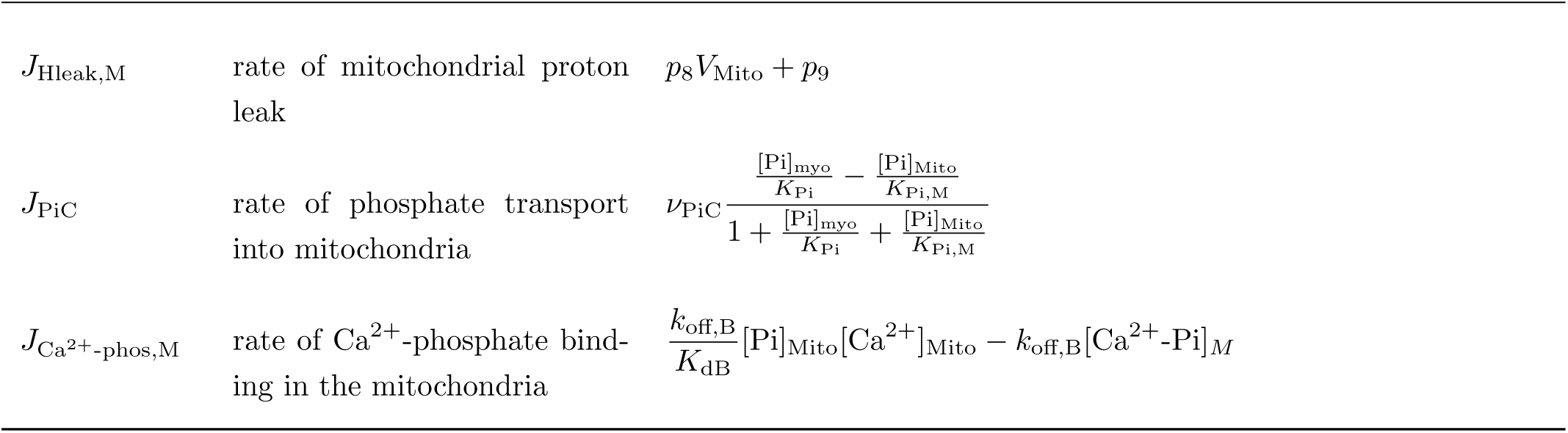
Reactions and fluxes.

**Table A6:** System of ODEs.

| Variable | Equation |
| --- | --- |
| **Junctional variables** |  |
| $\frac{d[\text{Na}^+]_{\text{TT}}}{dt}$ | $\frac{SA_{\text{TT}}\eta_{\text{conv},1}}{\text{vol}_{\text{TT}}F} (-0.1I_{\text{Na}^+,\text{TT}} + 3 \times 0.1I_{\text{NaK,TT}} - I_{\text{NCX,TT}}) + \frac{D_{\text{ion,EC}}SA_{\text{EC,bulk-junc}}}{\text{vol}_{\text{TT}}\Delta d_{\text{diff}}} ([\text{Na}^+]_{\text{EC}} - [\text{Na}^+]_{\text{TT}})$ |
| $\frac{d[\text{Na}^+]_{\text{myo,junc}}}{dt}$ | $\frac{SA_{\text{TT}}\eta_{\text{conv},1}}{\text{vol}_{\text{myo,junc}}F} (0.1I_{\text{Na}^+,\text{TT}} - 3 \times 0.1I_{\text{NaK,TT}} + I_{\text{NCX,TT}}) + \frac{D_{\text{ion}}SA_{\text{myo,bulk-junc}}}{\text{vol}_{\text{myo,junc}}\Delta d_{\text{diff}}} ([\text{Na}^+]_{\text{myo,bulk}} - [\text{Na}^+]_{\text{myo,junc}})$ |
| $\frac{dm_{\text{TT}}}{dt}$ | $\alpha_m(1 - m_{\text{TT}}) - \beta_m m_{\text{TT}},$<br>where $\beta_m = \beta_{m0} Q_{\beta_m} \exp\left(-\frac{V_{\text{TT}} - V_m}{K_{\beta_m}}\right)$ and $\alpha_m = \alpha_{m0} Q_{\alpha_m} \frac{V_{\text{TT}} - V_m}{1 - \exp\left(\frac{V_m - V_{\text{TT}}}{K_{\alpha_m}}\right)}$ |

**Table A6:** (continued) System of ODEs.
| Variable | Equation |
| --- | --- |
| $\frac{dh_{\text{TT}}}{dt}$ | $\alpha_h(1 - h_{\text{TT}}) - (\beta_h h_{\text{TT}}),$<br>where $\beta_h = Q_{\beta_h} \frac{\beta_{h0}}{1 + \exp\left(-\frac{V_{\text{TT}} - V_h}{K_{\beta_h}}\right)}$ and $\alpha_h = \alpha_{h0} Q_{\alpha_h} \exp\left(-\frac{V_{\text{TT}} - V_h}{K_{\alpha_h}}\right)$ |
| $\frac{dS_{\text{TT}}}{dt}$ | $\frac{S_{\infty} - S_{\text{TT}}}{\tau_S},$<br>where $S_{\infty} = \frac{1}{1 + \exp\left(\frac{V_{\text{TT}} - V_{S\infty}}{A_{S\infty}}\right)}$ and $\tau_S = \frac{60}{0.2 + 5.65 \left(\frac{V_{\text{TT}} + V_{\tau}}{100}\right)^2}$ |
| $\frac{d[\text{K}^+]_{\text{TT}}}{dt}$ | $\frac{SA_{\text{TT}}\eta_{\text{conv},1}}{\text{vol}_{\text{TT}}F}(-I_{\text{KIR,TT}} + 0.45I_{\text{KDR,TT}} - 2 \times 0.1I_{\text{NaK,TT}}) + \frac{D_{\text{ion,EC}}SA_{\text{EC,bulk-junc}}}{\text{vol}_{\text{TT}}\Delta d_{\text{diff}}}([\text{K}^+]_{\text{EC}} - [\text{K}^+]_{\text{TT}})$ |
| $\frac{d[\text{K}^+]_{\text{myo,junc}}}{dt}$ | $\frac{SA_{\text{TT}}\eta_{\text{conv},1}}{\text{vol}_{\text{myo,junc}}F}(I_{\text{KIR,TT}} - 0.45I_{\text{KDR,TT}} + 2 \times 0.1I_{\text{NaK,TT}}) + \frac{D_{\text{ion}}SA_{\text{myo,bulk-junc}}}{\text{vol}_{\text{myo,junc}}\Delta d_{\text{diff}}}([\text{K}^+]_{\text{myo,bulk}} - [\text{K}^+]_{\text{myo,junc}})$ |
| $\frac{dn_{\text{TT}}}{dt}$ | $\alpha_n(1 - n_{\text{TT}}) - \beta_n n_{\text{TT}},$<br>where $\beta_n = \beta_{n0} Q_{\beta_n} \exp\left(-\frac{V_{\text{TT}} - V_n}{K_{\beta_n}}\right)$ and $\alpha_n = \alpha_{n0} Q_{\alpha_n} \frac{V_{\text{TT}} - V_n}{1 - \exp\left(\frac{V_n - V_{\text{TT}}}{K_{\alpha_n}}\right)}$ |
| $\frac{dh_{\text{K,TT}}}{dt}$ | $\frac{h_{\text{K}\infty} - h_{\text{K,TT}}}{\tau_{h_{\text{K}}}},$<br>where $\tau_{h_{\text{K}}} = \exp\left(-\frac{V_{\text{TT}} + 40}{25.75}\right)$ and $h_{\text{K}\infty} = \left(1 + \exp\left(\frac{V_{\text{TT}} - V_{h_{\text{K}\infty}}}{A_{h_{\text{K}\infty}}}\right)\right)^{-1}$ |
| $\frac{d[\text{Cl}^-]_{\text{TT}}}{dt}$ | $-\frac{SA_{\text{TT}}\eta_{\text{conv},1}}{\text{vol}_{\text{TT}}F}(-0.1I_{\text{Cl}^-,\text{TT}}) + \frac{D_{\text{ion,EC}}SA_{\text{EC,bulk-junc}}}{\text{vol}_{\text{TT}}\Delta d_{\text{diff}}}([\text{Cl}^-]_{\text{EC}} - [\text{Cl}^-]_{\text{TT}})$ |

**Table A6:** (continued) System of ODEs.
| Variable | Equation |
| --- | --- |
| $\frac{d[\text{Cl}^-]_{\text{myo,junc}}}{dt}$ | $-\frac{SA_{\text{TT}}\eta_{\text{conv},1}}{vol_{\text{myo,junc}}F}(0.1I_{\text{Cl}^-,\text{TT}}) + \frac{D_{\text{ion}}SA_{\text{myo,bulk-junc}}}{vol_{\text{myo,junc}}\Delta d_{\text{diff}}}([\text{Cl}^-]_{\text{myo,bulk}} - [\text{Cl}^-]_{\text{myo,junc}})$ |
| $\frac{d[\text{Ca}^{2+}]_{\text{TT}}}{dt}$ | $\frac{SA_{\text{TT}}}{vol_{\text{TT}}\eta_{\text{conv},2}}(-J_{\text{Ca}^{2+}\text{ leak,TT}} - J_{\text{SOCE,TT}} + \frac{N_{\text{A}}}{F}I_{\text{NCX,TT}}) + J_{\text{PMCA,TT}}\frac{vol_{\text{myo,junc}}}{vol_{\text{TT}}} + \frac{D_{\text{ion,EC}}SA_{\text{EC,bulk-junc}}}{vol_{\text{TT}}\Delta d_{\text{diff}}}([\text{Ca}^{2+}]_{\text{EC}} - [\text{Ca}^{2+}]_{\text{TT}})$ |
| $\frac{d[\text{Ca}^{2+}]_{\text{myo,junc}}}{dt}$ | $\frac{SA_{\text{TT}}}{vol_{\text{myo,junc}}\eta_{\text{conv},2}}(J_{\text{Ca}^{2+}\text{ leak,TT}} + J_{\text{SOCE,TT}} - \frac{N_{\text{A}}}{F}I_{\text{NCX,TT}}) - J_{\text{PMCA,TT}} + \frac{SA_{\text{SRM,junc}}}{vol_{\text{SR,junc}}}(J_{\text{RyR,junc}} + J_{\text{Ca}^{2+}\text{ leak SR,junc}} - 0.1J_{\text{SERCA,junc}}) - J_{\text{Ca}^{2+}\text{-Parv,junc}} - J_{\text{Ca}^{2+}\text{-ATP,junc}} - \frac{vol_{\text{Mito,junc}}}{vol_{\text{myo,junc}}}(J_{\text{MCU,junc}} - J_{\text{NCX,Mito,junc}} + J_{\text{mPTP,junc}}) + \frac{D_{\text{ion}}SA_{\text{myo,bulk-junc}}}{vol_{\text{myo,junc}}\Delta d_{\text{diff}}}([\text{Ca}^{2+}]_{\text{myo,bulk}} - [\text{Ca}^{2+}]_{\text{myo,junc}})$ |
| $\frac{d[\text{Ca}^{2+}]_{\text{SR,junc}}}{dt}$ | $\frac{SA_{\text{SRM,junc}}}{vol_{\text{SR,junc}}}(-J_{\text{Ca}^{2+}\text{ leak SR,junc}} - J_{\text{RyR,junc}} + 0.1J_{\text{SERCA,junc}}) - J_{\text{Ca}^{2+}\text{-CSQN,junc}} + \frac{D_{\text{ion}}SA_{\text{SR,bulk-junc}}}{vol_{\text{SR,junc}}\Delta d_{\text{diff}}}([\text{Ca}^{2+}]_{\text{SR,bulk}} - [\text{Ca}^{2+}]_{\text{SR,junc}})$ |
| $\frac{dw_{\text{RyR}}}{dt}$ | $\alpha_w \left( 1 - w_{\text{RyR}} \left( 1 + \frac{[\text{Ca}^{2+}]_{\text{myo,junc}}}{K_w} \right) \right)$ |
| $\frac{d[\text{Ca}^{2+}\text{-Parv}]_{\text{myo,junc}}}{dt}$ | $J_{\text{Ca}^{2+}\text{-Parv,junc}} + \frac{D_{\text{Parv}}SA_{\text{myo,bulk-junc}}}{vol_{\text{myo,junc}}\Delta d_{\text{diff}}}([\text{Ca}^{2+}\text{-Parv}]_{\text{myo,bulk}} - [\text{Ca}^{2+}\text{-Parv}]_{\text{myo,junc}})$ |
| $\frac{d[\text{Mg}^{2+}\text{-Parv}]_{\text{myo,junc}}}{dt}$ | $J_{\text{Mg}^{2+}\text{-Parv,junc}} + \frac{D_{\text{Parv}}SA_{\text{myo,bulk-junc}}}{vol_{\text{myo,junc}}\Delta d_{\text{diff}}}([\text{Mg}^{2+}\text{-Parv}]_{\text{myo,bulk}} - [\text{Mg}^{2+}\text{-Parv}]_{\text{myo,junc}})$ |

**Table A6:** (continued) System of ODEs.
| Variable | Equation |
| --- | --- |
| $\frac{d[\text{ATP}]_{\text{myo,junc}}}{dt}$ | $-J_{\text{Ca}^{2+}\text{-ATP,junc}} - J_{\text{Mg}^{2+}\text{-ATP,junc}} - J_{\text{hydro,junc}} + J_{\text{Gly,junc}}N_{\text{ATP-per-NADH}} + J_{\text{ANT,junc}}\frac{vol_{\text{Mito,junc}}}{vol_{\text{myo,junc}}} + \frac{D_{\text{ATP}}SA_{\text{myo,bulk-junc}}}{vol_{\text{myo,junc}}\Delta d_{\text{diff}}}([\text{ATP}]_{\text{myo,bulk}} - [\text{ATP}]_{\text{myo,junc}})$ |
| $\frac{d[\text{Ca}^{2+}\text{-ATP}]_{\text{myo,junc}}}{dt}$ | $J_{\text{Ca}^{2+}\text{-ATP,junc}} + \frac{D_{\text{ATP}}SA_{\text{myo,bulk-junc}}}{vol_{\text{myo,junc}}\Delta d_{\text{diff}}}([\text{Ca}^{2+}\text{-ATP}]_{\text{myo,bulk}} - [\text{Ca}^{2+}\text{-ATP}]_{\text{myo,junc}})$ |
| $\frac{d[\text{Mg}^{2+}\text{-ATP}]_{\text{myo,junc}}}{dt}$ | $J_{\text{Mg}^{2+}\text{-ATP,junc}} + \frac{D_{\text{ATP}}SA_{\text{myo,bulk-junc}}}{vol_{\text{myo,junc}}\Delta d_{\text{diff}}}([\text{Mg}^{2+}\text{-ATP}]_{\text{myo,bulk}} - [\text{Mg}^{2+}\text{-ATP}]_{\text{myo,junc}})$ |
| $\frac{d[\text{Pi}]_{\text{myo,junc}}}{dt}$ | $J_{\text{hydro,junc}} - J_{\text{Gly,junc}}N_{\text{ATP-per-NADH}} - J_{\text{phos,SR,junc}} - J_{\text{PiC,junc}}\frac{vol_{\text{Mito,junc}}}{vol_{\text{myo,junc}}} + \frac{D_{\text{Pi}}SA_{\text{myo,bulk-junc}}}{vol_{\text{myo,junc}}\Delta d_{\text{diff}}}([\text{Pi}]_{\text{myo,bulk}} - [\text{Pi}]_{\text{myo,junc}})$ |
| $\frac{d[\text{Pi}]_{\text{SR,junc}}}{dt}$ | $J_{\text{phos,SR,junc}} - J_{\text{Ca}^{2+}\text{-phos,junc}} + \frac{D_{\text{Pi}}SA_{\text{SR,bulk-junc}}}{vol_{\text{SR,junc}}\Delta d_{\text{diff}}}([\text{Pi}]_{\text{SR,bulk}} - [\text{Pi}]_{\text{SR,junc}})$ |
| $\frac{d[\text{Ca}^{2+}\text{-Pi}]_{\text{SR,junc}}}{dt}$ | $J_{\text{Ca}^{2+}\text{-phos,junc}} + \frac{D_{\text{Pi}}SA_{\text{SR,bulk-junc}}}{vol_{\text{SR,junc}}\Delta d_{\text{diff}}}([\text{Ca}^{2+}\text{-Pi}]_{\text{SR,bulk}} - [\text{Ca}^{2+}\text{-Pi}]_{\text{SR,junc}})$ |
| $\frac{d[\text{CSQN}]_{\text{SR,junc}}}{dt}$ | $-J_{\text{Ca}^{2+}\text{-CSQN,junc}} + \frac{D_{\text{CSQN}}SA_{\text{SR,bulk-junc}}}{vol_{\text{SR,junc}}\Delta d_{\text{diff}}}([\text{CSQN}]_{\text{SR,bulk}} - [\text{CSQN}]_{\text{SR,junc}})$ |
| $\frac{d[\text{ADP}]_{\text{myo,junc}}}{dt}$ | $J_{\text{hydro,junc}} - J_{\text{Gly,junc}}N_{\text{ATP-per-NADH}} - J_{\text{ANT,junc}}\frac{vol_{\text{Mito,junc}}}{vol_{\text{myo,junc}}} + \frac{D_{\text{ATP}}SA_{\text{myo,bulk-junc}}}{vol_{\text{myo,junc}}\Delta d_{\text{diff}}}([\text{ADP}]_{\text{myo,bulk}} - [\text{ADP}]_{\text{myo,junc}})$ |

**Table A6:** (continued) System of ODEs.
| Variable | Equation |
| --- | --- |
| $\frac{d[\text{Ca}^{2+}]_{\text{Mito,junc}}}{dt}$ | $b_{\text{M}} (J_{\text{MCU,junc}} - J_{\text{NCX,Mito,junc}} + J_{\text{mPTP,junc}} - J_{\text{Ca}^{2+}\text{-phos,Mito,junc}})$ |
| $\frac{d[\text{Pi}]_{\text{Mito,junc}}}{dt}$ | $J_{\text{PiC,junc}} - J_{\text{Ca}^{2+}\text{-phos,Mito,junc}}$ |
| $\frac{d[\text{Ca}^{2+}\text{-Pi}]_{\text{Mito,junc}}}{dt}$ | $J_{\text{Ca}^{2+}\text{-phos,Mito,junc}}$ |
| $\frac{d[\text{ATP}]_{\text{Mito,junc}}}{dt}$ | $J_{\text{F1FO,junc}} - J_{\text{ANT,junc}}$ |
| $\frac{d[\text{NADH}]_{\text{Mito,junc}}}{dt}$ | $J_{\text{PDH,junc}} + J_{\text{AGC,junc}} - J_{\text{Ox,junc}}$ |
| $\frac{dV_{\text{Mito,junc}}}{dt}$ | $(a_1 J_{\text{Ox,junc}} - a_2 J_{\text{F1FO,junc}} - J_{\text{Hleak,Mito,junc}} - J_{\text{NCX,Mito,junc}} - 2J_{\text{MCU,junc}} - 2J_{\text{mPTP,junc}} - J_{\text{ANT,junc}} - J_{\text{AGC,junc}})/C_{\text{Eff,Mito}}$ |
| **Bulk variables** |  |
| $\frac{d[\text{Na}^+]_{\text{myo,bulk}}}{dt}$ | $\frac{SA_{\text{SL}}\eta_{\text{conv},1}}{vol_{\text{myo,bulk}}F}(I_{\text{Na}^+,\text{SL}} - 3I_{\text{NaK,SL}} + I_{\text{NCX,SL}}) + \frac{D_{\text{ion}}SA_{\text{myo,bulk-junc}}}{vol_{\text{myo,bulk}}\Delta d_{\text{diff}}}([\text{Na}^+]_{\text{myo,junc}} - [\text{Na}^+]_{\text{myo,bulk}})$ |
| $\frac{dm_{\text{SL}}}{dt}$ | $\frac{SA_{\text{SL}}\eta_{\text{conv},1}}{vol_{\text{myo,bulk}}F}(I_{\text{KIR,SL}} - I_{\text{KDR,SL}} + 2I_{\text{NaK,TT}}) + \frac{D_{\text{ion}}SA_{\text{myo,bulk-junc}}}{vol_{\text{myo,bulk}}\Delta d_{\text{diff}}}([\text{K}^+]_{\text{myo,junc}} - [\text{K}^+]_{\text{myo,bulk}})$ |

**Table A6:** (continued) System of ODEs.
| Variable | Equation |
| --- | --- |
| $\frac{dh_{\text{SL}}}{dt}$ | $\alpha_h(1 - h_{\text{SL}}) - (\beta_h h_{\text{SL}}),$<br>where $\beta_h = Q_{\beta_h} \frac{\beta_{h0}}{1 + \exp\left(-\frac{V_{\text{SL}} - V_h}{K_{\beta_h}}\right)}$ and $\alpha_h = \alpha_{h0} Q_{\alpha_h} \exp\left(-\frac{V_{\text{SL}} - V_h}{K_{\alpha_h}}\right)$ |
| $\frac{dS_{\text{SL}}}{dt}$ | $\frac{S_{\infty} - S_{\text{SL}}}{\tau_S},$<br>where $S_{\infty} = \frac{1}{1 + \exp\left(\frac{V_{\text{SL}} - V_{S\infty}}{A_{S\infty}}\right)}$ and $\tau_S = \frac{60}{0.2 + 5.65 \left(\frac{V_{\text{SL}} + V_{\tau}}{100}\right)^2}$ |
| $\frac{d[\text{K}^+]_{\text{myo,bulk}}}{dt}$ | $\frac{SA_{\text{SL}}\eta_{\text{conv},1}}{vol_{\text{myo,bulk}}F}(I_{\text{KIR,SL}} - I_{\text{KDR,SL}} + 2I_{\text{NaK,SL}}) + \frac{D_{\text{ion}}SA_{\text{myo,bulk-junc}}}{vol_{\text{myo,bulk}}\Delta d_{\text{diff}}}([\text{K}^+]_{\text{myo,junc}} - [\text{K}^+]_{\text{myo,bulk}})$ |
| $\frac{dn_{\text{SL}}}{dt}$ | $\alpha_n(1 - n_{\text{SL}}) - \beta_n n_{\text{SL}},$<br>where $\beta_n = \beta_{n0} Q_{\beta_n} \exp\left(-\frac{V_{\text{SL}} - V_n}{K_{\beta_n}}\right)$ and $\alpha_n = \alpha_{n0} Q_{\alpha_n} \frac{V_{\text{SL}} - V_n}{1 - \exp\left(\frac{V_n - V_{\text{SL}}}{K_{\alpha_n}}\right)}$ |
| $\frac{dh_{\text{K,SL}}}{dt}$ | $\frac{h_{\text{K}\infty} - h_{\text{K,SL}}}{\tau_{h_{\text{K}}}},$<br>where $\tau_{h_{\text{K}}} = \exp\left(-\frac{V_{\text{SL}} + 40}{25.75}\right)$ and $h_{\text{K}\infty} = \left(1 + \exp\left(\frac{V_{\text{SL}} - V_{h_{\text{K}\infty}}}{A_{h_{\text{K}\infty}}}\right)\right)^{-1}$ |
| $\frac{d[\text{Cl}^-]_{\text{myo,bulk}}}{dt}$ | $-\frac{SA_{\text{SL}}\eta_{\text{conv},1}}{vol_{\text{myo,bulk}}F}(I_{\text{Cl}^-, \text{SL}}) + \frac{D_{\text{ion}}SA_{\text{myo,bulk-junc}}}{vol_{\text{myo,bulk}}\Delta d_{\text{diff}}}([\text{Cl}^-]_{\text{myo,junc}} - [\text{Cl}^-]_{\text{myo,bulk}})$ |

**Table A6:** (continued) System of ODEs.
| Variable | Equation |
| --- | --- |
| $\frac{d[\text{Ca}^{2+}]_{\text{myo,bulk}}}{dt}$ | $\frac{SA_{\text{SL}}}{vol_{\text{myo,bulk}}\eta_{\text{conv},2}}(J_{\text{Ca}^{2+} \text{ leak,SL}} - \frac{N_{\text{A}}}{F}I_{\text{NCX,SL}}) - J_{\text{PMCA,SL}} +$ $\frac{SA_{\text{SRM,bulk}}}{vol_{\text{myo,bulk}}}(J_{\text{Ca}^{2+} \text{ leak SR,bulk}} - J_{\text{SERCA,bulk}}) - J_{\text{Ca}^{2+}\text{-Parv,bulk}} - J_{\text{Ca}^{2+}\text{-ATP,bulk}} -$ $\frac{vol_{\text{Mito,bulk}}}{vol_{\text{myo,bulk}}}(J_{\text{MCU,bulk}} - J_{\text{NCX,bulk}} + J_{\text{mPTP,bulk}}) + \frac{D_{\text{ion}}SA_{\text{myo,bulk-junc}}}{vol_{\text{myo,bulk}}\Delta d_{\text{diff}}}([\text{Ca}^{2+}]_{\text{myo,junc}} -$ $[\text{Ca}^{2+}]_{\text{myo,bulk}})$ |
| $\frac{d[\text{Ca}^{2+}]_{\text{SR,bulk}}}{dt}$ | $\frac{SA_{\text{SRM,bulk}}}{vol_{\text{SR,bulk}}}(-J_{\text{Ca}^{2+} \text{ leak SR,bulk}} + J_{\text{SERCA,bulk}}) - J_{\text{Ca}^{2+}\text{-CSQN,bulk}} +$ $\frac{D_{\text{ion}}SA_{\text{SR,bulk-junc}}}{vol_{\text{SR,bulk}}\Delta d_{\text{diff}}}([\text{Ca}^{2+}]_{\text{SR,junc}} - [\text{Ca}^{2+}]_{\text{SR,bulk}})$ |
| $\frac{d[\text{Ca}^{2+}\text{-Parv}]_{\text{myo,bulk}}}{dt}$ | $J_{\text{Ca}^{2+}\text{-Parv,bulk}} + \frac{D_{\text{Parv}}SA_{\text{myo,bulk-junc}}}{vol_{\text{myo,bulk}}\Delta d_{\text{diff}}}([\text{Ca}^{2+}\text{-Parv}]_{\text{myo,junc}} - [\text{Ca}^{2+}\text{-Parv}]_{\text{myo,bulk}})$ |
| $\frac{d[\text{Mg}^{2+}\text{-Parv}]_{\text{myo,bulk}}}{dt}$ | $J_{\text{Mg}^{2+}\text{-Parv,bulk}} + \frac{D_{\text{Parv}}SA_{\text{myo,bulk-junc}}}{vol_{\text{myo,bulk}}\Delta d_{\text{diff}}}([\text{Mg}^{2+}\text{-Parv}]_{\text{myo,junc}} - [\text{Mg}^{2+}\text{-Parv}]_{\text{myo,bulk}})$ |
| $\frac{d[\text{ATP}]_{\text{myo,bulk}}}{dt}$ | $-J_{\text{Ca}^{2+}\text{-ATP,bulk}} - J_{\text{Mg}^{2+}\text{-ATP,bulk}} - J_{\text{hydro,bulk}} + J_{\text{Gly,bulk}}N_{\text{ATP-per-NADH}} +$ $J_{\text{ANT,bulk}}\frac{vol_{\text{Mito,bulk}}}{vol_{\text{myo,bulk}}} - J_{\text{D}} - J_{\text{detach}} + \frac{D_{\text{ATP}}SA_{\text{myo,bulk-junc}}}{vol_{\text{myo,bulk}}\Delta d_{\text{diff}}}([\text{ATP}]_{\text{myo,junc}} - [\text{ATP}]_{\text{myo,bulk}})$ |
| $\frac{d[\text{Ca}^{2+}\text{-ATP}]_{\text{myo,bulk}}}{dt}$ | $J_{\text{Ca}^{2+}\text{-ATP,bulk}} + \frac{D_{\text{ATP}}SA_{\text{myo,bulk-junc}}}{vol_{\text{myo,bulk}}\Delta d_{\text{diff}}}([\text{Ca}^{2+}\text{-ATP}]_{\text{myo,junc}} - [\text{Ca}^{2+}\text{-ATP}]_{\text{myo,bulk}})$ |
| $\frac{d[\text{Mg}^{2+}\text{-ATP}]_{\text{myo,bulk}}}{dt}$ | $J_{\text{Mg}^{2+}\text{-ATP,bulk}} + \frac{D_{\text{ATP}}SA_{\text{myo,bulk-junc}}}{vol_{\text{myo,bulk}}\Delta d_{\text{diff}}}([\text{Mg}^{2+}\text{-ATP}]_{\text{myo,junc}} - [\text{Mg}^{2+}\text{-ATP}]_{\text{myo,bulk}})$ |

**Table A6:** (continued) System of ODEs.
| Variable | Equation |
| --- | --- |
| $\frac{d[\text{Pi}]_{\text{myo,bulk}}}{dt}$ | $J_{\text{hydro,bulk}} - J_{\text{phos,SR,bulk}} + J_{\text{post}} - J_{\text{PiC,bulk}} \frac{vol_{\text{Mito,bulk}}}{vol_{\text{myo,bulk}}} + \frac{D_{\text{Pi}} S A_{\text{myo,bulk-junc}}}{vol_{\text{myo,bulk}} \Delta d_{\text{diff}}} ([\text{Pi}]_{\text{myo,junc}} - [\text{Pi}]_{\text{myo,bulk}})$ |
| $\frac{d[\text{Pi}]_{\text{SR,bulk}}}{dt}$ | $J_{\text{phos,SR,bulk}} - J_{\text{Ca}^{2+}\text{-phos,bulk}} + \frac{D_{\text{Pi}} S A_{\text{SR,bulk-junc}}}{vol_{\text{SR,bulk}} \Delta d_{\text{diff}}} ([\text{Pi}]_{\text{SR,junc}} - [\text{Pi}]_{\text{SR,bulk}})$ |
| $\frac{d[\text{Ca}^{2+}\text{-Pi}]_{\text{SR,bulk}}}{dt}$ | $J_{\text{Ca}^{2+}\text{-phos,bulk}} + \frac{D_{\text{Pi}} S A_{\text{SR,bulk-junc}}}{vol_{\text{SR,bulk}} \Delta d_{\text{diff}}} ([\text{Ca}^{2+}\text{-Pi}]_{\text{SR,junc}} - [\text{Ca}^{2+}\text{-Pi}]_{\text{SR,bulk}})$ |
| $\frac{d[\text{CSQN}]_{\text{SR,bulk}}}{dt}$ | $-J_{\text{Ca}^{2+}\text{-CSQN,bulk}} + \frac{D_{\text{CSQN}} S A_{\text{SR,bulk-junc}}}{vol_{\text{SR,bulk}} \Delta d_{\text{diff}}} ([\text{CSQN}]_{\text{SR,junc}} - [\text{CSQN}]_{\text{SR,bulk}})$ |
| $\frac{d[\text{Ca}^{2+}\text{Trop}]}{dt}$ | $J_{\text{Trop-Ca}^{2+}} - J_{\text{Trop-2Ca}^{2+}}$ |
| $\frac{d[\text{Trop-2Ca}^{2+}]}{dt}$ | $J_{\text{Trop-2Ca}^{2+}} - J_D$ |
| $\frac{d[D]}{dt}$ | $J_D - J_{\text{pre}} + J_{\text{detach}}$ |
| $\frac{d[A_{\text{pre}}]}{dt}$ | $J_{\text{pre}} - J_{\text{post}}$ |
| $\frac{d[A_{\text{post}}]}{dt}$ | $J_{\text{post}} - J_{\text{detach}}$ |

**Table A6:** (continued) System of ODEs.
| Variable | Equation |
| --- | --- |
| $\frac{d[\text{ADP}]_{\text{myo,bulk}}}{dt}$ | $J_{\text{hydro,bulk}} - J_{\text{Gly,bulk}} N_{\text{ATP-per-NADH}} - J_{\text{ANT,bulk}} \frac{vol_{\text{Mito,bulk}}}{vol_{\text{myo,bulk}}} + J_{\text{detach}} + \frac{D_{\text{ATP}} S A_{\text{myo,bulk-junc}}}{vol_{\text{myo,bulk}} \Delta d_{\text{diff}}} ([\text{ADP}]_{\text{myo,junc}} - [\text{ADP}]_{\text{myo,bulk}})$ |
| $\frac{d[\text{Ca}^{2+}]_{\text{Mito,bulk}}}{dt}$ | $b_{\text{M}} (J_{\text{MCU,bulk}} - J_{\text{NCX,Mito,bulk}} + J_{\text{mPTP,bulk}} - J_{\text{Ca}^{2+}\text{-phos,Mito,bulk}})$ |
| $\frac{d[\text{Pi}]_{\text{Mito,bulk}}}{dt}$ | $J_{\text{PiC,bulk}} - J_{\text{Ca}^{2+}\text{-phos,Mito,bulk}}$ |
| $\frac{d[\text{Ca}^{2+}\text{-Pi}]_{\text{Mito,bulk}}}{dt}$ | $J_{\text{Ca}^{2+}\text{-phos,Mito,bulk}}$ |
| $\frac{d[\text{ATP}]_{\text{Mito,bulk}}}{dt}$ | $J_{\text{F1FO,bulk}} - J_{\text{ANT,bulk}}$ |
| $\frac{d[\text{NADH}]_{\text{Mito,bulk}}}{dt}$ | $J_{\text{PDH,bulk}} + J_{\text{AGC,bulk}} - J_{\text{Ox,bulk}}$ |
| $\frac{dV_{\text{Mito,bulk}}}{dt}$ | $(a_1 J_{\text{Ox,bulk}} - a_2 J_{\text{F1FO,bulk}} - J_{\text{Hleak,Mito,bulk}} - J_{\text{NCX,Mito,bulk}} - 2J_{\text{MCU,bulk}} - 2J_{\text{mPTP,bulk}} - J_{\text{ANT,bulk}} - J_{\text{AGC,bulk}}) / C_{\text{Eff,Mito}}$ |

**Table A7:**
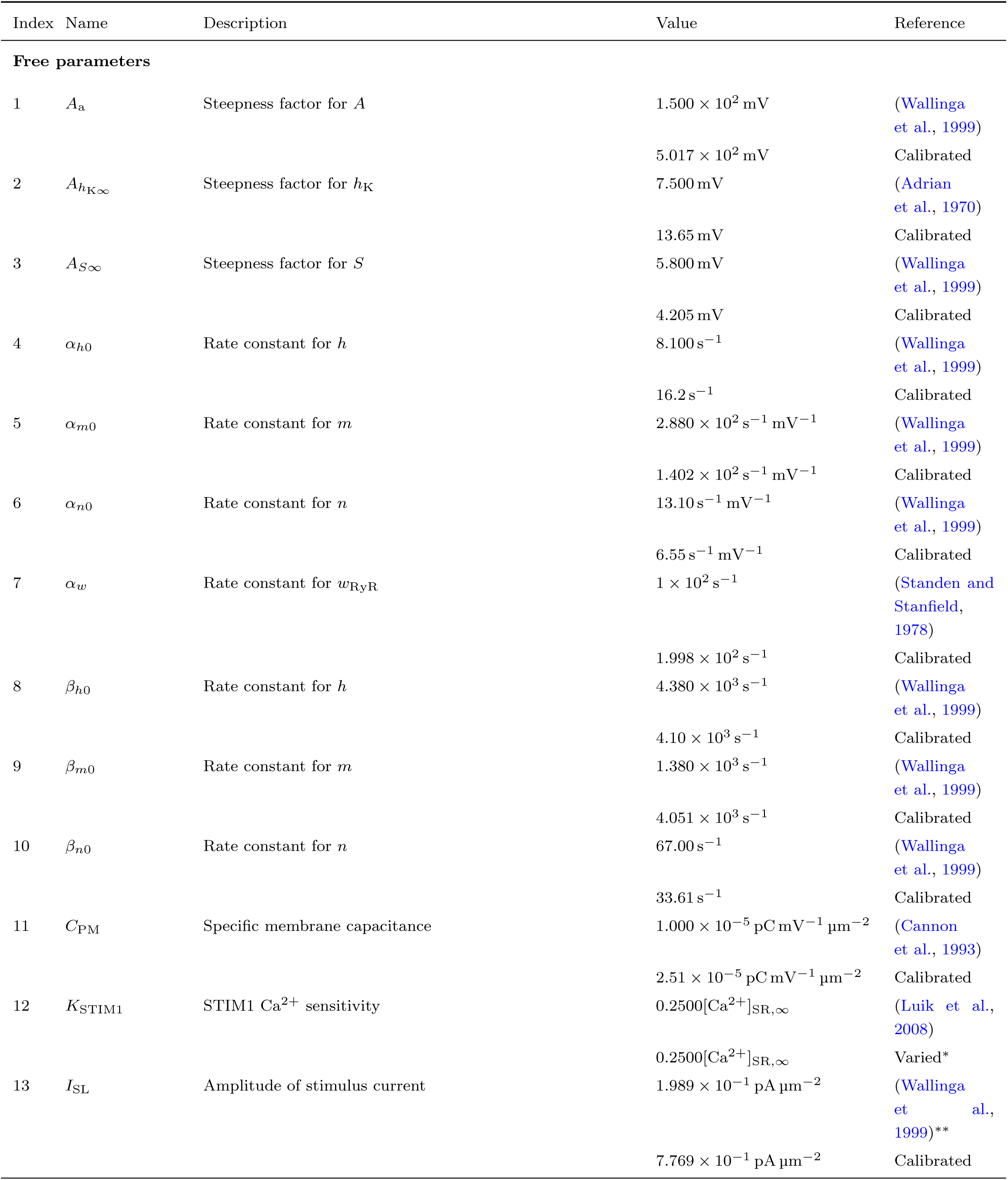
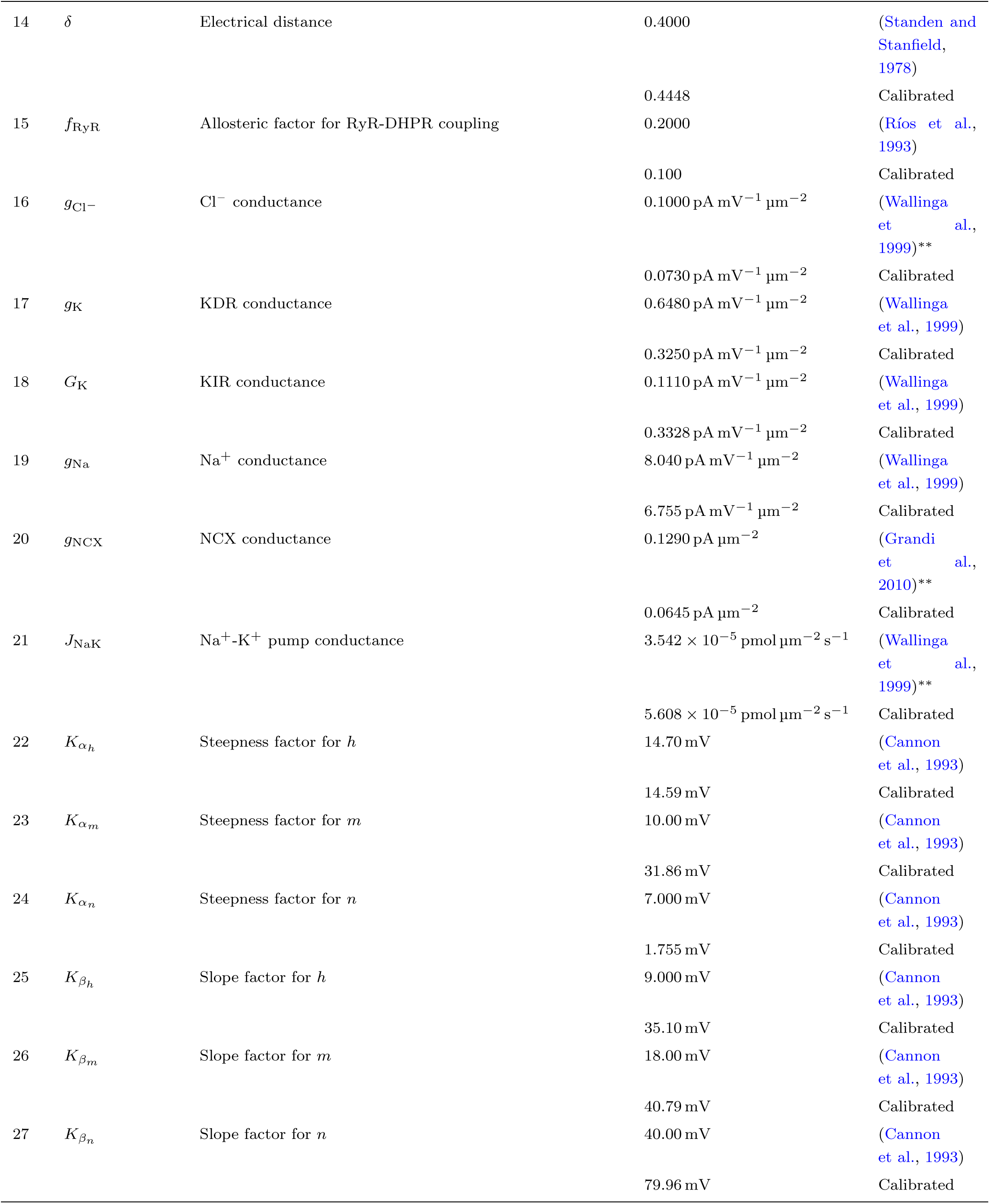
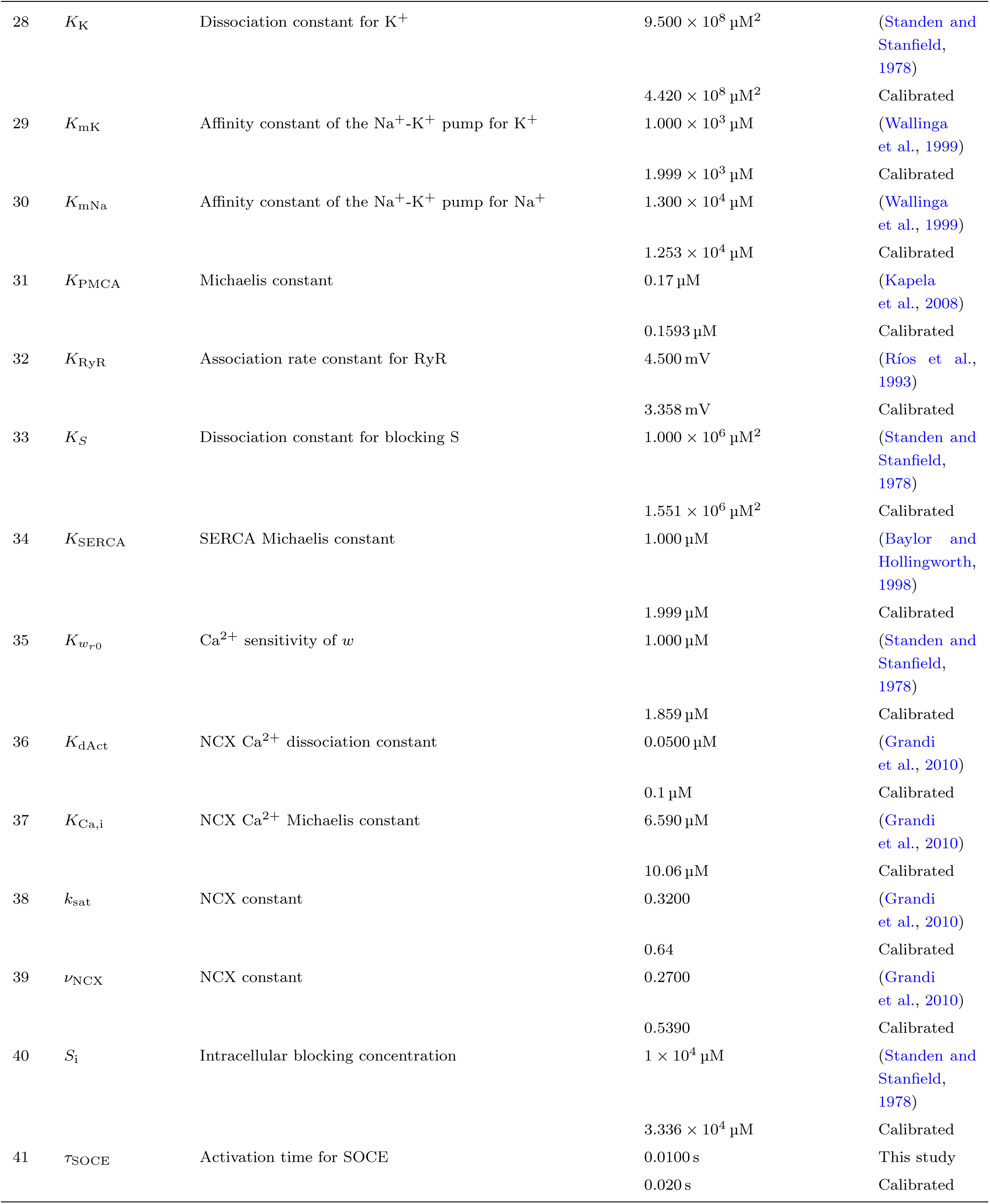
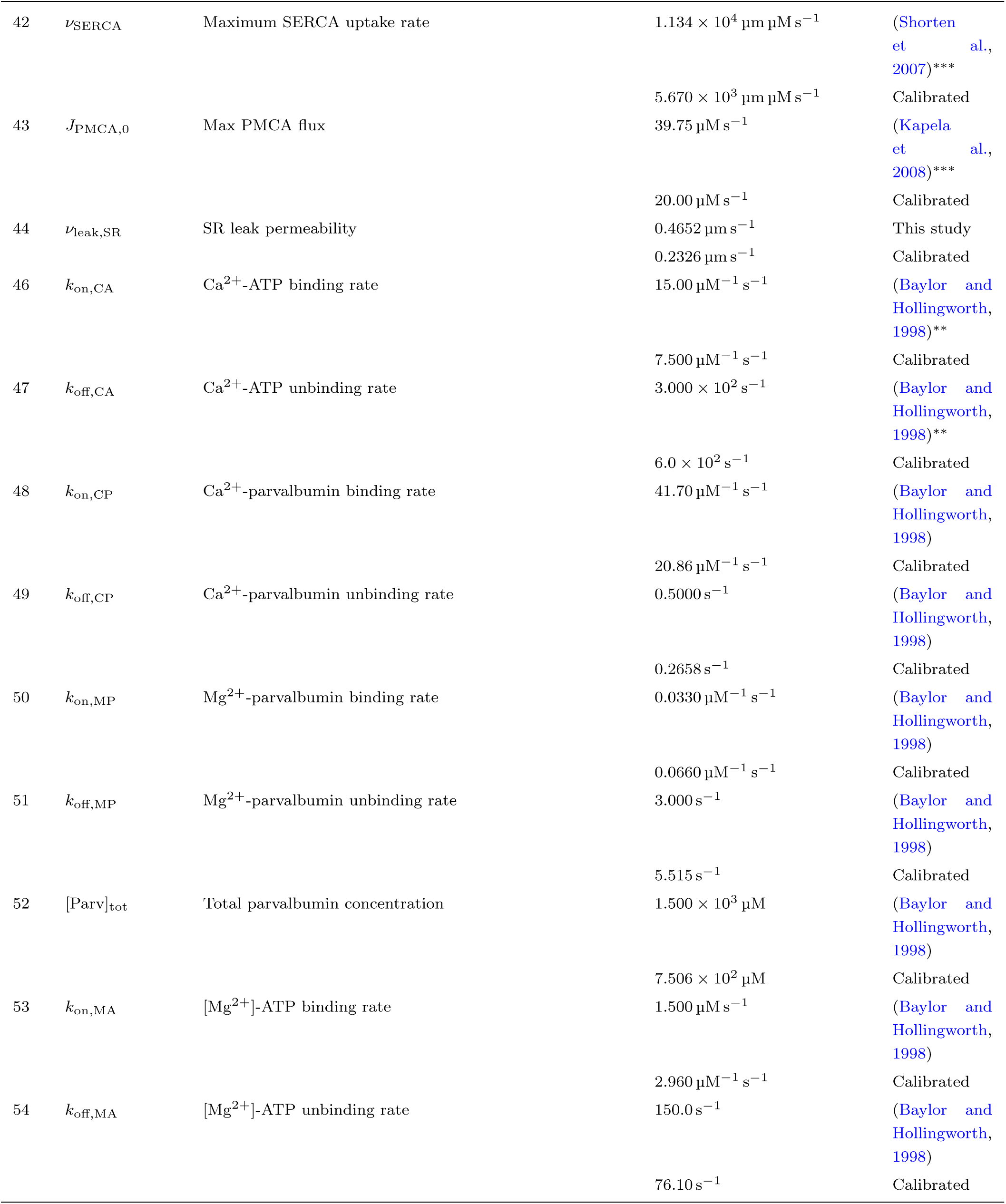
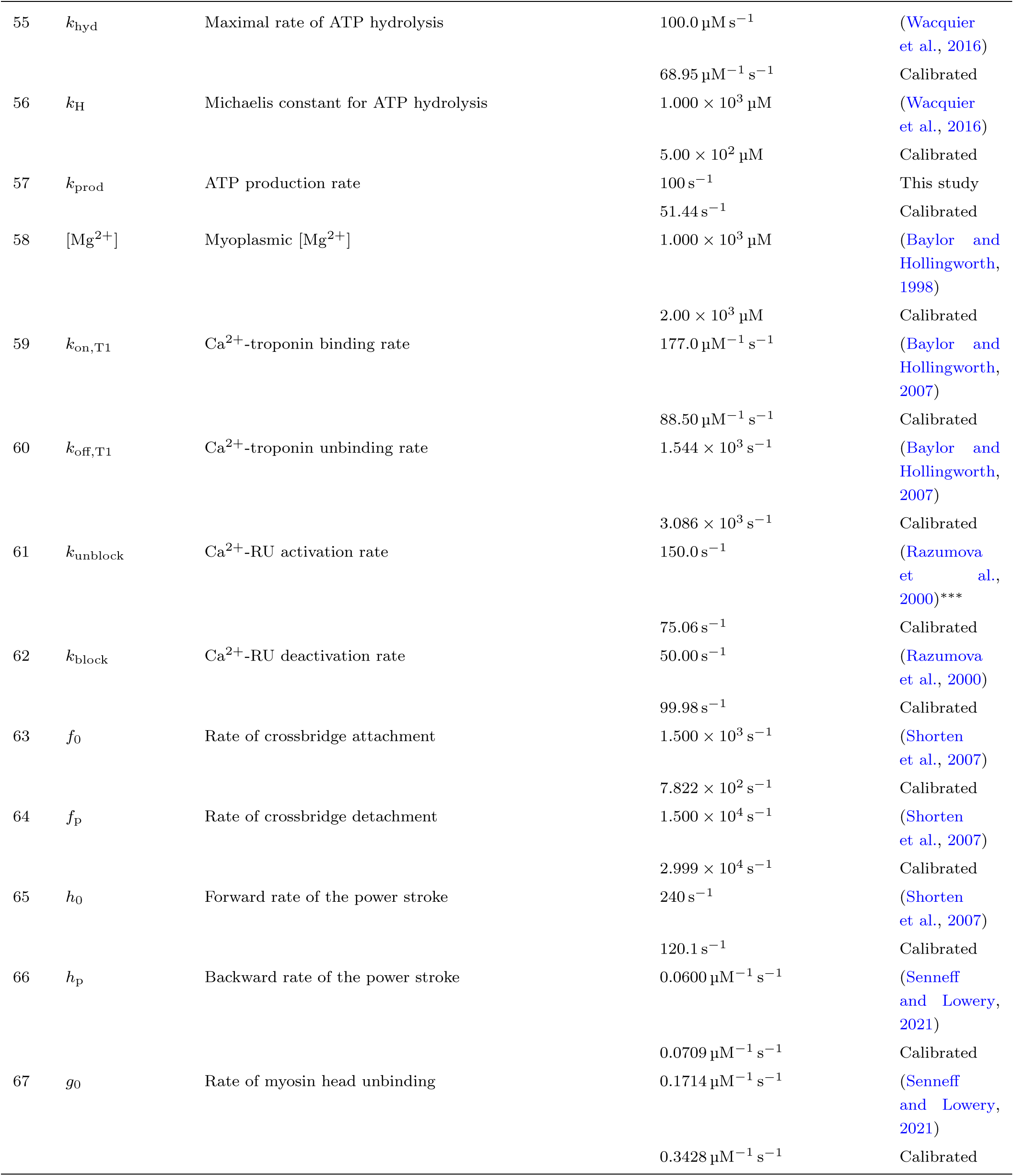
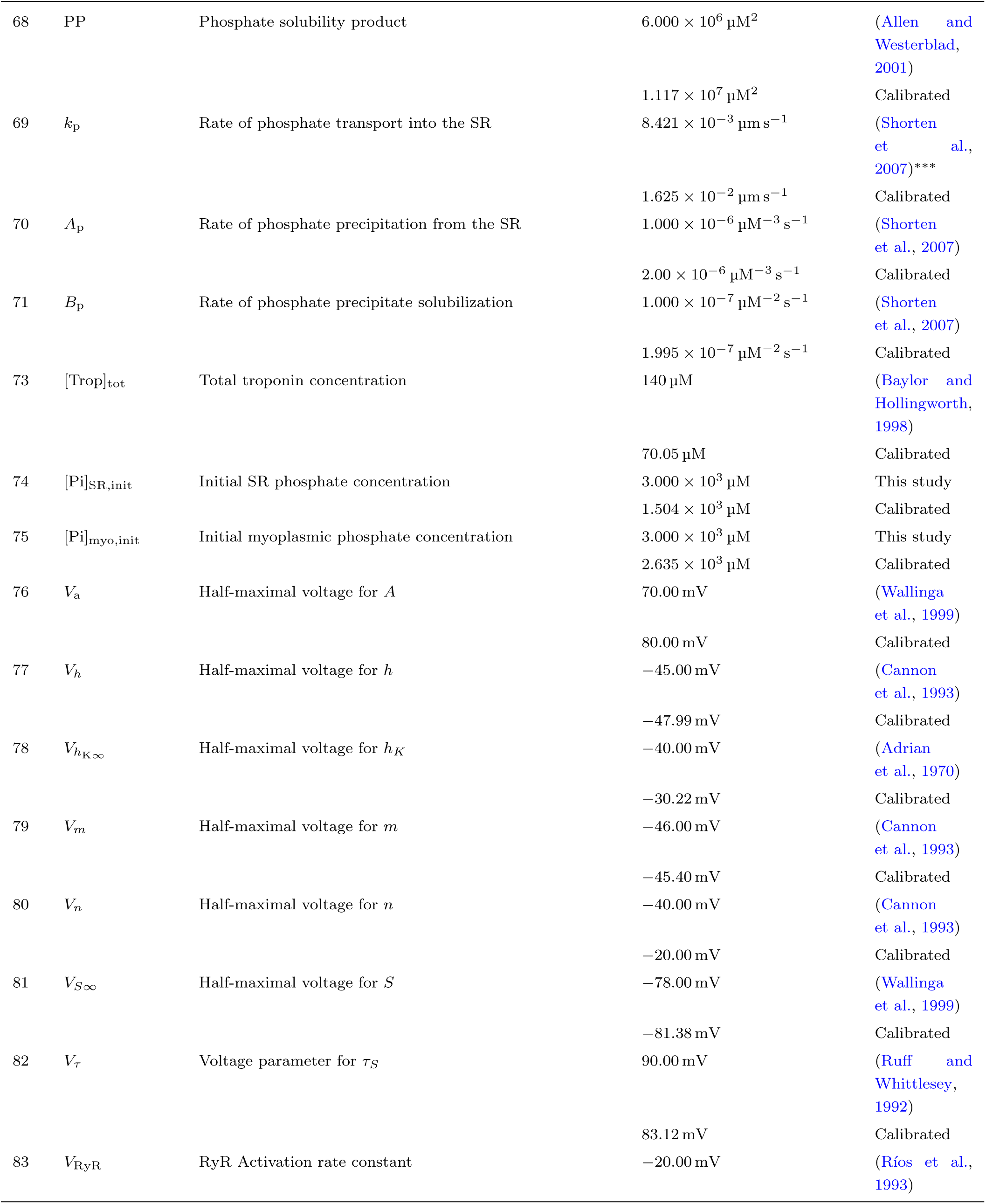
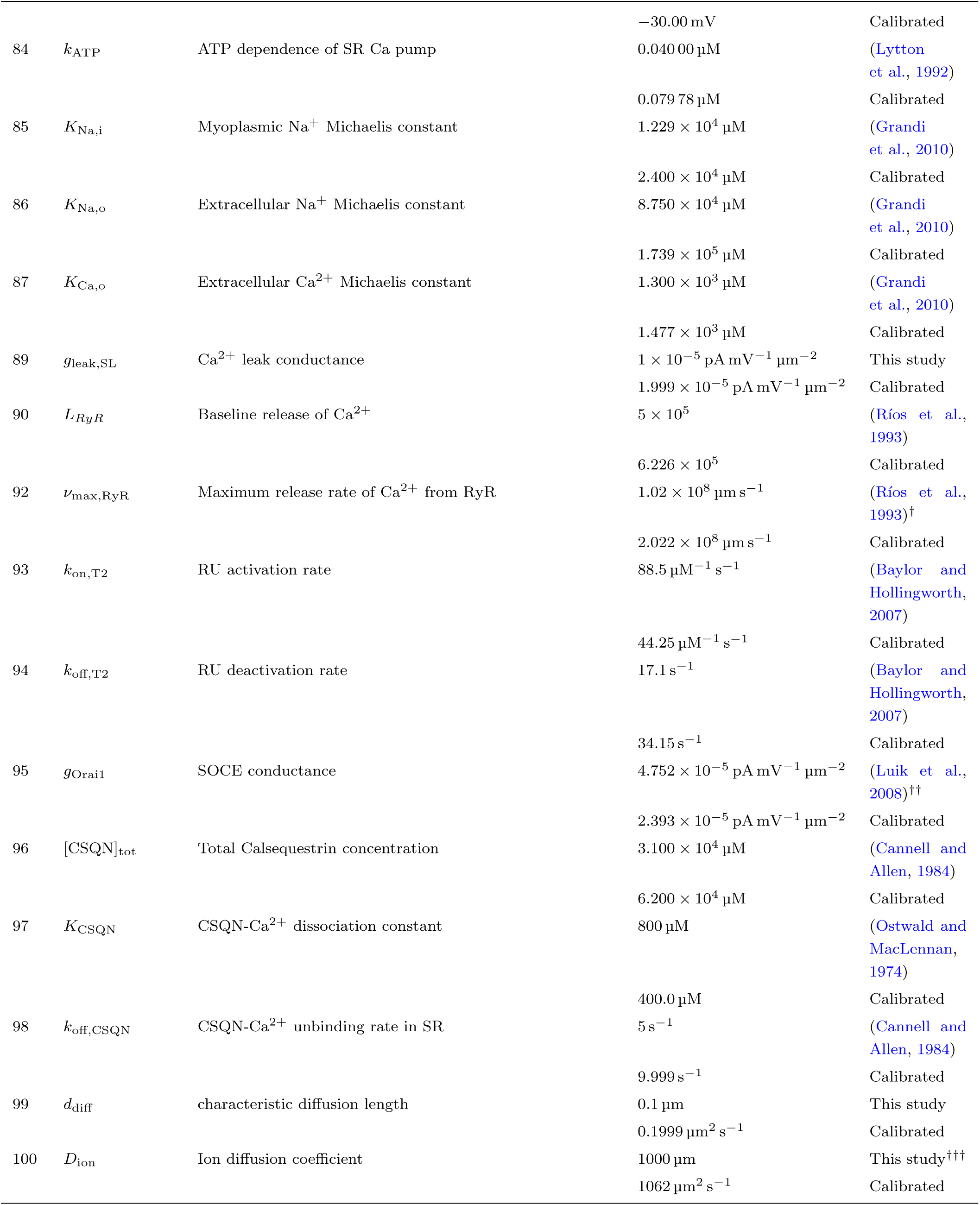
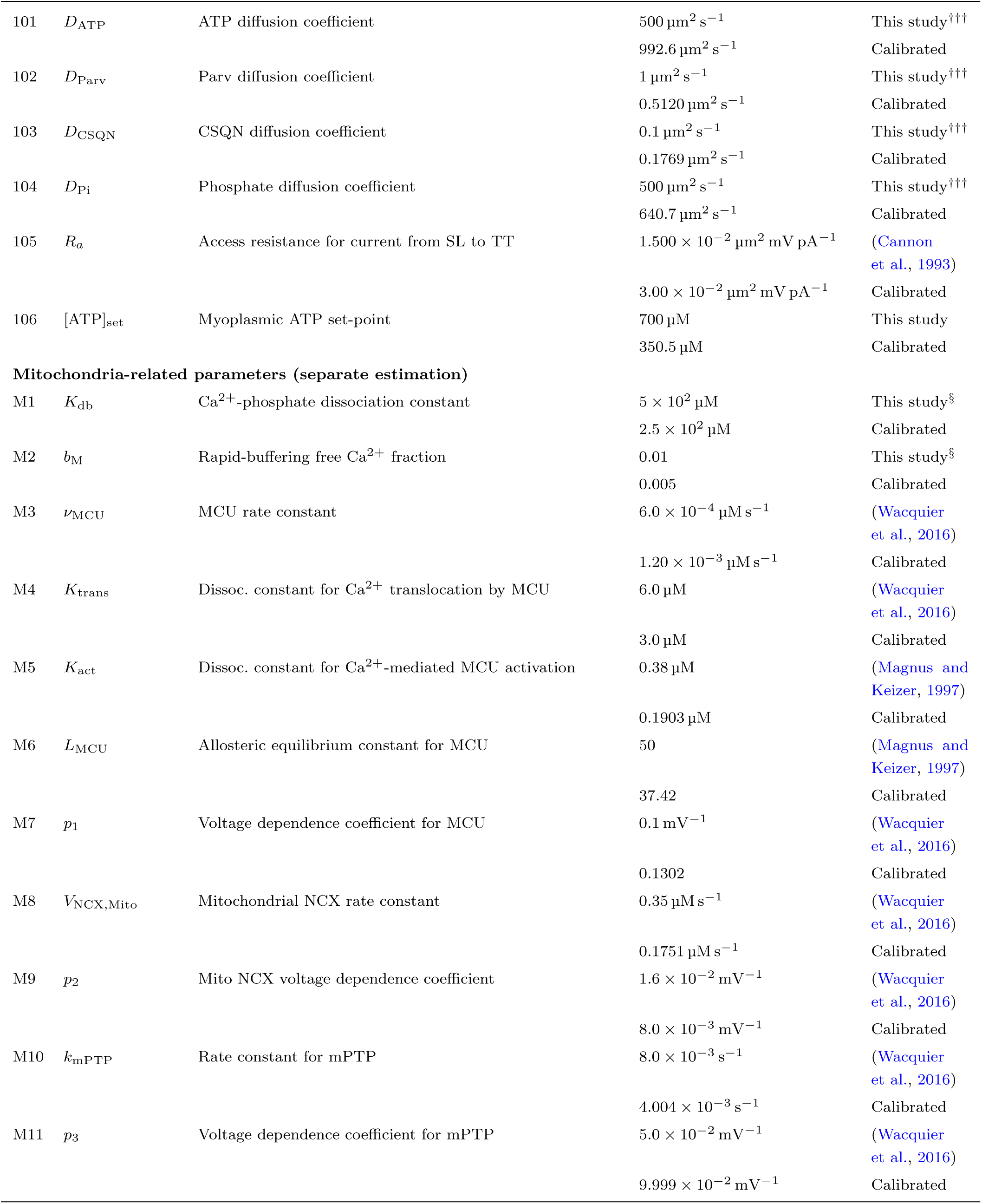
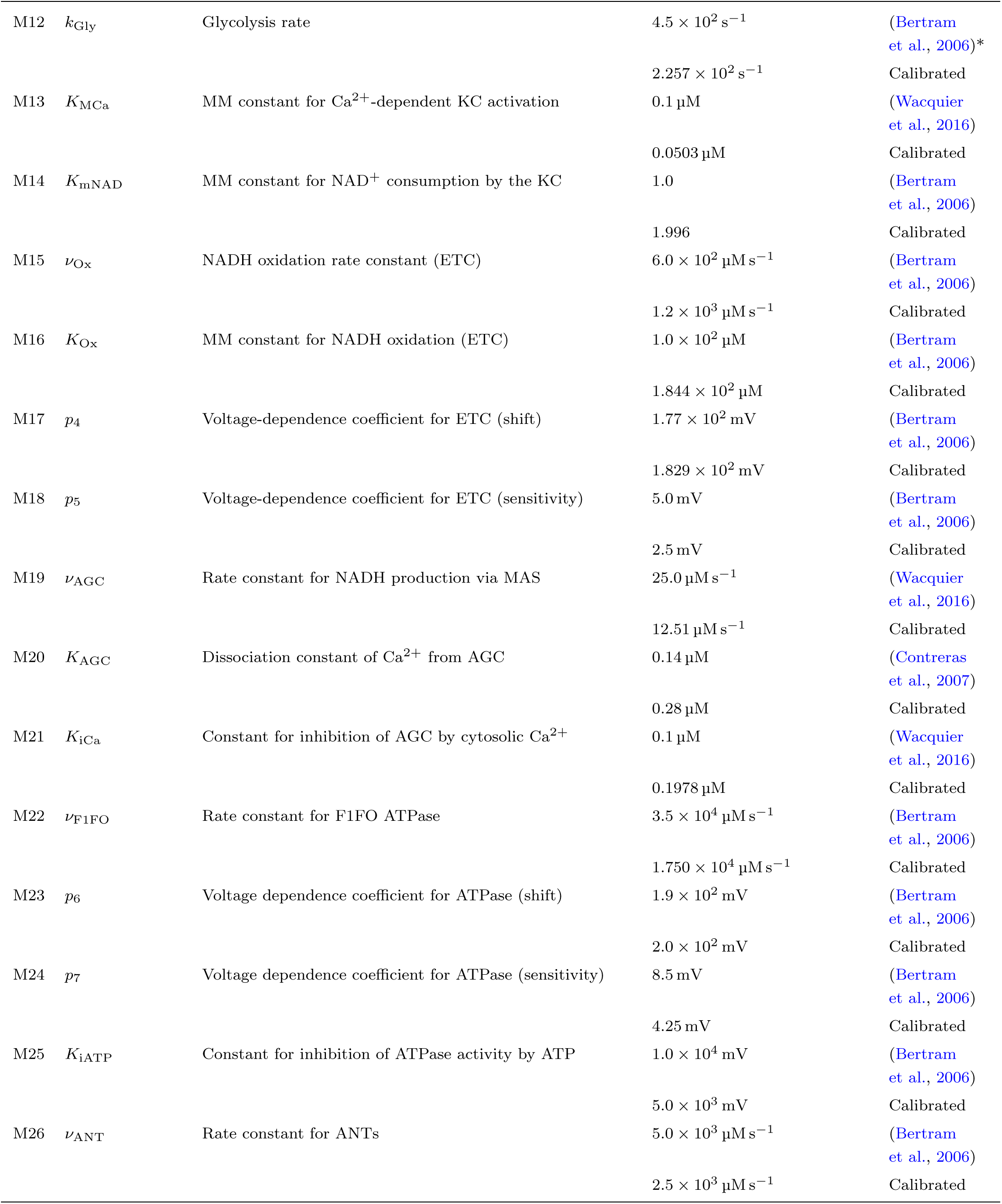
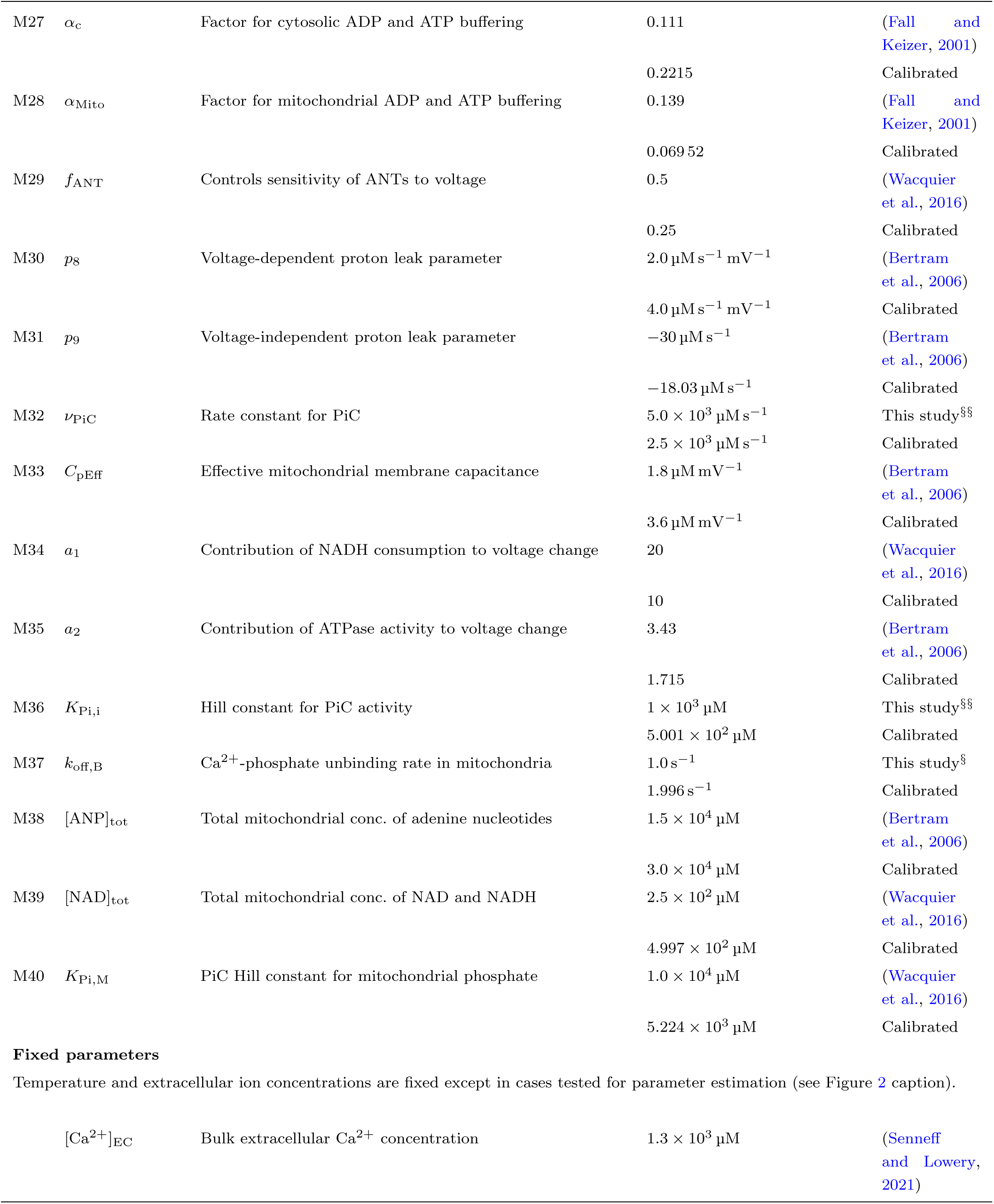
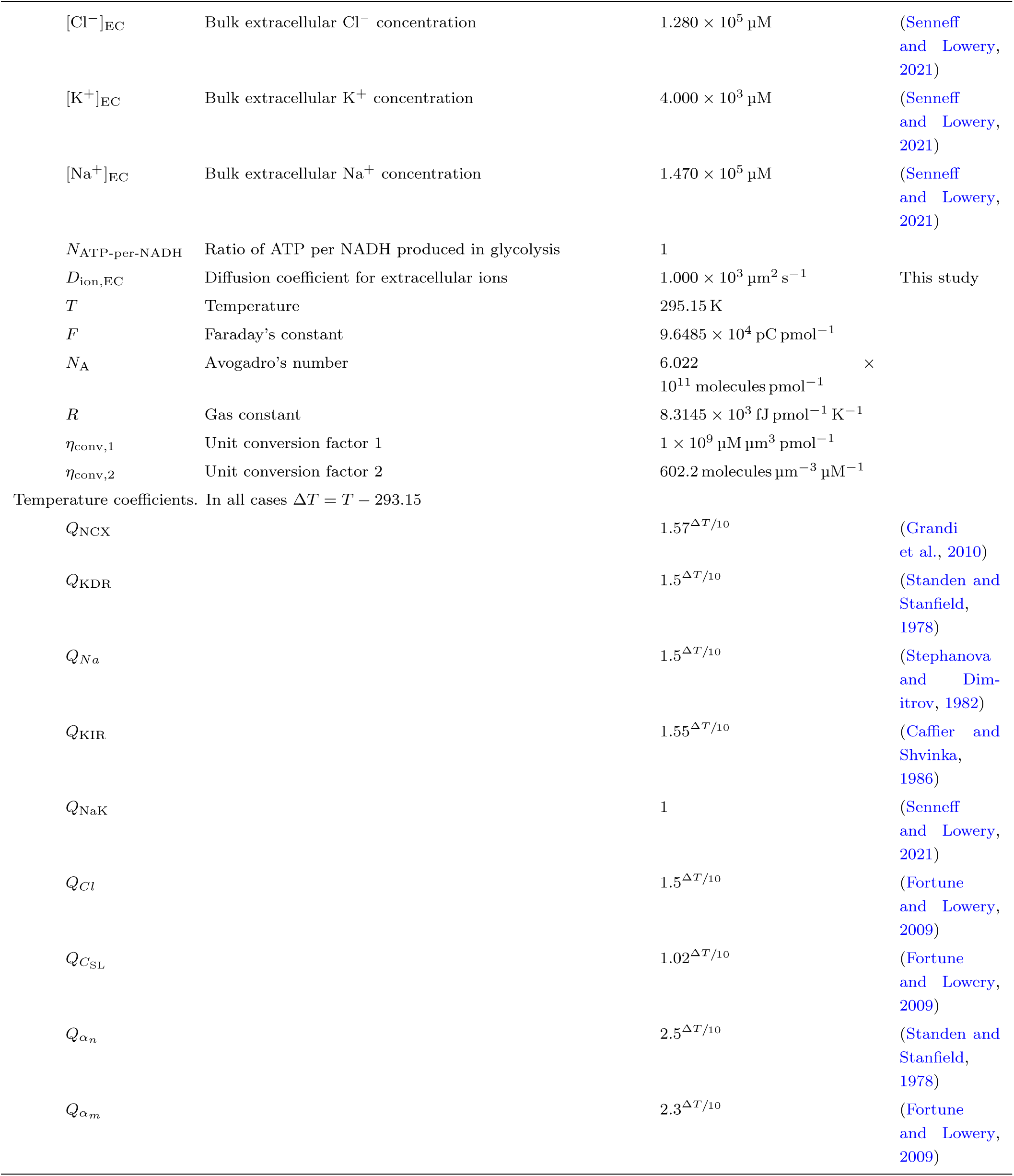
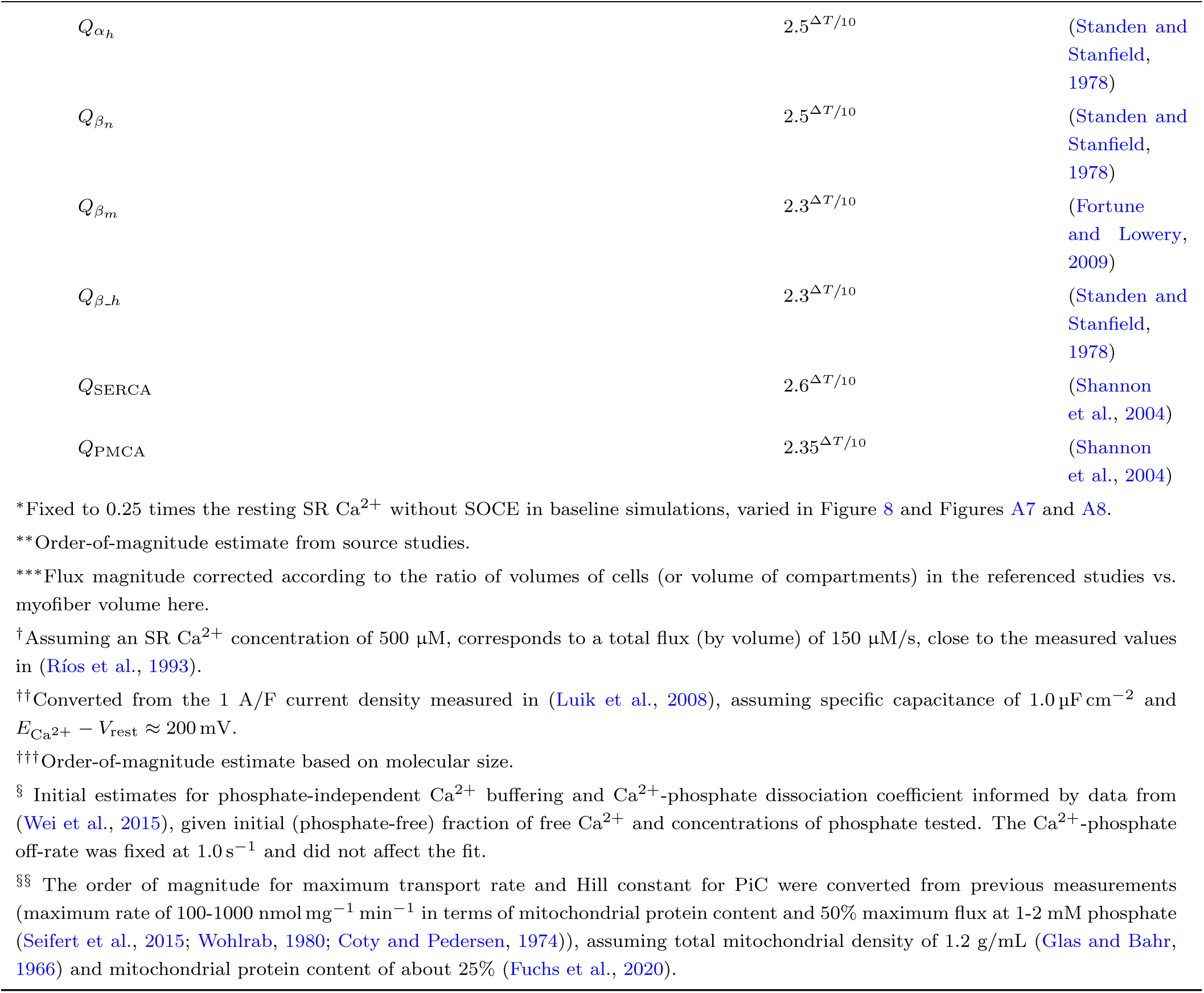
Model parameters. Acronyms used here and not previously defined – AGC: aspartate/glutamate carrier, MM: Michaelis-Menten, KC: Krebs cycle, ETC: electron transport chain, MAS: malate-aspartate shuttle.

## Appendix B Supplementary figures

**Figure A1:**
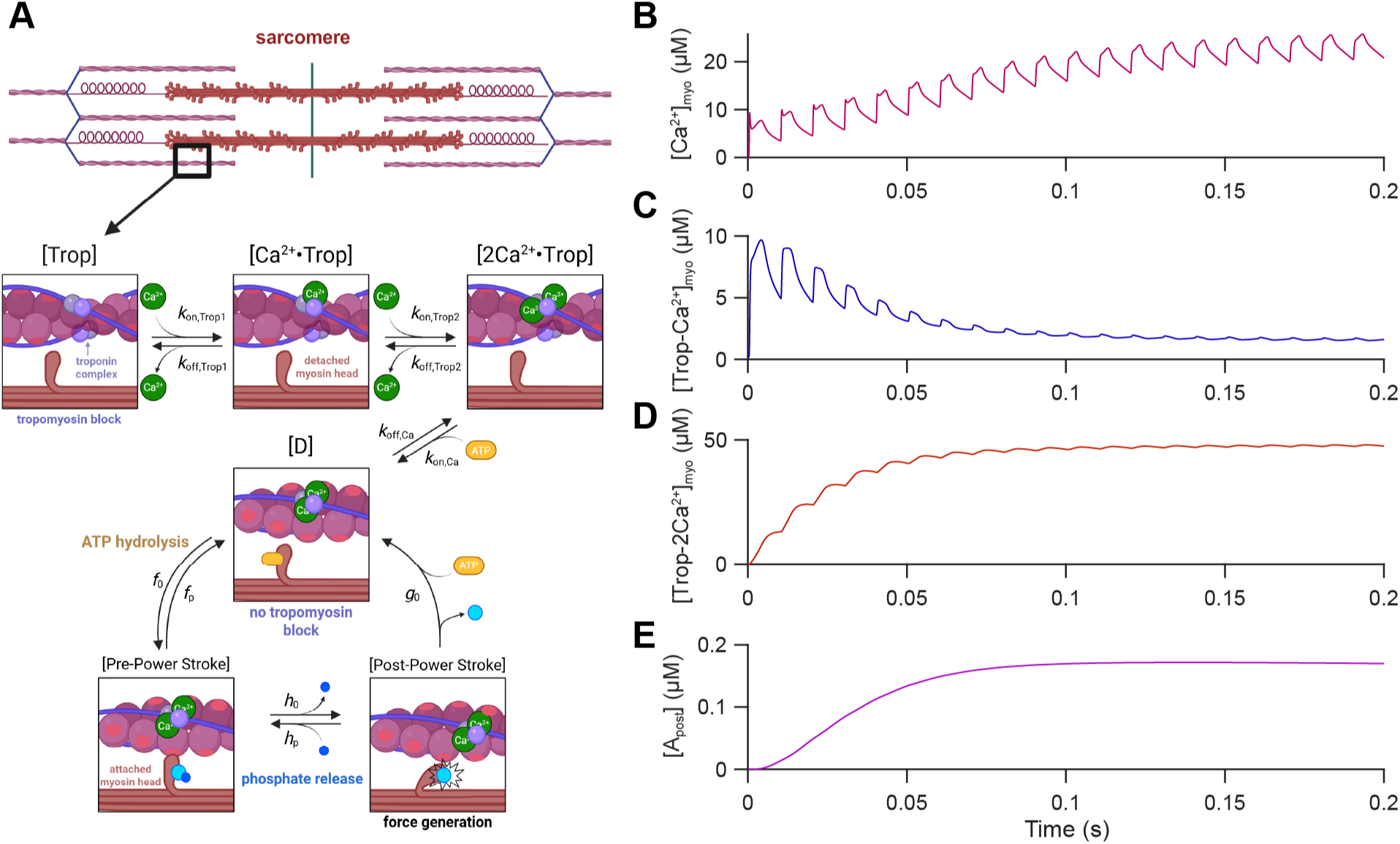
Summary of six-state cross-bridge model. A) Schematic of cross-bridge model. Two calcium ions bind to troponin, facilitating unblocking of the myosin-head binding sites on actin. ATP hydrolysis enables myosin-head binding to actin, and phosphate release drives the power stroke. Figure created in BioRender. B-E) Model predictions of myoplasmic Ca^2+^ (B) and different cross-bridge states during repetitive stimuli, including troponin bound to one Ca^2+^ (C), troponin bound to two Ca^2+^ (D), and cross-bridges post-power stroke (E).

**Figure A2:**
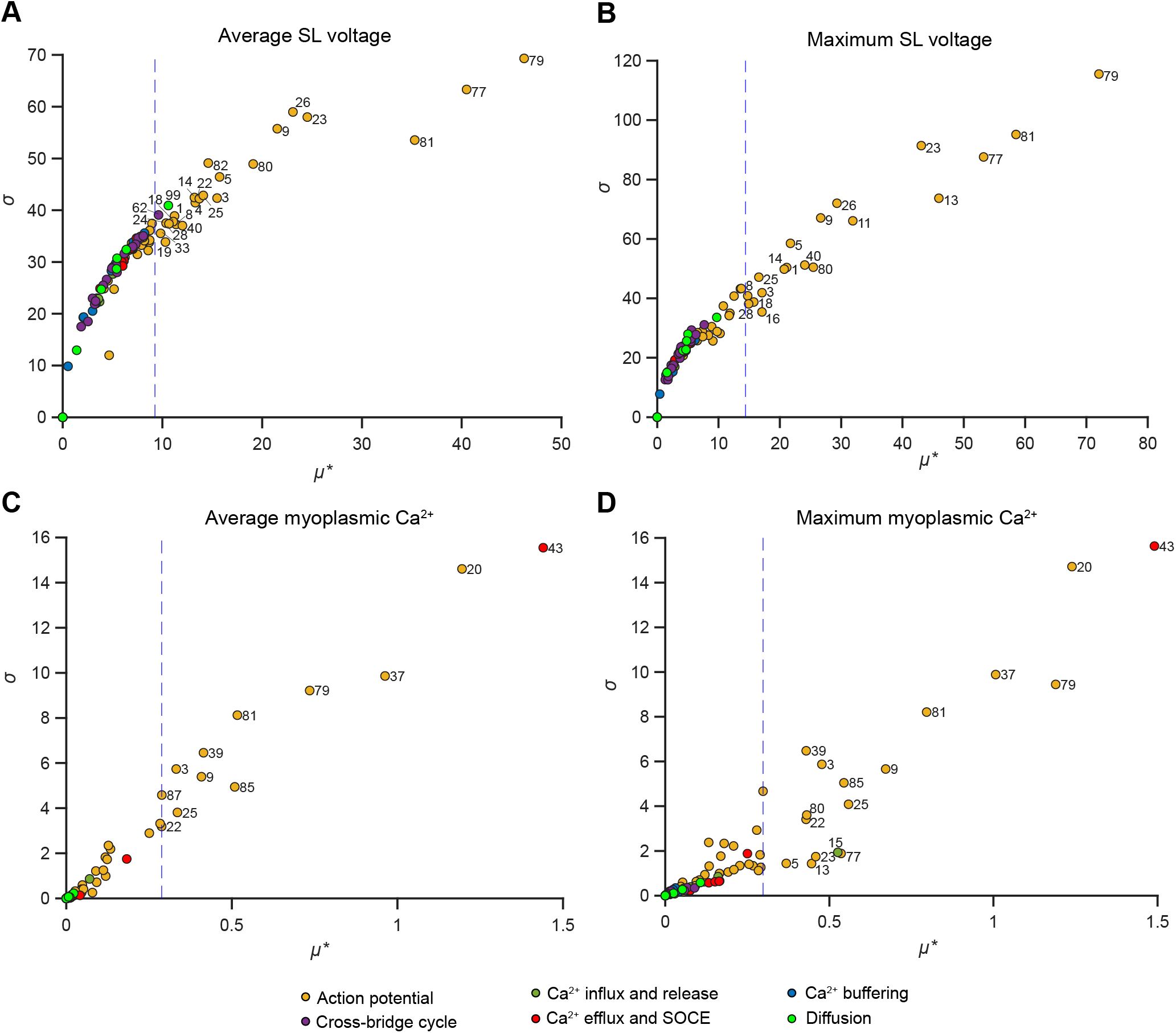
Morris sensitivity analysis for four main quantities of interest. Model sensivity was assessed with respect to average sarcolemma voltage (A), peak sarcolemma voltage (B), average myoplasmic Ca^2+^ (C), and peak myoplasmic Ca^2+^ (D). In plots associated with each QOI, the horizontal axis displays the absolute value of the mean, *µ^∗^* and the vertical axis shows the standard deviation, *σ*, of elementary effects. All parameters with *µ^∗^* greater than 20% of the maximum value are labeled according to their assigned number in Table A7.

**Figure A3:**
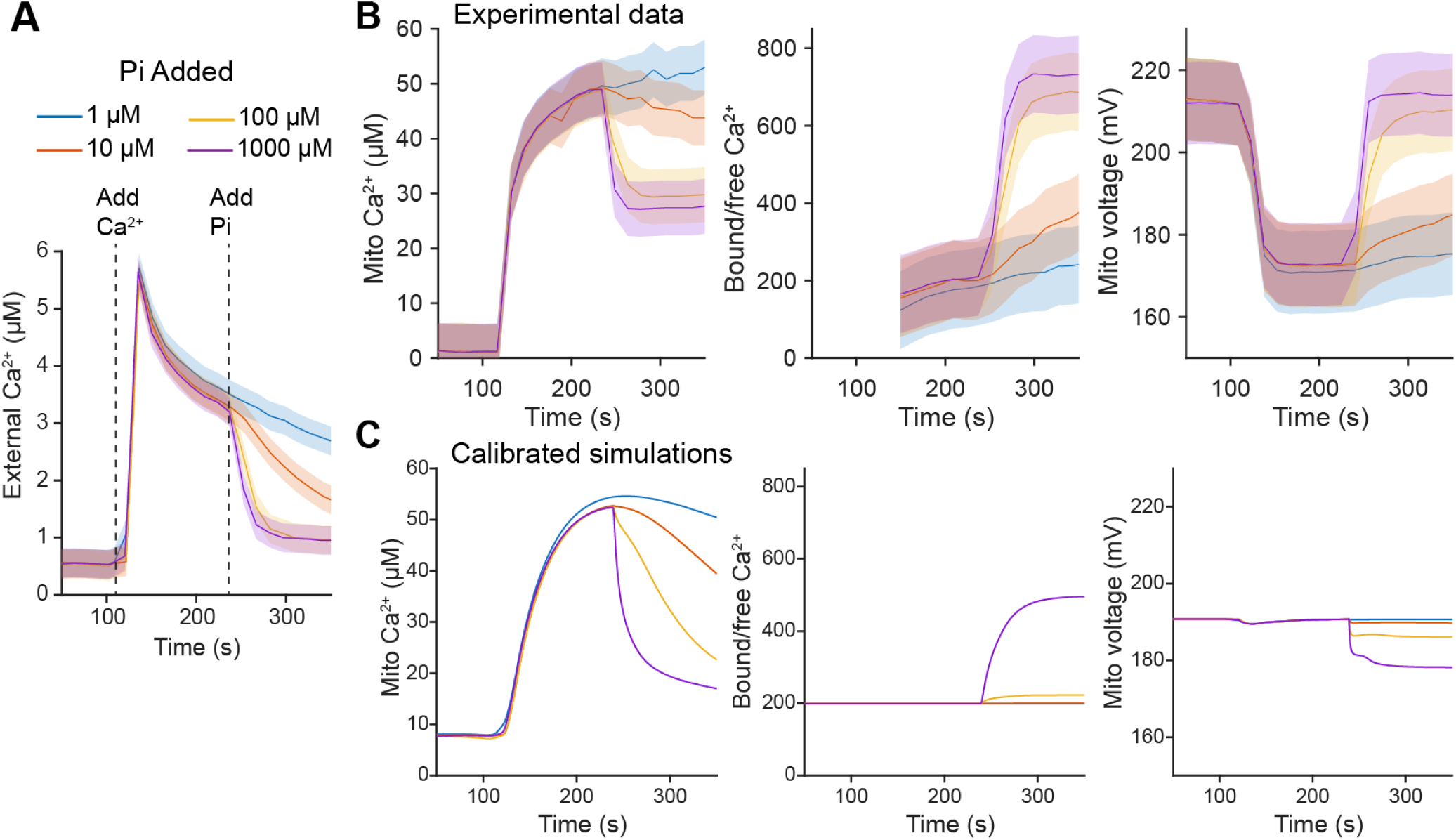
Calibration of mitochondrial Ca^2+^ and phosphate handling for Module 5. A) Experimental measurements of Ca^2+^ in the extra-mitochondrial solution from (Wei et al., 2015). The mean curve is used as a driving function in the simulations shown in panel C. B) Experimental measurements from (Wei et al., 2015) (*n* = 3 experiments in each case). Error bars in A and B indicate the assumed standard error of the mean in our calibration procedure (see Equation (17)). C) Predictions of our model with calibrated parameters as reported in Table A7, assuming time-dependent Ca^2+^ from panel A and time-dependent phosphate as a Heaviside function with the indicated concentration, starting at time *t* = 240 s. The minimized weighted sum of square errors (defined in Equation (17)) was 38.518.

**Figure A4:**
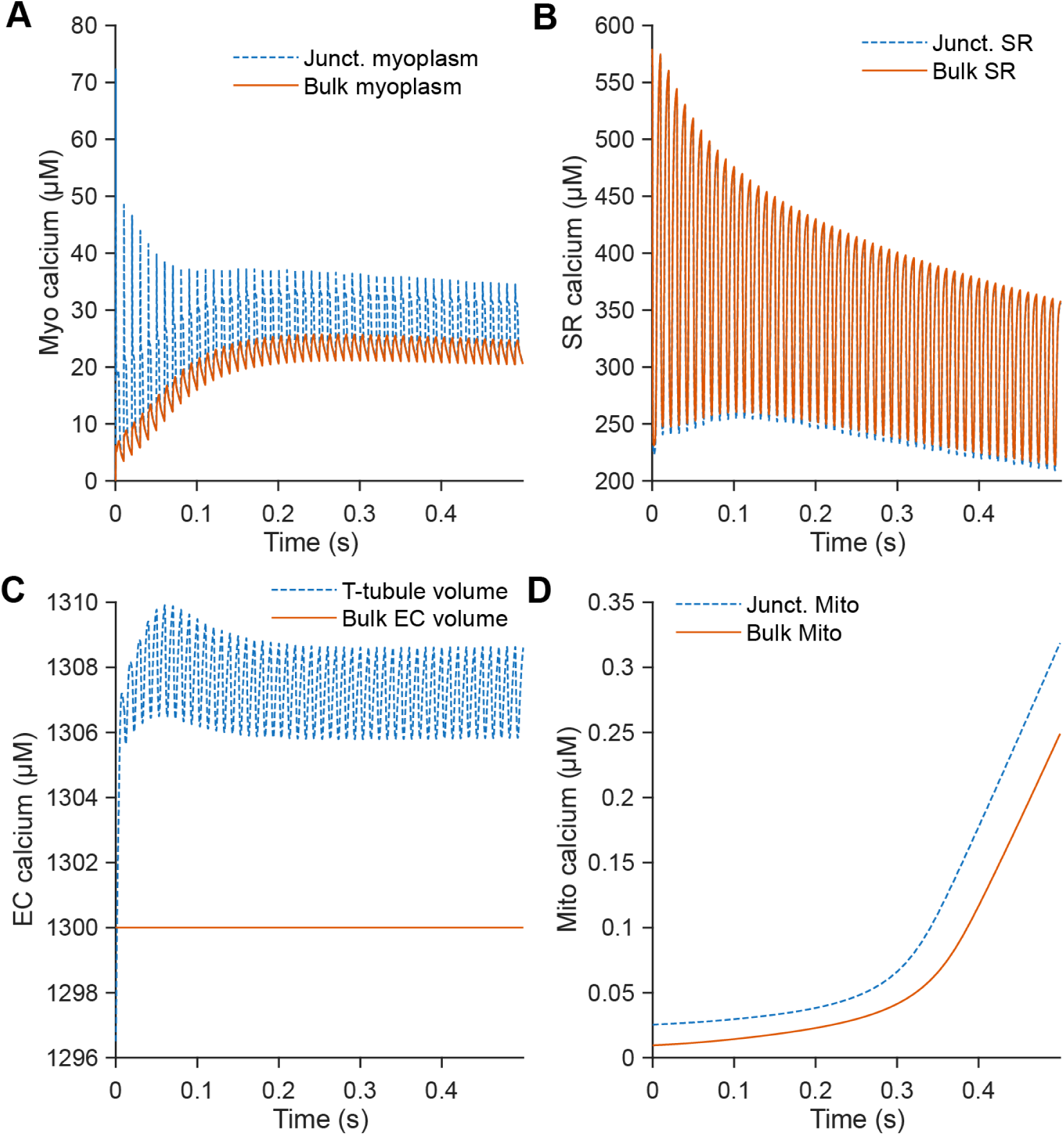
Calcium concentrations in different subregions of the compartmental model. A) Schematic summarizing the subregions of extracellular, myoplasmic, and SR compartments. B) Myoplasmic Ca^2+^ in the bulk vs. junctional myoplasm. B) SR Ca^2+^ in the bulk vs. junctional SR. C) Ca^2+^ in the bulk extracellular volume vs. the T-tubule volume. C) Ca^2+^ in bulk mitochondria vs. junctional mitochondria. All concentrations were computed from the same test conditions shown in Figure 3.

**Figure A5:**
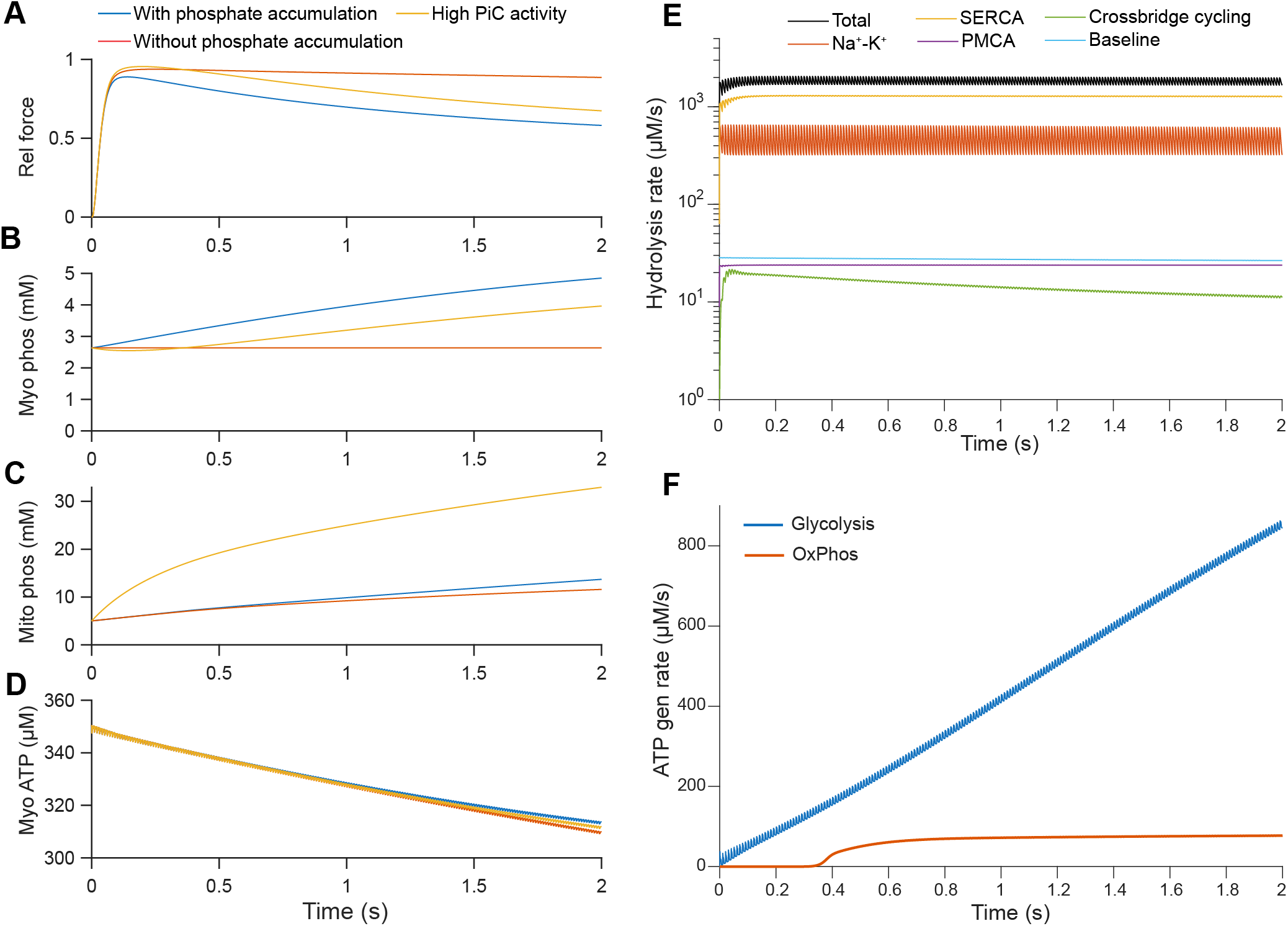
Effects of phosphate accumulation due to ATP hydrolysis in the myoplasm. A-D) Model predictions with and without phosphate accumulation, or with 10 times the baseline value for PiC permeability. Plots show relative force (A), myoplasmic phosphate (B), mitochondrial phosphate (C), and myoplasmic ATP (D). E) Contributions of each ATP-consuming process in the model for a simulation with normal phosphate accumulation. F) Contribution of glycolysis vs. oxidative phosphorylation to ATP generation over time for a simulation with normal phosphate accumulation.

**Figure A6:**
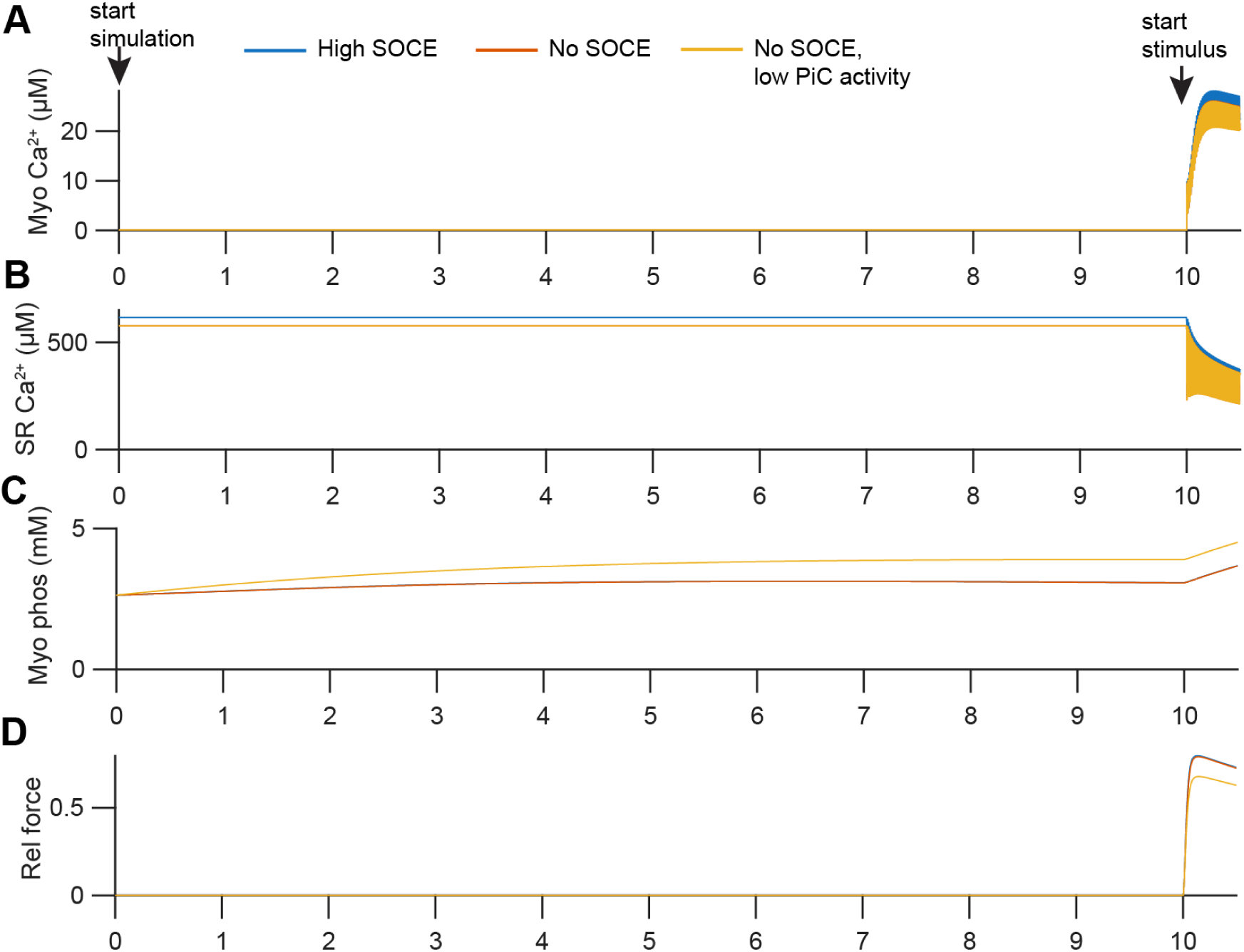
Initialization protocol for simulations shown in Figure 7. After normal initialization of the system (as described in Methods), these tests were run for 10 s before the stimulus occurred. As shown, myoplasmic (A) and SR (B) Ca^2+^ concentrations remain mainly constant prior to stimulus, whereas myoplasmic phosphate gradually accumulates due to resting ATP hydrolysis (C). The resulting predictions for relative force are shown in panel D.

**Figure A7:**
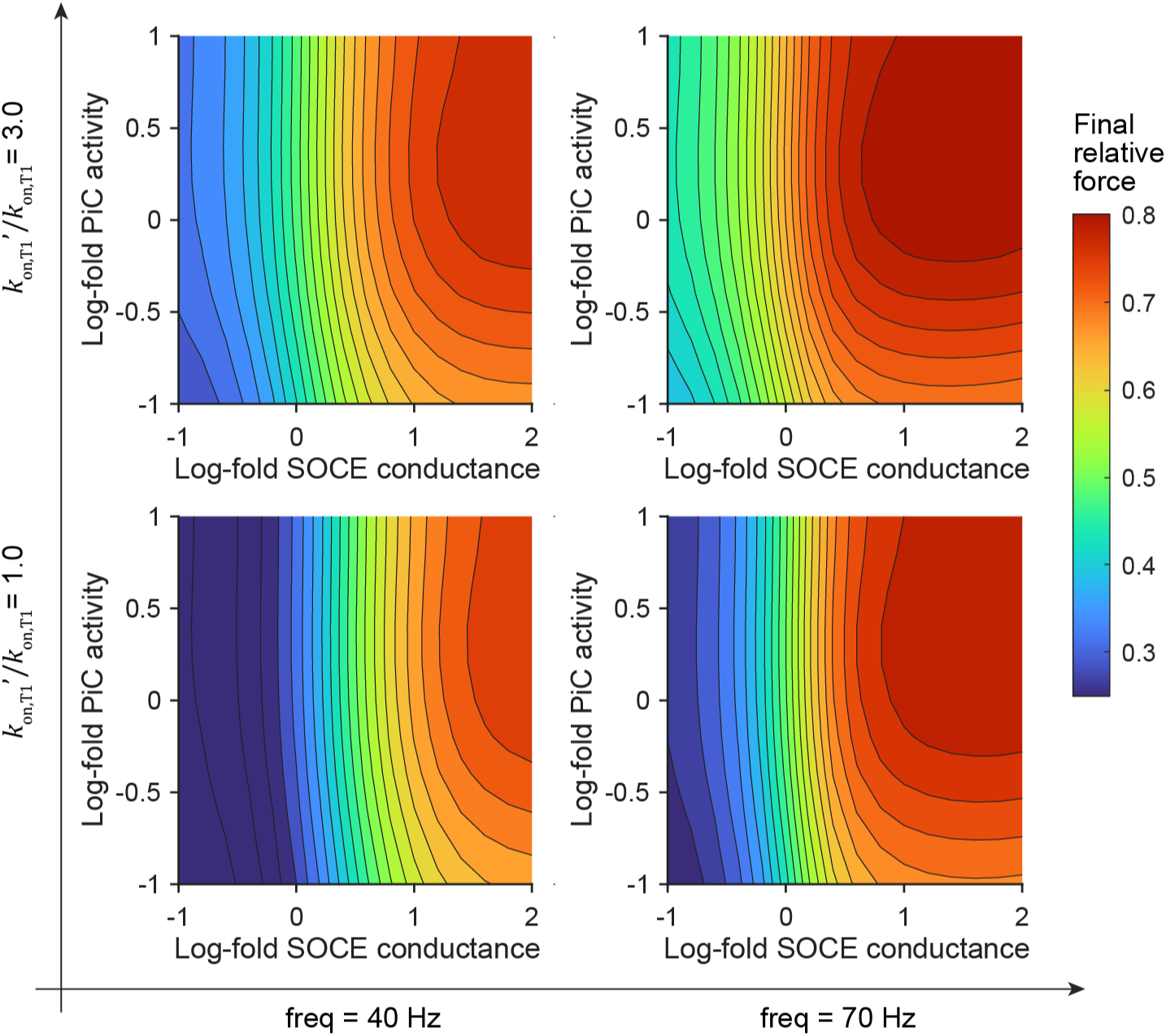
Effect of SOCE on force during resistance exercise, tested over different parameter regimes. Force as a function of Orai1 conductance and PiC activity in the final repetition of a 60 s resistance exercise simulation for stimulus frequencies of 40 Hz (left) or 70 Hz (right) and for troponin Ca^2+^ sensitivity at 100% (lower) or 300% (upper) of the estimated value.

**Figure A8:**
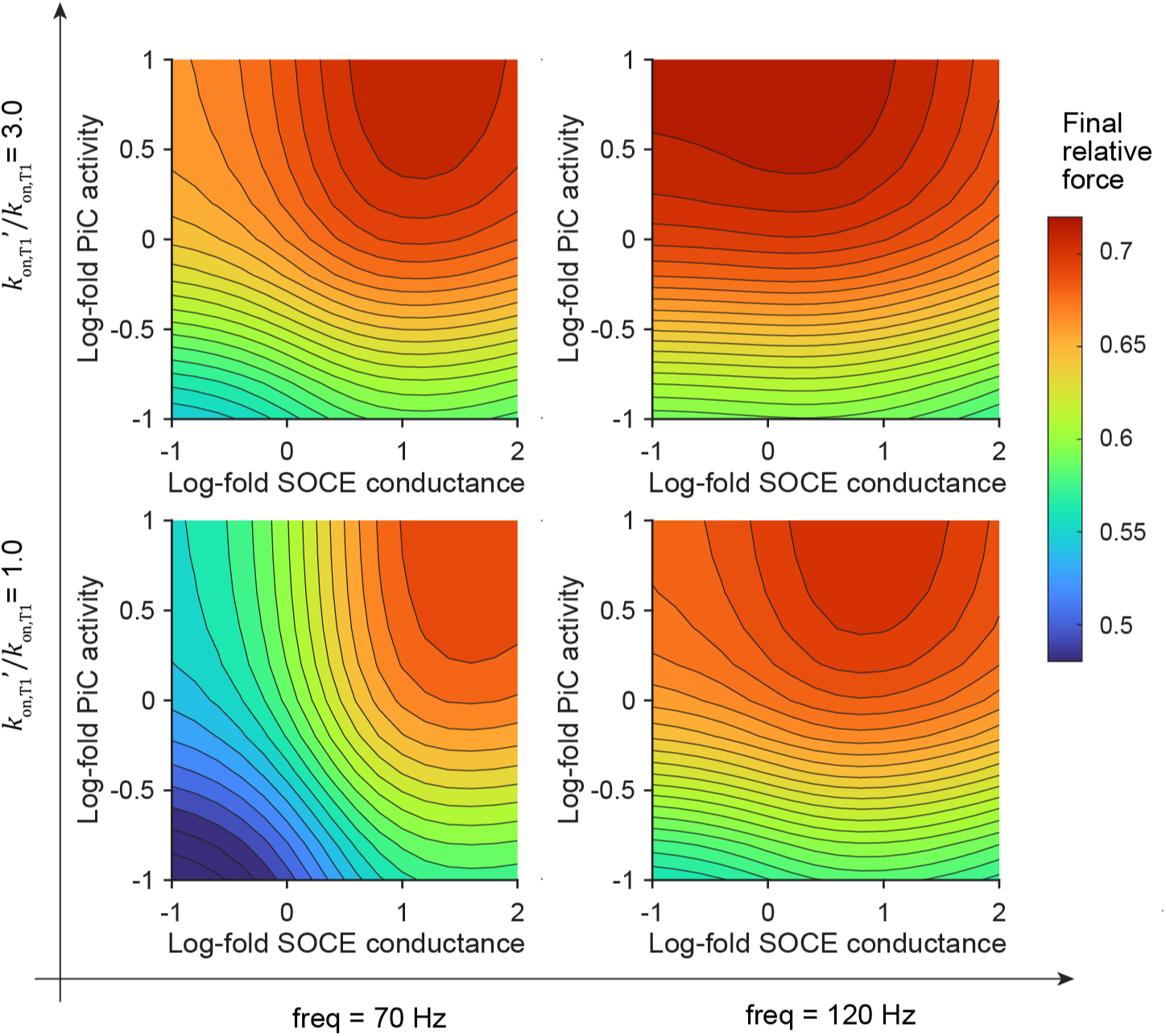
Effect of SOCE on force during high-intensity interval training (HIIT), tested over different parameter regimes. Force as a function of Orai1 conductance and PiC activity in the final stride of a 20 s HIIT simulation for stimulus frequencies of 70 Hz (left) or 120 Hz (right) and for troponin Ca^2+^ sensitivity at 100% (lower) or 300% (upper) of the estimated value.

## Notes

https://doi.org/10.5281/zenodo.15485446

### Competing Interest Statement

Padmini Rangamani is a consultant for Simula Research Laboratory in Oslo, Norway and receives income. The terms of this arrangement have been reviewed and approved by the University of California, San Diego in accordance with its conflict-of-interest policies.

### Summary of Updates

Our model has been significantly revised to include the effects of mitochondrial calcium and phosphate handling. These model additions revealed a crucial role for mitochondrial phosphate uptake in mitigating fatigue in certain cases of high store-operated calcium entry during extended exercise. All figures were updated to align with the new model and the text was updated throughout.

